# Multi-omics integration analysis identifies novel genes for alcoholism with potential link to neurodegenerative diseases

**DOI:** 10.1101/2020.10.15.341750

**Authors:** Manav Kapoor, Michael Chao, Emma C. Johnson, Gloriia Novikova, Dongbing Lai, Jacquelyn Meyers, Jessica Schulman, John I Nurnberger, Bernice Porjesz, Yunlong Liu, COGA collaborators, Tatiana Foroud, Howard J. Edenberg, Edoardo Marcora, Arpana Agrawal, Alison Goate

## Abstract

**Significance:** Identification of causal variants and genes underlying genome-wide association study (GWAS) loci is essential to understanding the biology of alcohol use disorder (AUD).

**Methods:** Integration of “multi-omics” data is often necessary to nominate candidate causal variants and genes and prioritize them for follow up studies. Here, we used Mendelian randomization to integrate AUD and drinks per week (DPW) GWAS summary statistics with the gene expression and methylation quantitative trait loci (eQTLs and mQTLs) in the largest brain and myeloid datasets. We also used AUD-related single cell epigenetic data to nominate candidate causal variants and genes associated with DPW and AUD.

**Results:** Our multi-omics integration analyses prioritized unique as well as shared genes and pathways among AUD and DPW. The GWAS variants associated with both AUD and DPW showed significant enrichment in the promoter regions of fetal and adult brains. The integration of GWAS SNPs with mQTLs from fetal brain prioritized variants on chromosome 11 in both AUD and DPW GWASs. The co-localized variants were found to be overlapping with promoter marks for *SPI1,* specifically in human microglia, the myeloid cells of the brain. The co-localized SNPs were also strongly associated with *SPI1* mRNA expression in myeloid cells from peripheral blood. The prioritized variant at this locus is predicted to alter the binding site for a transcription factor, RXRA, a key player in the regulation of myeloid cell function. Our analysis also identified *MAPT* as a candidate causal gene specifically associated with DPW. mRNA expression of *MAPT* was also correlated with daily amounts of alcohol intake in post-mortem brains (frontal cortex) from alcoholics and controls (N = 92). Results may be queried and visualized in an online public resource of these integrative analysis (https://lcad.shinyapps.io/alc_multiomics/). These results highlight overlap between causal genes for neurodegenerative diseases, alcohol use disorder and alcohol consumption.

**In conclusion:** integrating GWAS summary statistics with multi-omics datasets from multiple sources identified biological similarities and differences between typical alcohol intake and disordered drinking highlighting molecular heterogeneity that might inform future targeted functional and cross-species studies. Interestingly, overlap was also observed with causal genes for neurodegenerative diseases.

## Introduction

Alcohol use disorders (AUD) are complex, moderately heritable (50-60%)^1–4^, psychiatric disorders associated with heightened morbidity and mortality^5^. An AUD diagnosis includes aspects of physiological dependence, loss of control over drinking, as well as persistent alcohol intake despite physiological, psychological and interpersonal consequences^6,7^. In contrast, typical alcohol intake, as assessed using measures such as drinks per week, represents the distribution of alcohol use from casual or social drinking to excessive drinking demarcating risk for AUD^8,9^ While heritable, measures such as drinks per week (DPW) are more likely to be influenced by environmental and socio-cultural factors and have complex and variable associations with morbidity and mortality^8,9^

Genome-wide association studies (GWASs) of AUD and DPW have identified multiple risk loci. The largest GWAS of problematic alcohol use (PAU; N=435,563) which meta-analyzed AUD with a GWAS of the problem-subscale of the Alcohol Use Disorders Identification Test (AUDIT-P) reported genome-wide associations at 29 loci encompassing 66 genes, the largest tranche of signals for any addictive disorder to date^10^. Alongside, the largest GWAS of typical alcohol intake (N=941,280) identified more than 200 independent genome-wide significant variants within or near more than 150 genes at 81 independent loci^11^. Despite a genetic correlation (SNP-rg) of 0.77 between PAU and DPW (less so for AUD and DPW, SNP-rg=0.67), genetic correlations between these aspects of alcohol involvement and other anthropometric, cardio-metabolic and psychiatric disorders revealed marked distinctions^10–14^. For instance, while AUD and PAU appear to be consistently associated with increased genetic liability for other psychiatric disorders and negatively with liability to educational achievement, DPW is genetically uncorrelated with most psychiatric disorders (except ADHD and tobacco use disorder) but correlated negatively with educational achievement and cardio-metabolic disease (which remains uncorrelated with PAU or AUD)^10–14^. These findings strongly hint at some common pathological underpinnings to AUD, PAU and other mental illnesses while genetic liability to DPW appears to be confounded with socio-economic correlates of alcohol use^10,11,13,14^.

Few studies have examined the intersection between the loci and genes associated with AUD and DPW, especially with respect to their functional and regulatory significance. As observed in other large GWAS, most genome-wide significant variants associated with AUD and DPW are intergenic and thus not directly mappable to a specific gene^15,16^. Furthermore, positionally mapping a non-coding variant to the nearest gene often does not identify the causal gene(s)^15–17^. Indeed, most variants identified by GWAS reside within and affect the activity of regulatory elements (e.g., enhancers and promoters) that regulate the expression of target causal genes in specific cell types; the affected genes are often located at quite a distance from the risk variant/regulatory element^13,15,16,18^. Several recent studies have integrated GWAS data with expression

QTLs (eQTLs) using colocalization or integration methods to identify causal variants and genes associated with schizophrenia, Alzheimer’s disease and many other complex disorders^13,15,16,18^. While similar efforts have been targeted at AUD and DPW, they have predominantly relied on bulk mRNA expression data from the small number of brain tissue samples in GTEx^10–14^

In the current study, we used a multi-omics systems approach to identify causal variants and genes associated with AUD and DPW. Using Mendelian Randomization-based methods on the largest available transcriptomic and epigenomic data for brain tissues and myeloid cells, we prioritized regulatory variants that influence AUD and DPW, specifically as well as simultaneously. We also used mRNA expression data from the brains of individuals diagnosed with AUD and controls (N=138) to validate the differential expression of genes prioritized in the GWAS integration analyses. To our knowledge, this is the first and the largest systematic multi-omics integration analysis to identify the functional impact of variants and genes associated with two correlated but etiologically distinct aspects of alcohol involvement.

## Results

### A UD meta-analysis

The large meta-analysis of AUD GWAS summary statistics (N=48,545 AUD cases and 187,065 controls) from the Million Veterans Program (MVP)^19^, the Psychiatric Genetics consortium (PGC-SUD)^12^ and the Collaborative Studies on Genetics of Alcoholism (COGA)^20^ identified 1157 SNPs (31 independent lead SNPs) within or near 79 genes at 10 independent loci associated with AUD. We didn’t include UKB-AUDIT-P in this meta-analysis to specifically focus on AUD. Many of these loci were shared between the AUD GWAS and the drinks per week (DPW) GWAS by Liu et al^11^ who identified 81 independent loci represented by 5197 (> 200 independent lead) SNPs. A total of 360 SNPs associated with AUD and DPW were in common (i.e., p < 5 × 10^−8^ in both GWAS). A large proportion (45%) of AUD and DPW associated SNPs were within intronic, UTR and non-coding regions of the genome.

### LDSC analysis using tissue specific epigenetic annotations

We used the stratified linkage disequilibrium score (LDSC) regression^21^ to test whether the heritability of AUD and DPW is enriched in regulatory regions surrounding genes in a specific tissue. Using multi-tissue chromatin (ROADMAP and ENTEX) data^22^, we observed a significant enrichment of promoter-specific epigenetic markers (H3K4me1/me3) in the fetal and the adult (germinal matrix, frontal-cortex) brain (P < 5 × 10^−8^) (Figure 2; Supplementary table 1) for the SNPs associated with AUD and DPW respectively.

### Integration of GWAS and eQTL/mQTL data from fetal and adult brain

Summary based Mendelian Randomization (SMR) analysis of genome-wide AUD summary statistics with eQTLs and mQTLs in the adult and fetal brain identified 21 genes at 18 loci across the genome (P-SMR FDR < 20%; Heidi > 0.05) [Supplementary Table 3a]. Among these 18 loci, SMR analysis nominated a single candidate causal gene at 16 loci, while more than 1 causal gene was nominated at 3p21.31 *(GPX1, AMT)* and 11p11.2 *(SPI1, MTCH2, NUP160).* To avoid the occurrence of false positive co-localizations that might be exclusively driven by stronger eQTL/mQTL signals, we focused on the loci where the strongest SNP was at least suggestively significant in the GWAS (P-GWAS < 5 × 10^−5^) [Table 1a; Figure 1]. Because of the much larger sample size of the DPW GWAS, 61 genes at nearly 31 loci passed the threshold for significance (P-SMR FDR < 20%; Heidi > 0.05; GWAS P < 5 × 10^−5^) [Table 1b; Figure 1]. On chromosome 11p11.2, our SMR based integration analysis co-localized a fetal brain specific mQTL *(SPIT)* and an adult brain specific eQTL *(NUP160)* with both traits (AUD and DPW). On chromosome 17q.21.31, the integration analysis prioritized different candidate genes for AUD *(MAP3K14’)* and DPW *(MAPT, CRHR1* and *LRRC37A).* Indeed, the AUD and DPW associations at 11p11.2 are likely to be two distinct loci, because the lead co-localized SNPs for each phenotype were not in LD (r^2^= 0.2). The DPW association tagged the H2 haplotype at 17q.21.31, while AUD’s association with *MAP3K14* was outside the inversion area (Supplementary figure), defined by the H1/H2 haplotypes in this region.

**Figure 1:**
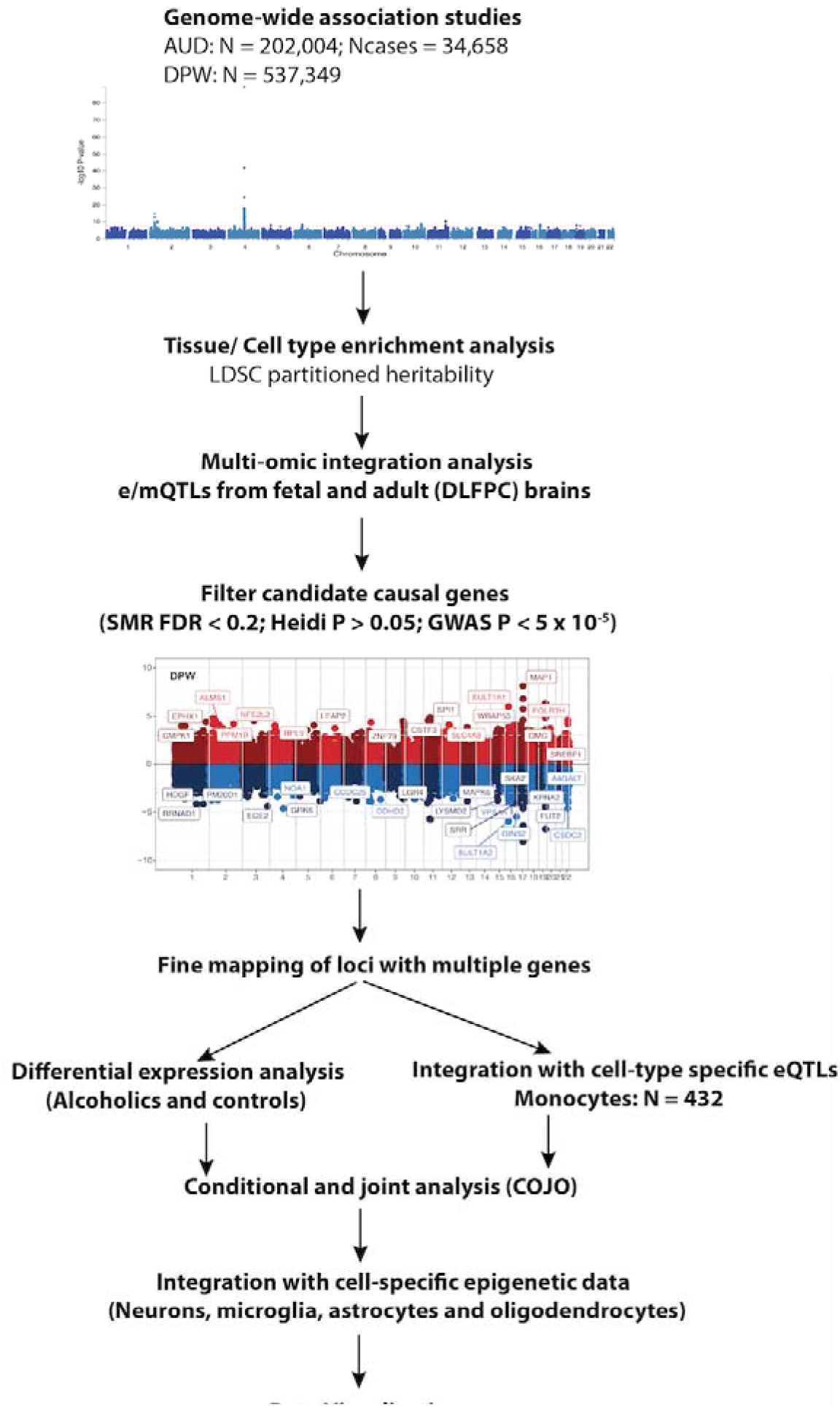
Overview of the study. Series of analyses were undertaken to identify the candidate causal genes associated with risk of AUD and DPW. We used the stratified linkage disequilibrium score (LDSC) regression to test whether the heritability of AUD and DPW is enriched in regions surrounding genes with chromatin markers in a specific tissue. This analysis helped us to identify the large eQTL/mQTLs datasets in the relevant tissues to perform the multi-omic integration analysis using SMR. The candidate causal SNPs and genes prioritized using SMR were further filtered according to threshold of association in GWAS and linkage disequlibrium (Heidi P and COJO). The complex loci with multiple genes were further validated and prioritized by exploring differential gene expression data from brains of alcoholics and controls. Integration of eQTL data from monocytes also helped to prioritize candidate genes specifically expressed in the myeloid cells. The cell type specific epigenetic data from the human brain was also used to identify the causal SNP/s associated with DPW and AUD.

**Table 1a:** SMR analysis results with summary statistics of AUD GWAS.

| Chr | BP | Gene | Adult brain |  | Fetal brain |  |
| --- | --- | --- | --- | --- | --- | --- |
|  |  |  | eQTL<br>p value | mQTL<br>p value | eQTL<br>p value | mQTL<br>p value |
| 11 | 46399942 | *SPII* | x | x | x | 1.91E-04 |
|  | 46664086 | *MTCH2* | 1.89E-05 | x | x | x |
|  | 46843734 | *NUP160* | 3.88E-04 | x | x | x |
| 3 | 48395716 | *GPX1* | 4.93E-05 | x | x | x |
|  | 48459884 | *AMT* | 2.07E-04 | x | 4.39E-04 | 4.47E-01 |
| 17 | 43361331 | *MAP3K14* | x | x | x | 2.99E-05 |

Column names: Chr (Chromosome), BP (Start position of the gene), Gene (Candidate causal gene), p value (SMR P values for integration of AUD summary statistics with respective eQTL/mQTL annotation from adult or fetal brain). X denotes the missing values either due to non-significant results or sub-threshold expression or methylation values in the respective tissue.

**Table 1b:**
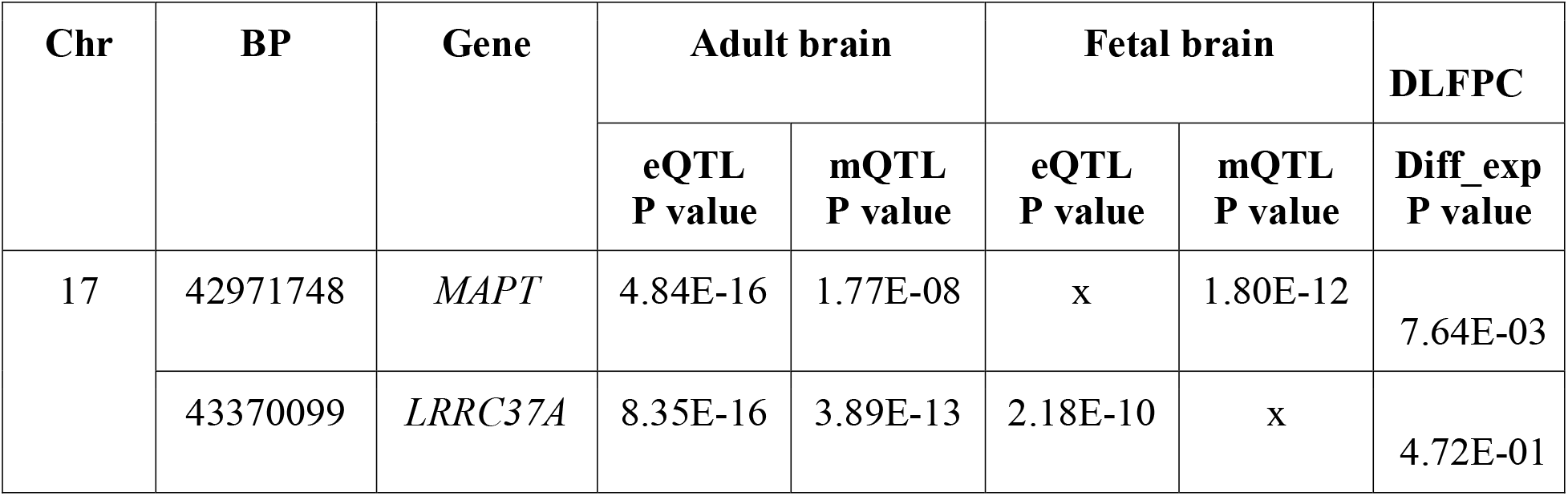
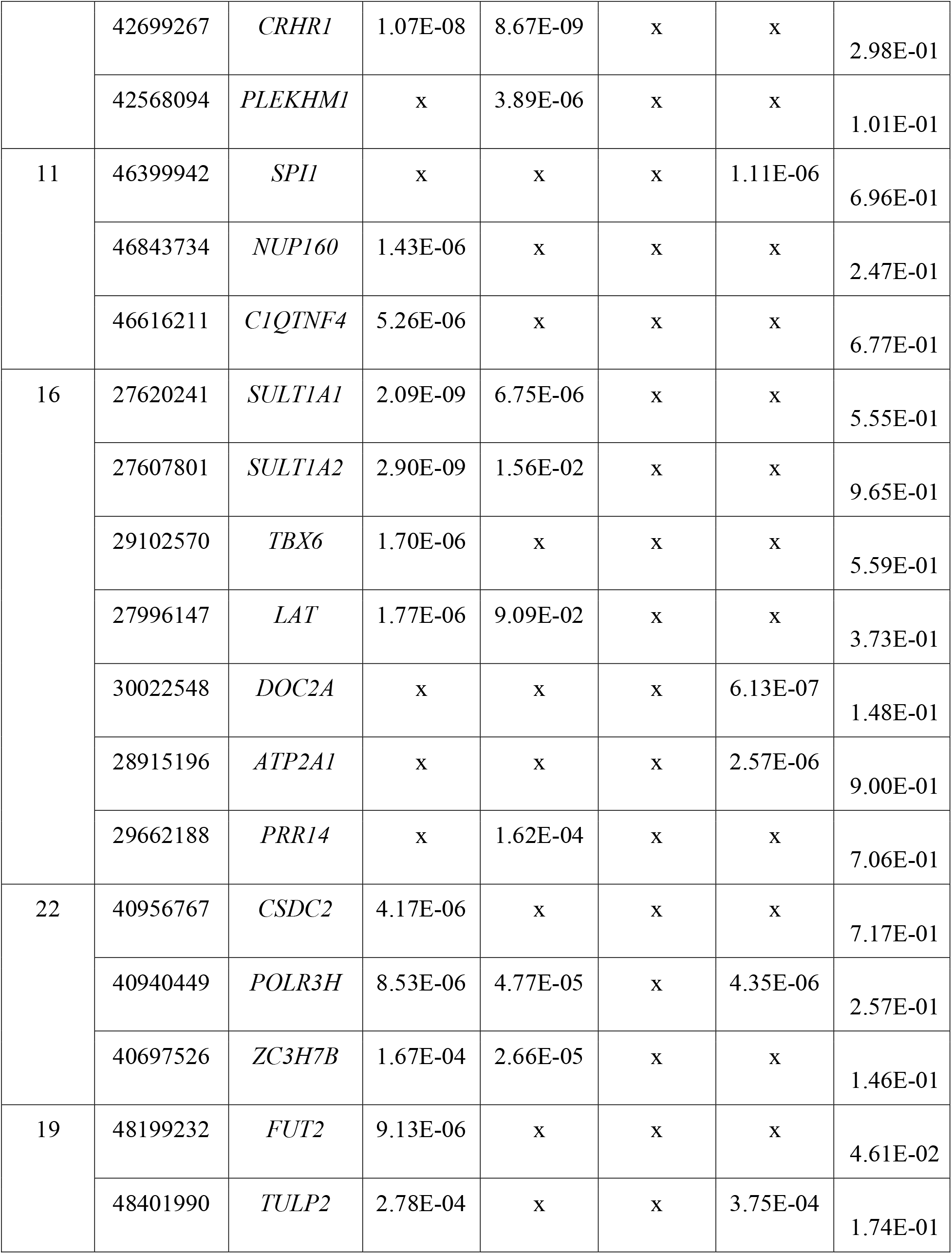
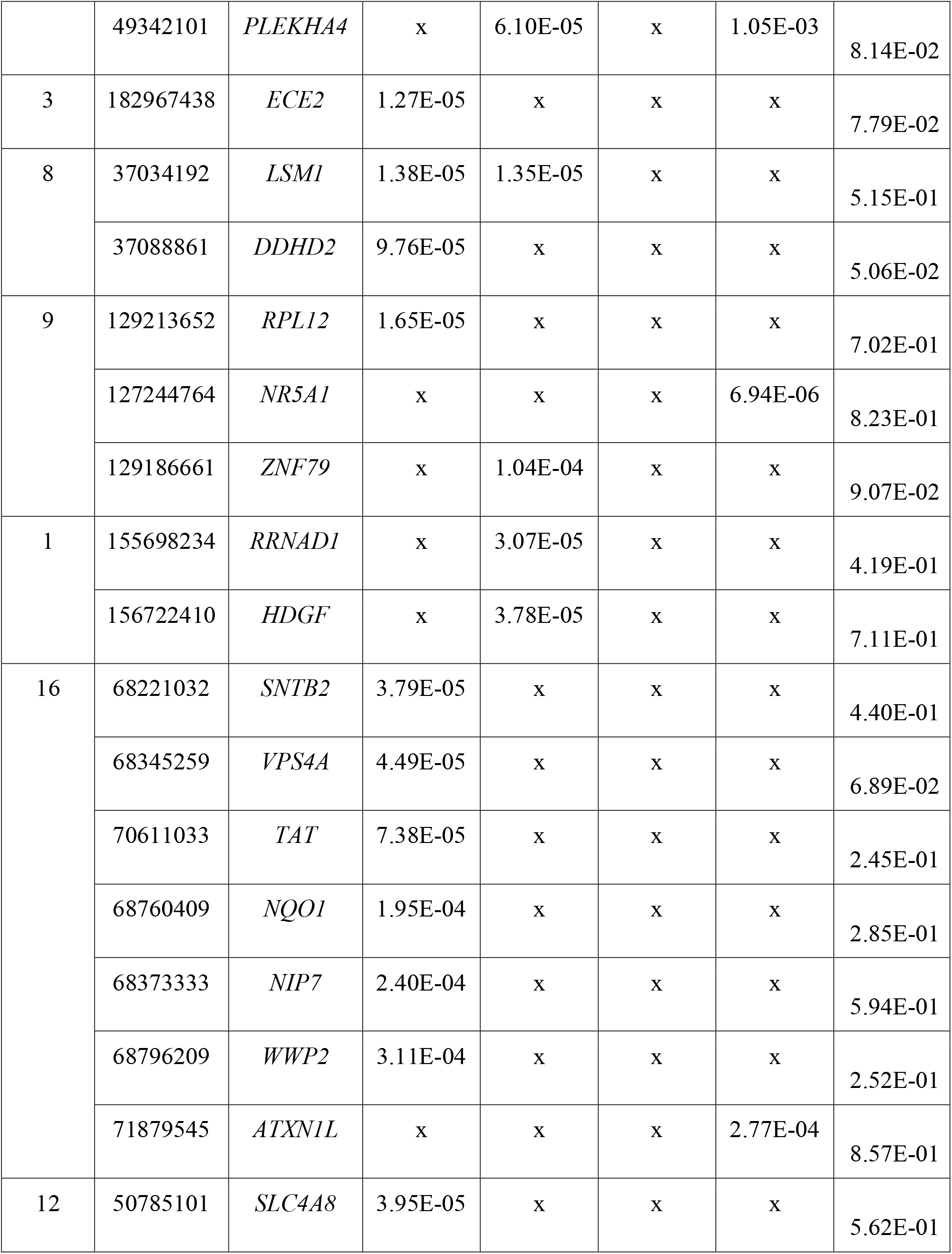
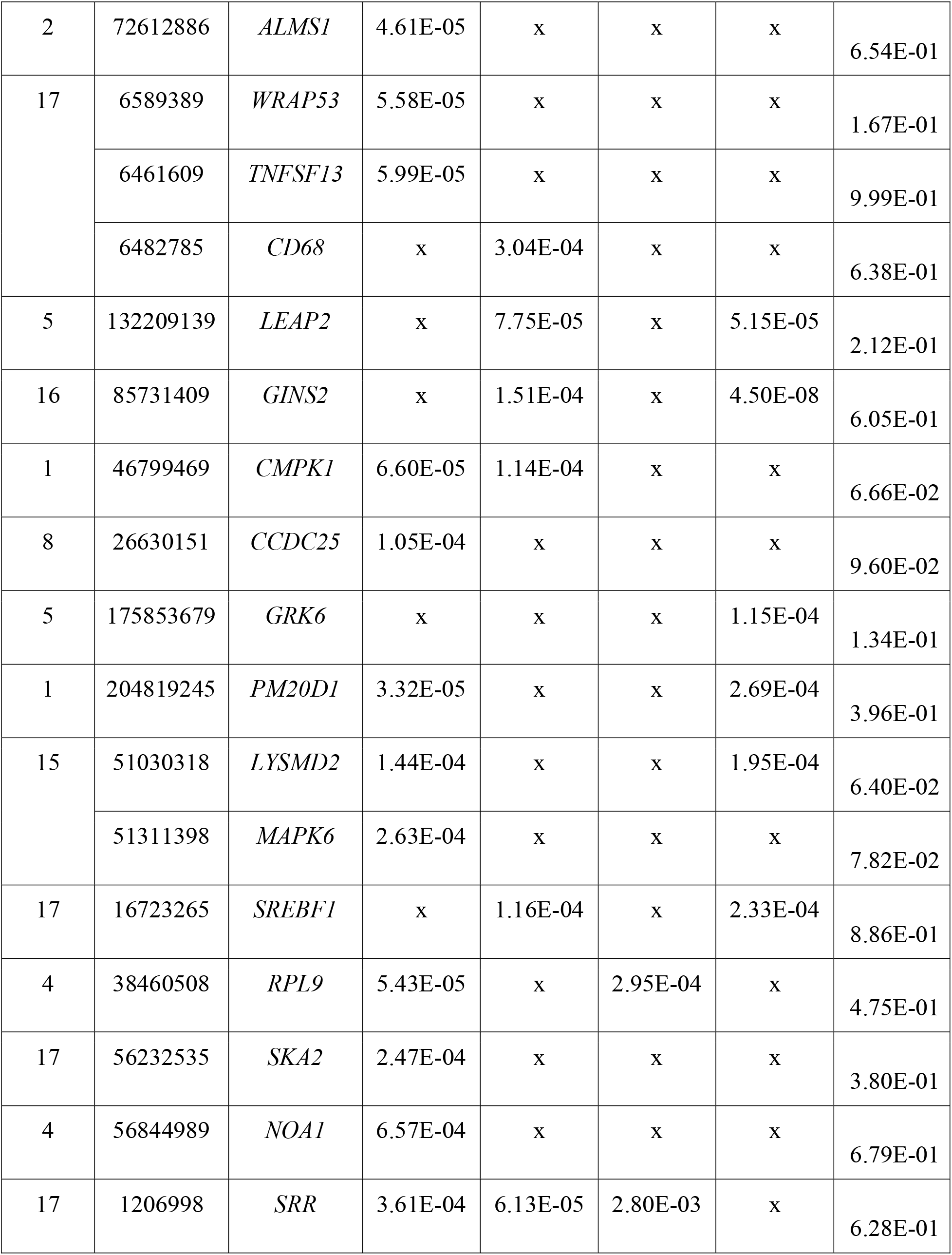
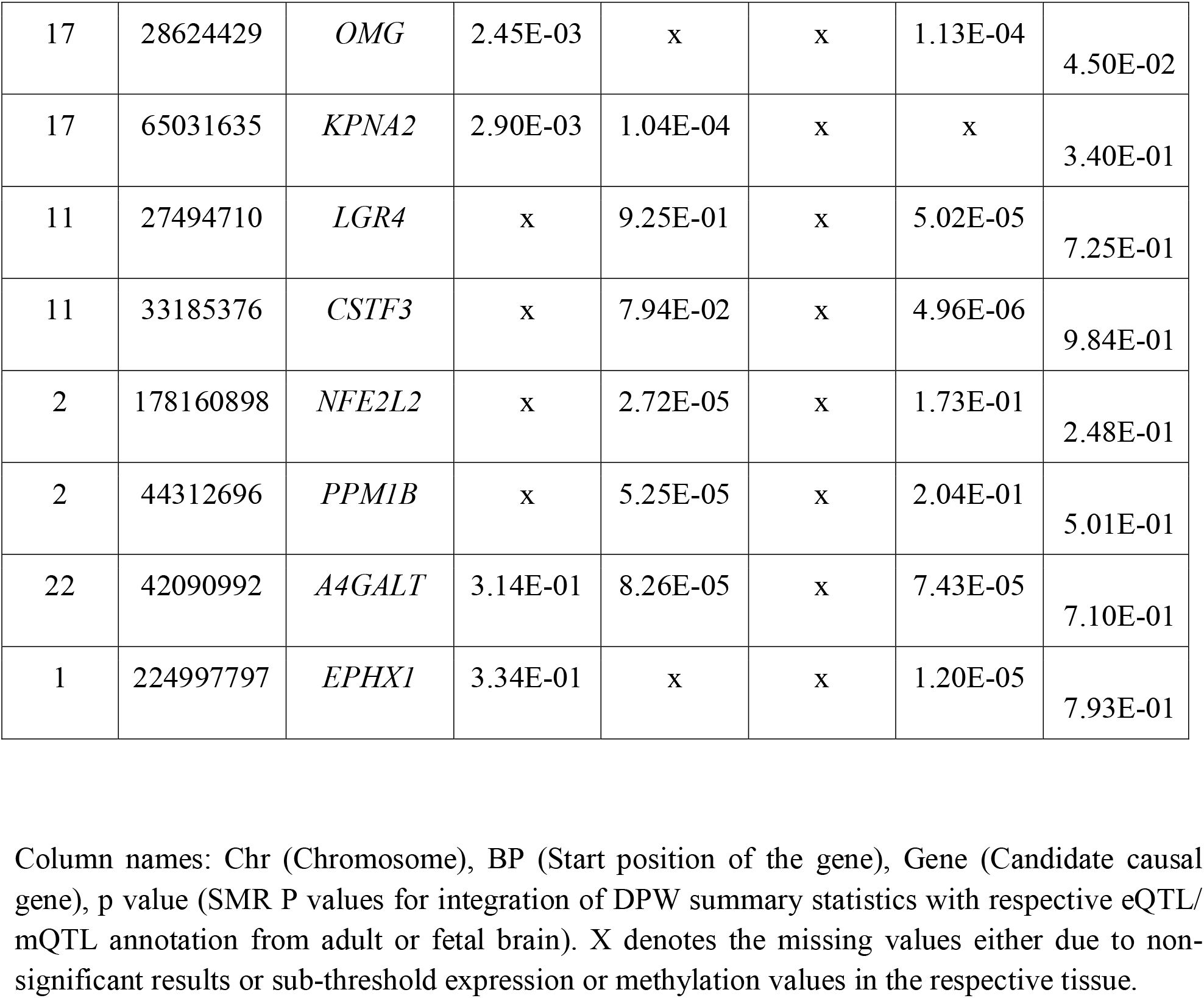
SMR analysis results with summary statistics of DPW GWAS.

### Fine mapping of 17q.21.31

At *17q.21.31,* eQTL and/or mQTL from both fetal and adult brains co-localized with DPW signals. We observed stronger evidence of co-localization (SMR P < 5 × 10-15) for DPW with *MAPT* and *LRRC37A,* than at any other locus. These genes are within a large inversion polymorphism (approximately 900 kb) that arose about 3 million years ago^23^. Since that time, these haplotypes have been recombinationally suppressed and have accumulated many haplotype-specific variants. As a result, there is extended LD within more than 1 Mb, which makes it difficult to fine map the causal variants and genes at this locus. Integration analysis of adult brain eQTL data with the DPW GWAS predicted that increased *MAPT* expression (SMR Beta = 0.01) is associated with increased number of alcoholic drinks per week, while decreased expression of *LRRC37A* (SMR beta = −0.02) was associated with a decrease in drinks per week. The predicted gene expression results from AUD and DPW GWAS were compared with observed expression differences in the brains of AUD subjects and controls to validate the results. Our differential expression analysis of alcohol consumption in the human brain indeed showed that the mRNA expression of *MAPT* was associated with increased alcohol consumption (Figure 2c). We did not observe any association between expression of *LRR37A* and level of alcohol intake. The co-localized SNPs within the 17q.21.31 locus were also compared with the promoter (H3K27ac, H3K4me3), enhancer (ATAC-Seq) and promoter-enhancer interactome (PLAC-Seq) data from 4 specific brain cell-types (microglia, neuron, astrocytes and oligodendrocytes) to elucidate the functional significance of these variants (Supplementary table). The co-localized mQTLs (rs3785884 and rs17651887) overlapped with the chromatin interaction region specifically in oligodendrocytes and these interactions looped at the *MAPT* promoter (Figure 2b). This observation combined with differential expression data in the human brain provides strong supporting evidence that *MAPT* is likely to be the causal gene at this locus associated with increased alcohol consumption.

**Figure 2:**
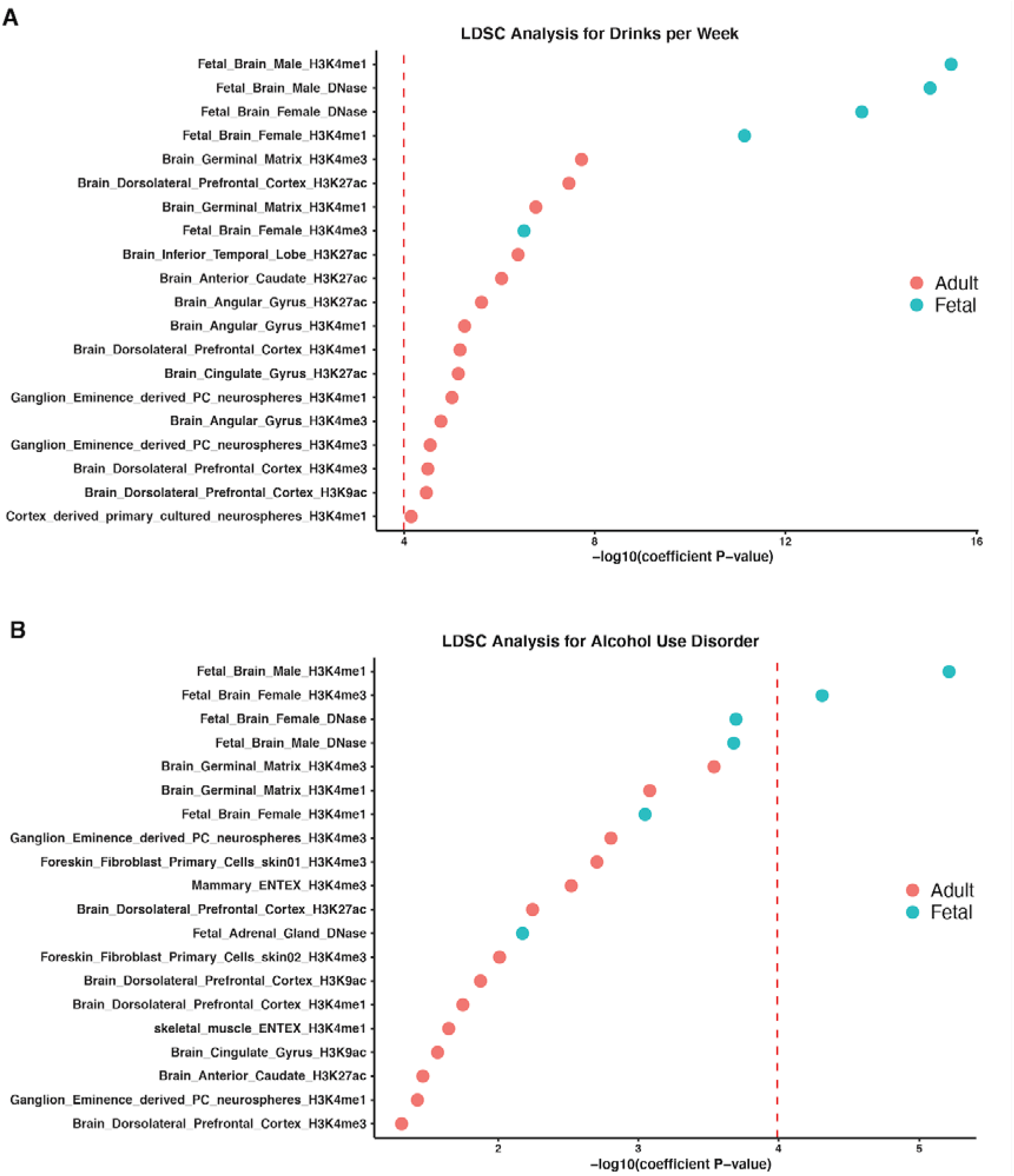
LDSC analysis using tissue specific chromatin data. LDSC analysis showed significant enrichment of promoter specific markers (H3K4me1/me3) in the fetal and adult brain for the SNPs identified in A) DPW and B) AUD GWAS analysis. Y-axis represents the annotations and X-axis represents the −log 10 P value for enrichment. The dotted red line represents the threshold of multiple test correction according to Bonferroni.

### Fine mapping at 11p11.2

SMR analysis with mQTLs from fetal brain and eQTLs from adult brain prioritized *SPI1* and *NUP160* respectively at *11p11.2* for association with DPW and AUD (Figure 3A). Both *SPI1* and *NUP160* are primarily expressed in myeloid cells. The predicted causal *SPI1* mQTL (rs56030824) and *NUP160* eQTL (rs10838753) were in fact in low LD (R^2^= 0.31) with each other. rs56030824 (mQTL) showed a stronger association with both AUD (P = 8.91 × 10^−6^) and DPW (4.90 × 10^−12^) than rs10838753 (eQTL) (DPW P = 1.28 × 10^−10^; AUD p = 4.85 × 10^−5^). Adding rs56030824 as a covariate in conditional analyses had a larger effect on the association between rs10838753 and AUD (P_orig_= 4.85 × 10^−5^; P_cond_= 0.09) than with DPW (P_orig_= 1.28 × 10^−10^; P_cond_ = 8.1 × 10^−3^). rs56030824 remained significantly associated with both AUD (P_orig_ = 8.91 × 10^−6^; P_Cond_ = 2.0 × 10^−2^) and DPW (P_orig_ = 4.90 × 10^−12^; P_cond_ = 7.0 × 10^−4^) even after adding rs10838753 as a covariate. Rs56030824 overlapped the promoter marks (H3K27ac, H3K4me3) for *SPI1* specifically in microglia (Figure 3A). This SNP also alters the binding site regulatory motif for RXRA, a transcription factor which is involved in promotion of myelin debris phagocytosis and remyelination by macrophages^24^. Since *SPI1* is expressed in myeloid lineage cells, its mRNA expression in the bulk brain was too low to perform differential expression or integration analysis. Therefore, we chose eQTLs from a large sample of peripheral blood monocytes to examine if rs56030824 is associated with expression of *SPI1* in these cells. The effect sizes of eQTLs at *11p11.2* locus were linearly correlated with effect sizes from the DPW GWAS at this locus (Figure 3B and 3C). In fact, rs56030824 had the strongest effect size for *SPI1* expression and DPW in the common variant category (Figure 3D). These observations together established rs56030824 as a stronger candidate to be considered as a causal variant and *SPI1* as a potential candidate gene associated with AUD and DPW.

**Figure 3:**
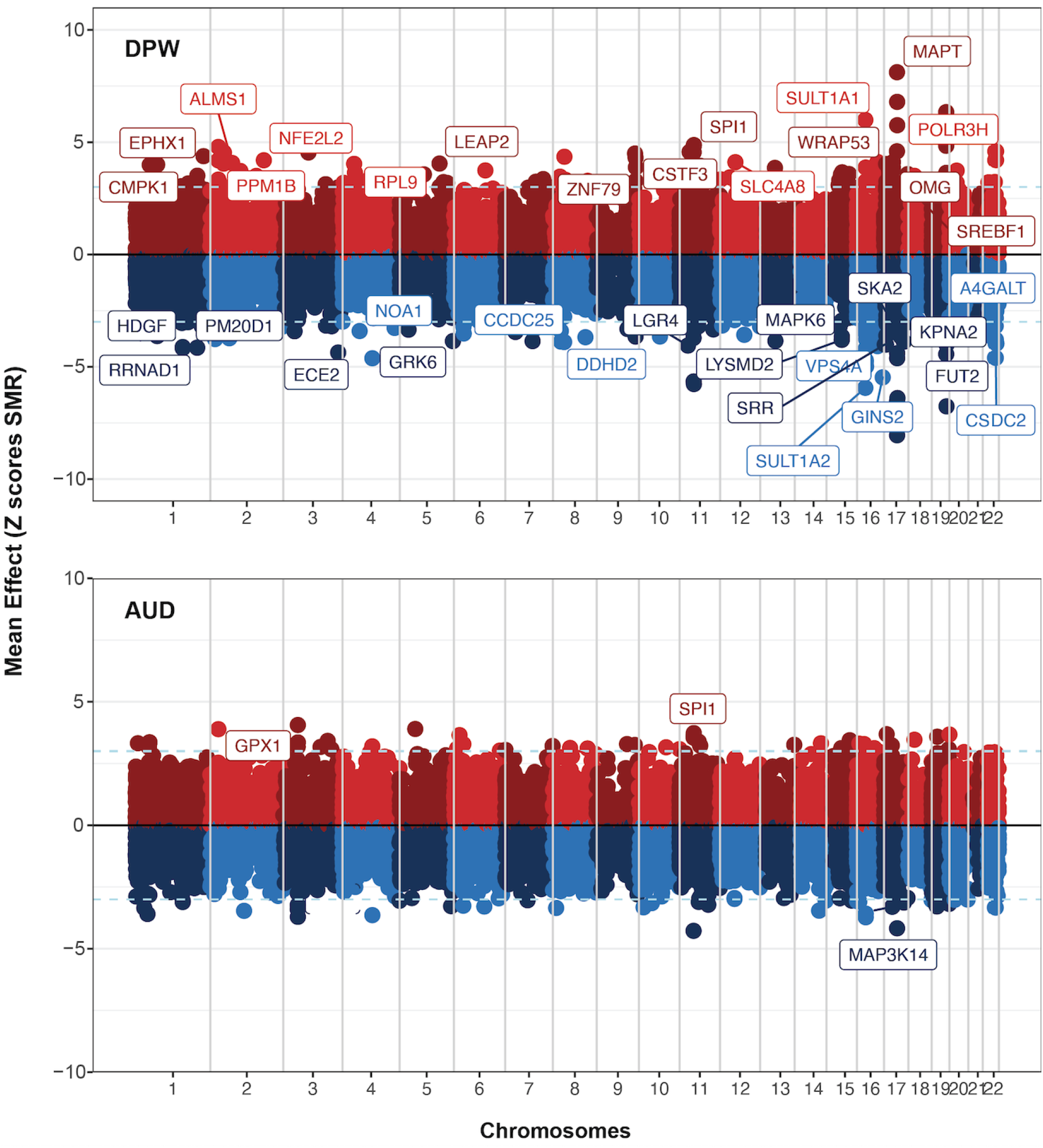
Results of SMR based integration analysis of DPW and AUD GWAS with eQTL/mQTL from fetal and adult brain. X-axis represents the chromosomes and Y axis shows the direction of effect (Z scores) on gene expression/methylation. Genes marked on the plots represent the genes nominated through strict threshold of co-localization (FDR < 20%; SMR_Heidi_ P > 0.05; GWAS P < 5 × 10^−5^) and/or multiple levels of transcriptomic and epigenetic evidence.

**Figure 4:**
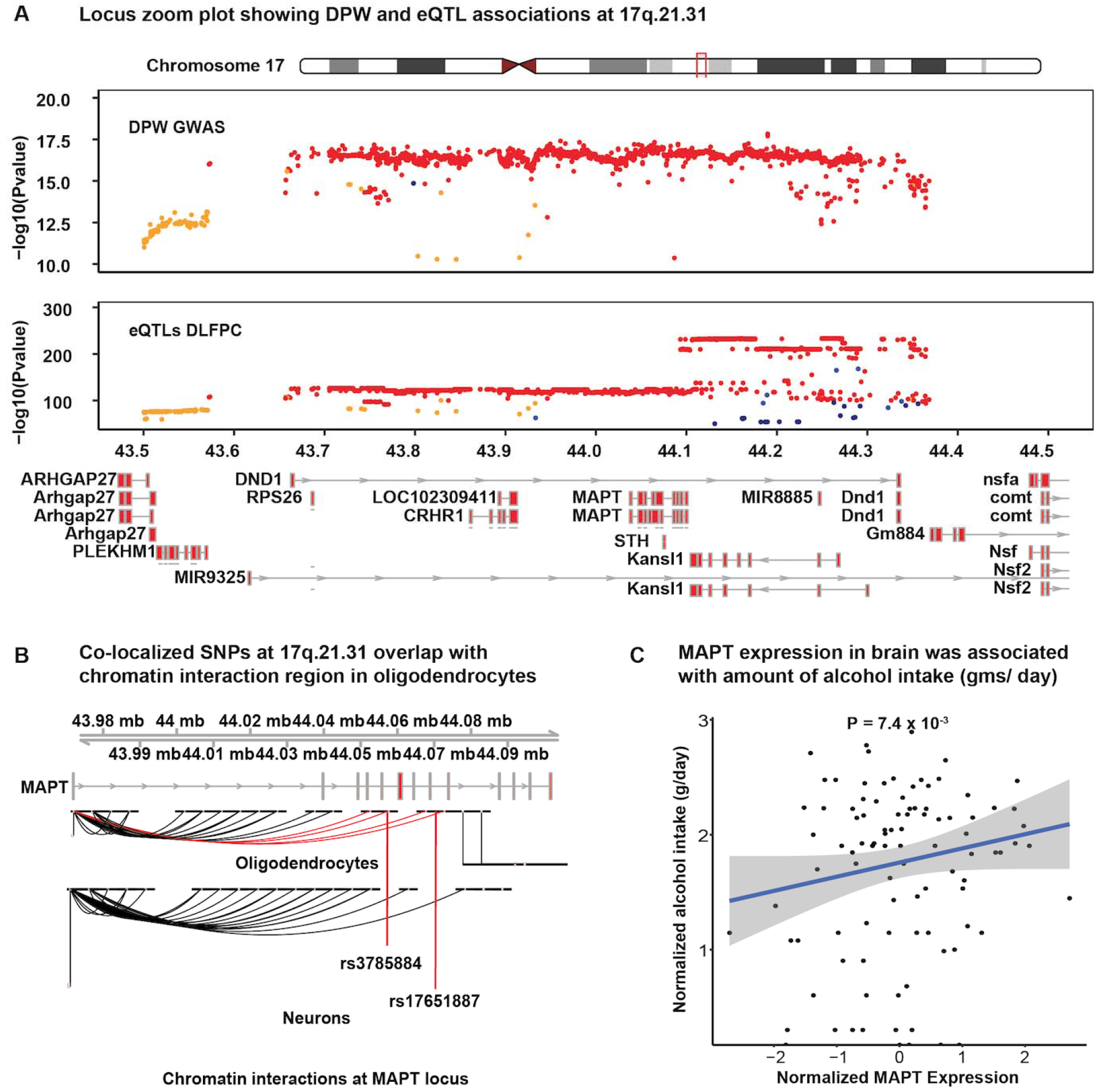
*MAPT* was identified as a candidate gene associated with increased DPW. A. Locus zoom plot showing DPW and eQTL (DLFPC) associations at 17q.21.3. X-axis represents the positions along chromosome 17 and y-axis represents the P values of each SNP at this locus. Color of each dot presents the R^2^ for LD at the locus (Red = 0.8 - 1.0; Orange 0.6-0.79; Green 0.4-0.59; Blue 0.2-0.39 and dark blue < 2.0). B. The co-localized SNPs were found to be overlapping with the chromatin interaction region that loops back to the promoter of the *MAPT* gene. C. In independent transcriptomic data from the human brain (N = 92), mRNA expression of MAPT was found to be associated with the alcohol consumption.

**Figure 5:**
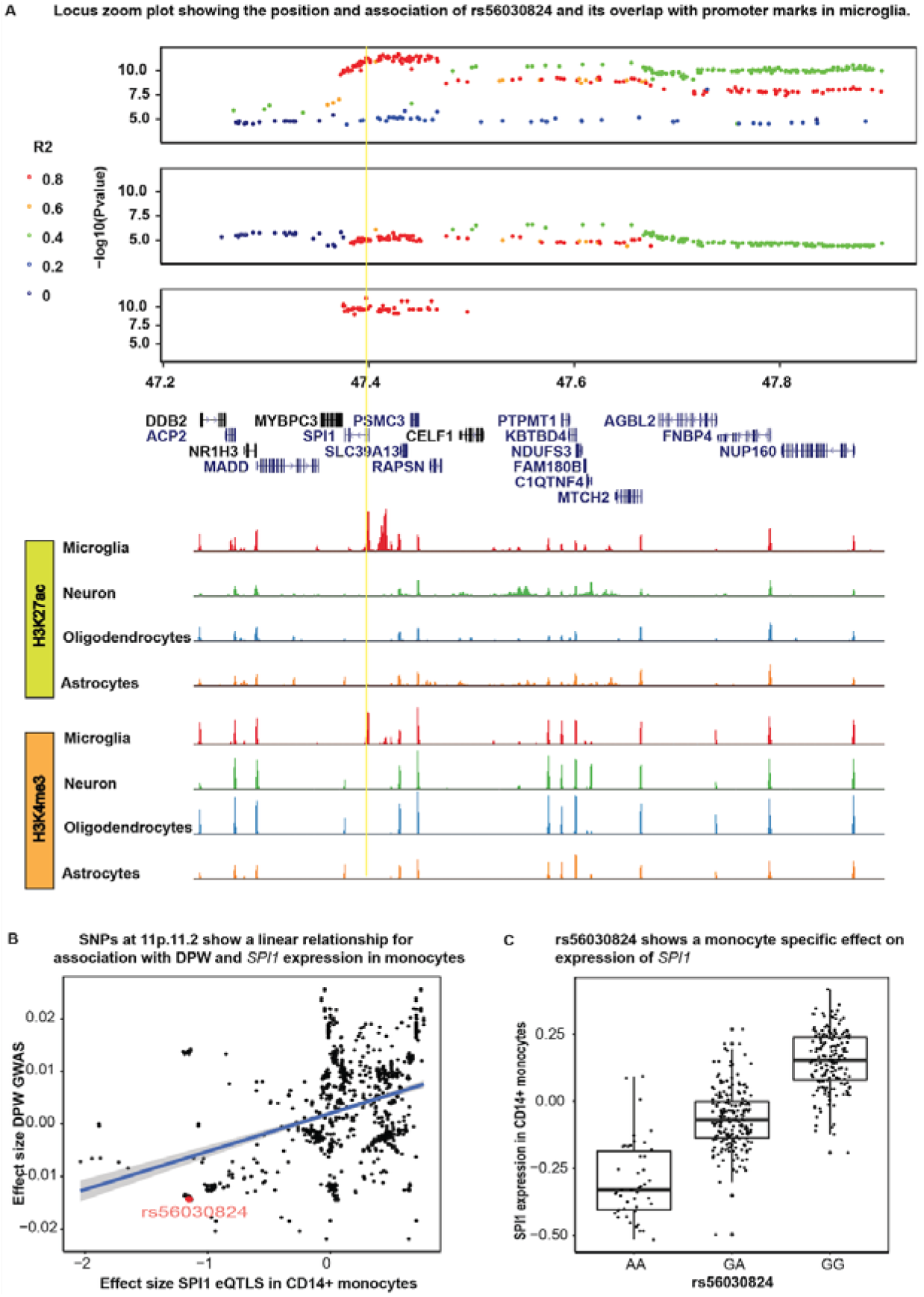
*SPI1* was nominated as candidate gene associated with increased DPW and AUD. Locus zoom plot showing DPW, AUD and mQTL (Fetal brain) associations at 11p.11.2. X-axis represents the positions along chromosome 11 and y-axis represents the P values of each SNP at this locus. Color of each dot presents the R^2^ for LD at the locus (Red = 0.8 - 1.0; Orange 0.6-0.79; Green 0.4-0.59; Blue 0.20.39 and dark blue < 2.0). Yellow line represents the position of rs56030824 identified as a functional variant co-localized with AUD, DPW and mQTLs in the fetal brain. The tracks show the peaks for promoter marks in 4 major cell types of the brain. Rs56030824 was found to overlap with promoter specific marks (H3K4me3 and H3K27ac), specifically in microglia. B) Effect sizes for DPW GWAS and *SPI1* expression in CD 14+ monocytes were found to be correlated i.e. decreased alcohol intake was associated with decreased *SPI1* expression. Rs56030824 showed the strongest association with DPW and mQTL in the common variant category. C) rs56030824 is a strong eQTL and associated with *SPI1* expression in CD14+ monocytes.

This study identified several other genes for DPW with multiple lines of evidence (eQTL, mQTL, differential expression; FDR < 20%; HEIDI P > 0.05; GWAS P < 5 × 10^−5^). For example, at locus 16p11.2, *SULT1A1* and *SULT1A2* were the strongest candidates with co-localization evidence emerging from mQTL and eQTLs from adult brain tissue (Table 1b). On chromosome 19, *FUT2* was the strongest candidate; mRNA expression of *FUT2* was also associated with increased alcohol consumption (Beta = 0.09, P = 4.6 × 10^−2^) when comparing the DLPFC of individuals with AUD and control subjects (Table 1b).

### Pathway and network analysis

Ingenuity Pathway Analysis of the prioritized genes associated with DPW showed significant enrichment for pathways related to TR (Thyroid hormone receptor)/RXR (Retinoic X receptor) activation, Lipoate biosynthesis, Estrogen biosynthesis and Sirutuin signaling (Figure 6, supplementary table 4). The DPW associated genes were also part of networks associated with immune cell trafficking and cellular movements (cell migration). Due to insufficient power and a smaller number of genes passing threshold of significance, we were not able to perform the pathway enrichment analysis for AUD.

**Figure 6:**
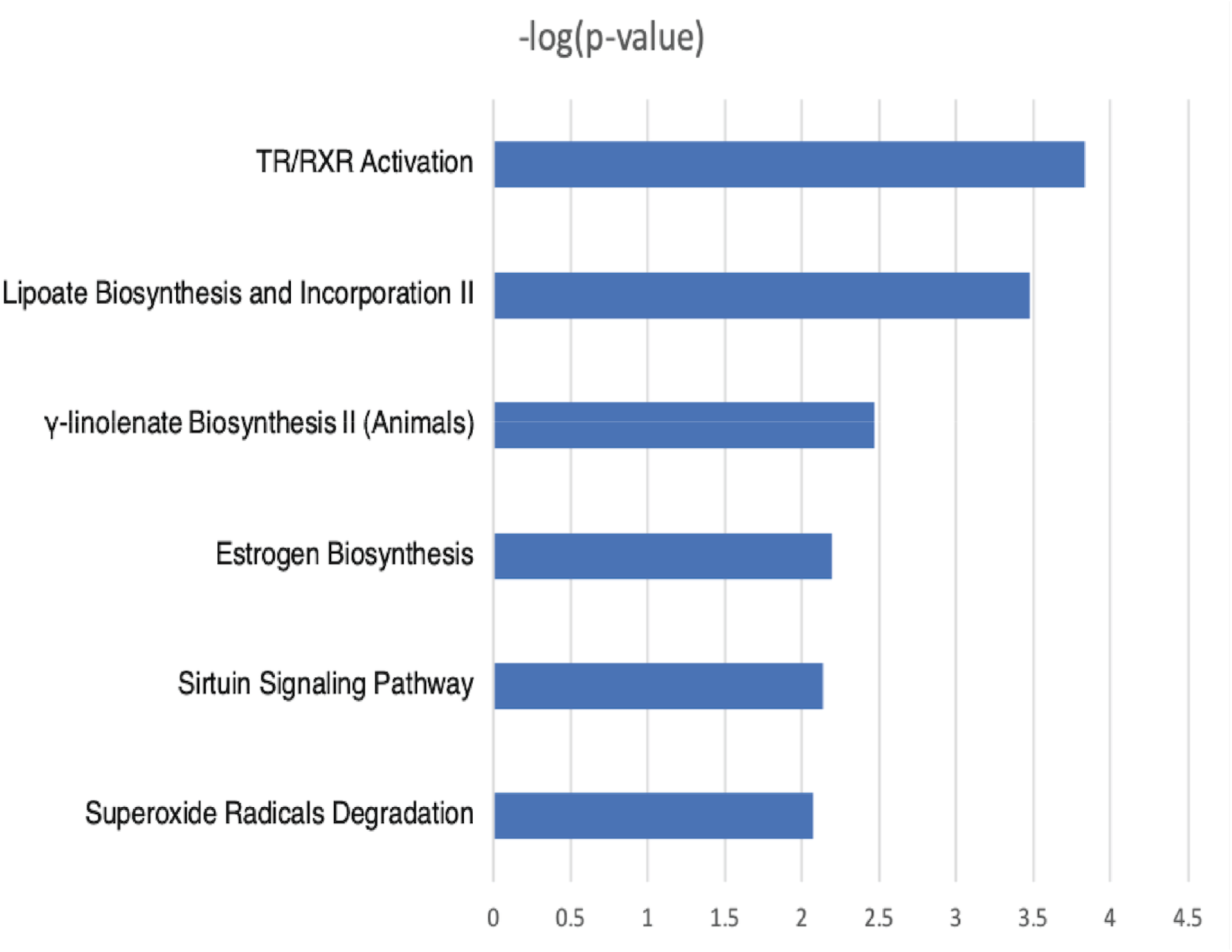
Genes nominated by multi-omic integration analysis were enriched in several pathways. The x-axis represents the −log 10 P value (Fisher’s exact test) and y-axis shows the significant pathways.

## Discussion

In this study, we used a multi-omics integration approach to detect genes relevant to typical drinking (DPW) and alcohol use disorder (AUD). The AUD GWAS used in the current analysis specifically focused on the diagnosis rather than the disordered drinking. Importantly, our work highlights that GWAS variants for alcohol use disorder and drinks per week are enriched in promoter regions of the fetal and adult brain. Using large-scale transcriptomic and epigenomic data from these tissues, we successfully fine mapped complex loci *(17q.21.31, 11p11N, 16p11.2)* and identified likely functional variants and candidate causal genes associated with alcoholism. Prior transcriptomic data from human and animal brains highlighted the contribution of immune networks in drinking behaviors^25–31^. But these observations were never consistent with results from GWAS of AUD and DPW, most likely due to lack of power in these genomic studies. Transcriptomic changes can be either a cause or a consequence of chronic excessive alcohol consumption. The identification of genes and/or pathways involved in immune signaling (*SPI1, RXR activation),* lipid metabolism (RXR and sulfotransferases) and regulation of alcohol metabolism (Sirutuin signaling) are therefore important as an attempt to fill gaps in our understanding of disease predisposition and underlying biological mechanisms in a genomic context.

For example, our LDSC based enrichment analysis shows that GWAS variants for both AUD and DPW are enriched in the genes expressed during early brain development. Drinking in later years might interact with this genetic predisposition making individuals more susceptible not only to AUD but also to other neuropsychiatric disorders^32^. Identification of *SPI1* and *MAPT* as genes for AUD are good examples of pleiotropy and/or causal links between the alcohol intake and susceptibility to AUD, other psychiatric disorders (e.g., depression) and even Alzheimer’s disease^15,33^ and other neurodegenerative diseases. We found that increased *SPI1* expression in myeloid lineage cells was associated with a higher DPW and higher risk for AUD. Recently, Zhang and colleagues^33^ observed that protein expression levels of *SPI1* in the cerebellum and spleen from subjects with Major depressive disorder and schizophrenia were significantly higher than in controls. In the past, we have also demonstrated that functional variants related to *SPI1* expression are associated with risk of Alzheimer’s disease^15^. Similarly, higher levels of expression are associated with increased risk for Alzheimer’s disease.

*SPI1* (Spi-1 Proto-Oncogene) encodes an ETS-domain transcription factor (PU.1) that regulates gene expression during myeloid and B-lymphoid cell development and homeostasis. This nuclear protein binds to a purine-rich sequence known as the PU-box found near the promoters of target genes and, in coordination with other transcription factors and cofactors, regulates their expression; among the genes are LXR/RXR nuclear receptors^34^. In the brain, *SPI1* is specifically expressed in microglia^15^. Given *SPI1’s* control over expression of several downstream genes, this gene may be a major reason enrichment of immune pathways is observed in transcriptomic analysis of human and animal brains. Because of the small fraction of microglia in bulk brain tissue, it is difficult to study expression of this transcription factor in transcriptomic datasets from whole brains. Some studies have reported that chronic alcohol consumption can influence expression of PU.1 in isolated microglia^35^ and peripheral lung macrophages^36,37^. However, these studies report the consequences of drinking on PU.1 expression whereas our study uses genomic evidence to demonstrate that regulation of innate immune response likely underlies, at least in part, susceptibility to increased drinking and eventual risk for AUD.

*MAPT* is another example of a pleiotropic relationship between AUD and other neuropsychiatric and neurodegenerative disorders. Located on chromosome 17, *MAPT*, encodes the tau proteins best known medically for their role in central nervous system disorders such as Alzheimer’s disease^38^, frontotemporal dementia^39^, Parkinson’s disease^38^, and the primary tauopathies progressive supranuclear palsy and corticobasal degeneration^40^. Recently, Hoffman and colleagues^41^ showed that alcohol use can upregulate the expression of pTau (Ser199/Ser202) in the hippocampus of C57BL/6J mice. Another study in humans observed differences in CSF-Tau levels in demented alcoholics vs Alzheimer disease patients^42^. *CRHR1* (corticotropin-releasing hormone type I receptor) is another gene on *17q.21.31,* that has been reported to be associated with alcoholism^43^. However, in our analysis, we did not observe an association between *CRHR1* expression and alcohol consumption

We also identified other genes that might be involved in increased alcohol consumption through a variety of biological mechanisms. For example, *VPS4A* at 16q23.1 has been implicated in dopamine regulation, reward anticipation, and hyperactivity in an fMRI study^44^. We also identified functional variants for *SULT1A1* and *SULT1A2* genes that encode for Sulfotransferase Family 1A enzymes catalyzing the sulfate conjugation of many hormones, neurotransmitters, drugs, and xenobiotic compounds^45^. In IPA disease enrichment analysis we also observed a significant overlap between genes implicated in DPW with other neurological, behavioral and immune related disorders (Supplementary Fig 1). The genes associated with DPW also showed significant enrichment for pathways related to TR/RXR activation, Lipoate biosynthesis, Estrogen biosynthesis and Sirtuin signaling (Figure 6, Supplementary Table). TRs (Thyroid hormone receptor) control the expression of target genes involved in diverse physiological processes and diseases, such as metabolic syndrome, obesity, and cancer, and, therefore, are considered as important targets for therapeutic drug development^46^. RXRs (Retinoic X Receptor) are also known to potentially regulate the ethanol metabolizing enzymes after chronic alcohol consumption^47^. It has been reported that the human aldehyde dehydrogenase-2 (*ALDH2*) promoter contains a retinoid response element, which might be contributing to the regulation of the gene^47^. Sirtuins signaling has been shown to play an important role in cocaine and morphine Action in the Nucleus Accumbens. Ferguson and colleagues^48^ demonstrated that systemic administration of a nonselective pharmacological activator of all sirtuins can increase the cocaine reward.

In conclusion, our study prioritizes risk variants and genes for subsequent experimental followup, which will help interrogate the molecular mechanisms underlying the link between alcohol consumption and AUD. Our database of multi-omics analysis in the fetal and adult brain is also made available with this study (see URLs) and provides a starting point to elucidate the biological mechanisms underlying AUD. We have demonstrated that individuals susceptible to AUD may have altered expression of disease-causing genes at earlier stages of life. Moreover, our results show the pleiotropic role of AUD-related variants in a variety of other brain disorders including Alzheimer’s disease. We expect results of multi-omic integration analysis will help researchers to design genetically informed experiments to identify biological mechanisms and drug targets related to AUD.

## Materials and methods

### Samples

#### Alcohol Use Disorder

We meta-analyzed three published GWAS: the Million Veteran Program (MVP)^19^ GWAS of AUD (EUR N = 202,004; Ncases = 34,658), with case status derived from International Classification of Diseases (ICD) codes of alcohol-related diagnoses from electronic health records (EHR) data, the Psychiatric Genomics Consortium (PGC) GWAS of alcohol dependence^12^ (cases based on DSM-IV diagnoses; EUR unrelated genotyped N = 28,757; Ncases = 8,485; AFR N = 5,799; Ncases = 2,991) and the Collaborative Studies on Genetics of Alcoholism (COGA) GWAS of alcohol dependence (cases based on DSM-IV diagnoses; EUR unrelated genotyped N = 4,849; Ncases = 2,411)^20^.

#### Drinks per week

We used genome-wide summary statistics for drinks per week (DPW; EUR N = 537,349 without the 23andMe samples) from GSCAN^11^ to contrast with AUD.

#### eQTLs from adult brain

We meta-analyzed three eQTL datasets with data from the dorsolateral prefrontal cortex (DLFPC): PsychEncode (N = 1387)^49^, ROSMAP (N = 461)^50^ and COGA-INIA (N = 138). We genotyped brain samples acquired from the New South Wales Brain Bank (NSWBB) using the UK Biobank Axiom array as part of COGA-INIA collaboration.

#### mQTLs from adult brain

Brain-mMeta mQTL summary data (Qi et al. 2018 Nat Commun)^51^ in SMR binary (BESD) format were obtained from the SMR data resource. This is a set of mQTL data from a metaanalysis of ROSMAP^52^, Hannon et al.^53^ and Jaffe et al.^54^

#### eQTLs from fetal brain

Summary data for eQTLs from developing human brains were obtained from an online repository shared by Heath O’Brien and Nicholas J. Bray^55^. The analyses were performed on 120 human fetal brains from the second trimester of gestation (12 to 19 post-conception weeks).

#### mQTLs from fetal brain

Summary data for mQTLs from developing human brains were obtained from an online repository shared by Ellis Hannon and Jonathan Mill^53^. The mQTLs were characterized in a large collection (n=166) of human fetal brain samples spanning 56-166 days post-conception, identifying >16,000 fetal brain mQTLs.

#### eQTL data from CD14+ monocytes

We used the gene expression and genotype data generated on primary monocytes from 432 healthy Europeans to quantify the relationship between the co-localized SNPs and expression of myeloid linseage gene^56^.

#### Whole genome transcriptomic data in brain of alcoholics

mRNA expression data in the DLFPC region of the human brain was generated in 138 brains obtained from the New South Wales Brain Bank (NSWBB). We also had access to alcohol consumption (gm/day) data in a subset of 92 brains. Alcohol-dependence diagnoses and consumption data were collected by physician interviews, review of hospital medical records, questionnaires to next-of-kin, and from pathology, radiology, and neuropsychology reports.

#### Brain cell type specific enhancer and promoter data

We used the promoter (H3K27ac, H3K4me3), enhancer (ATAC-Seq) and promoter-enhancer interactome (PLAC-Seq) data for 4 specific cell types of brain (microglia, neuron, astrocytes and oligodendrocytes) to elucidate the functional significance of colocalized SNPs^57^.

### Analysis

#### eQTL meta-analysis in adult brain

RNA Sequencing data on DLFPC region of the brain for 138 samples was generated as part of COGA-INIA collaboration^28^. We also genotyped the brain samples using the UK Biobank Axiom array. All NSWBB samples were imputed to 1000 Genomes using the cosmopolitan reference panel (Phase 3, version 5, NCBI GRCh37) using SHAPEIT then Impute2^58^ within each array. Only variants with non A/T or C/G alleles, missing rates < 5%, MAF >5%, and HWE P - values >.0001 were used for imputation. Imputed variants with R2 < .30 were excluded, and genotype probabilities were converted to genotypes if probabilities ≥90. All genotyped and imputed variants (4,615,871 SNPs) with missing rates < 10%, MAF ≥5% and HWE P-values >1 × 10^−6^ were included in the downstream analyses using MatrixQTL program. The gene expression was corrected for the batch, age, sex, rin, PMI and alcohol status using the “removeBatchEffect” option from limma package. The eQTL summary statistics from all three datasets were processed and munged together at single bp and allele level to remove ambiguity due to dbSNP rsids. The gene labels in all three datasets were also matched to Ensembl ids. The summary statistics were saved in binary format files (BESD) using the SMR. SMR “--meta” option was used to perform the meta-analysis in all three datasets.

#### LDSC analysis

We performed the partition heritability analyses for functional annotation using LDSC program. We obtained the weights for the multi-cell and tissue chromatin marks and performed the LDSC partition heritability analyses on munged summary statistics of AUD and DPW GWAS^21^.

#### SMR analysis

To examine whether the GWAS variants associated with both AUD and DPW are mediated by changes in methylation and gene expression patterns, we conducted a summary data-based Mendelian randomization (SMR) analysis^59^ on a set of mQTLs and eQTLs from fetal and adult brains. SMR is a Mendelian randomization-based analysis which integrates GWAS summary statistics with eQTL data in order to test whether the effect size of a SNP on the phenotype of interest is mediated by gene expression. We used this method as it does not require raw eQTL data to build the weights, so we were able to use the meta-analysis of eQTL data for the integration analysis. The gene and SNP positions for the summary statistics of the eQTL and mQTL datasets were standardized and aligned using an in-house summary statistics munging pipeline. The summary statistics were then converted into binary format (BESD) to perform the SMR analysis. The European subset of ADGC GWAS (phs000372.v1.p1) was used as the LD reference panel to perform the SMR analyses. The genes below FDR threshold (FDR < 20%) and with heterogeneity P value > 0.05 were considered to be causal within each combination of analysis.

#### Conditional and joint (COJO) analysis

We used a summary data based conditional analysis approach to identify the independent lead SNP associated with AUD and DPW. This conditional and Joint analysis (COJO)^60^ approach is implemented in Genome-wide Complex Trait Analysis (GCTA) software^61^ package and is valuable when the individual level genotype data is not available for the conditional analysis. To perform the COJO analysis we used the summary statistics of AUD and DPW GWAS along with the European subset of COGA samples as the LD reference panel.

#### Differential Expression analysis

We first performed a linear regression with alcohol intake as a dependent variable to identify possible covariates (e.g. sex, age, PMI). Gene-level analyses started with the featureCounts-derived sample-by-gene read count matrix. The basic normalization and adjustment pipeline for the expression data matrix consisted of: (i) removal of low expression genes (<1 CPM in > 50% of the individuals); (ii) differential gene expression analysis based upon adjustment for the chosen covariates. We filtered out all genes with lower expression in a substantial fraction of the cohort, with 18,463 genes with at least 1 CPM in at least 50% of the individuals; note that only these genes were carried forward in all subsequent analyses. The log10 normalized alcohol consumption (from NSWBB brains) was used for differential expression analysis using the DeSeq2 program. The analysis was controlled for sex, age, PMI, BMI, RIN, batch and severity of alcoholism (AUDIT scores).

#### Pathway analysis

The results of integration analysis for the DPW and AUD GWAS (FDR < 20%, Heidi P > 0.05) were used to perform gene ontology and pathway enrichment analyses using the EnrichR and Ingenuity Pathway Analysis (IPA).

#### Database for query and visualization

We used ShinyApp to create a database for query and visualization of the results of integration analyses. Users can create volcano, Manhattan plots and heatmaps to visualize the results of eQTL, mQTL and epigenetic integration analyses with summary statistics of AUD and DPW GWAS. Users will also be able to see whether the genes of interest are differentially expressed in the brains of alcoholics and controls.Data availability

Results of the SMR analysis for all the conditions can be found in Supplementary Tables 5a-5h. Additionally all the results can be visualized at our Shiny web app (https://lcad.shinyapps.io/alc_multiomics/).

#### Code availability

Standard tools (LDSC, SMR, GCTA-COJO, UCSC browser) were used to perform all the integrative analysis reported in this manuscript. Pipelines used in the analysis can be accessed at the GitHub repository (https://github.com/kapoormanav/alc_multiomics).

## Acknowledgements

COGA: The Collaborative Study on the Genetics of Alcoholism (COGA), Principal Investigators B. Porjesz, V. Hesselbrock, T. Foroud; Scientific Director, A. Agrawal; Translational Director, D. Dick, includes 11 different centers: University of Connecticut (V. Hesselbrock); Indiana University (H.J. Edenberg, T. Foroud, J. Nurnberger Jr., Y. Liu); University of Iowa (S. Kuperman, J. Kramer); SUNY Downstate (B. Porjesz, J. Meyers, C. Kamarajan, A. Pandey); Washington University in St. Louis (L. Bierut, J. Rice, K. Bucholz, A. Agrawal); University of California at San Diego (M. Schuckit); Rutgers University (J. Tischfield, A. Brooks, R. Hart); The Children’s Hospital of Philadelphia, University of Pennsylvania (L. Almasy); Virginia Commonwealth University (D. Dick, J. Salvatore); Icahn School of Medicine at Mount Sinai (A. Goate, M. Kapoor, P. Slesinger); and Howard University (D. Scott). Other COGA collaborators include: L. Bauer (University of Connecticut); L. Wetherill, X. Xuei, D. Lai, S. O’Connor, M. Plawecki, S. Lourens (Indiana University); L. Acion (University of Iowa); G. Chan (University of Iowa; University of Connecticut); D.B. Chorlian, J. Zhang, S. Kinreich, G. Pandey (SUNY Downstate); M. Chao (Icahn School of Medicine at Mount Sinai); A. Anokhin, V. McCutcheon, S. Saccone (Washington University); F. Aliev, P. Barr (Virginia Commonwealth University); H. Chin and A. Parsian are the NIAAA Staff Collaborators. We continue to be inspired by our memories of Henri Begleiter and Theodore Reich, founding PI and Co PI of COGA, and also owe a debt of gratitude to other past organizers of COGA, including Ting Kai Li, P. Michael Conneally, Raymond Crowe, Wendy Reich, and Robert E. Taylor, for their critical contributions. This national collaborative study is supported by NIH Grant U10AA008401 from the National Institute on Alcohol Abuse and Alcoholism (NIAAA) and the National Institute on Drug Abuse (NIDA). The authors acknowledge Office of Research Infrastructure of the National Institutes of Health under award numbers S10OD018522 and S10OD026880. Dr. Kapoor’s also receives part of his salary from NIAAA funded R21 grant R21AA026388.

## Competing Financial Interests Statement

Alison Goate is on the scientific advisory board for Denali Therapeutics and has served as a consultant for AbbVie and Cognition Therapeutics. Alison Goate is also listed as an inventor on Issued U.S. Patent 8,080,371, “Markers for Addiction” covering the use of certain variants in determining the diagnosis, prognosis, and treatment of addiction.

## Author Contributions

M.K. and A.G. planned the study and acquired the data for the analysis. M.K. performed the data analysis, created the visualization tools and wrote the first draft of manuscript. E.M, M.C., E. C.J., G.N., D.L. and J.S. assisted and helped in the data analysis. COGA collaborators including J.I.N., J.M., B.P., Y.L., T.F., H.J.E., and A.A. provide critical comments and suggestions on the manuscript.

## Notes

https://github.com/kapoormanav/alc_multiomics

https://lcad.shinyapps.io/alc_multiomics/

### Competing Interest Statement

Dr. Alison Goate is on the scientific advisory board for Denali Therapeutics and has served as a consultant for AbbVie and Cognition Therapeutics. Dr. Alison Goate is also listed as an inventor on Issued U.S. Patent (Markers for Addiction) covering the use of certain variants in determining the diagnosis, prognosis, and treatment of addiction.

