## Supplementary figures and images for "Multi-omics integration analysis identifies novel genes for alcoholism with potential link to neurodegenerative diseases"

### Supplementary figure 1

Analysis: genes_sig_dpw_brain - 2020-03-27 11:36 AM

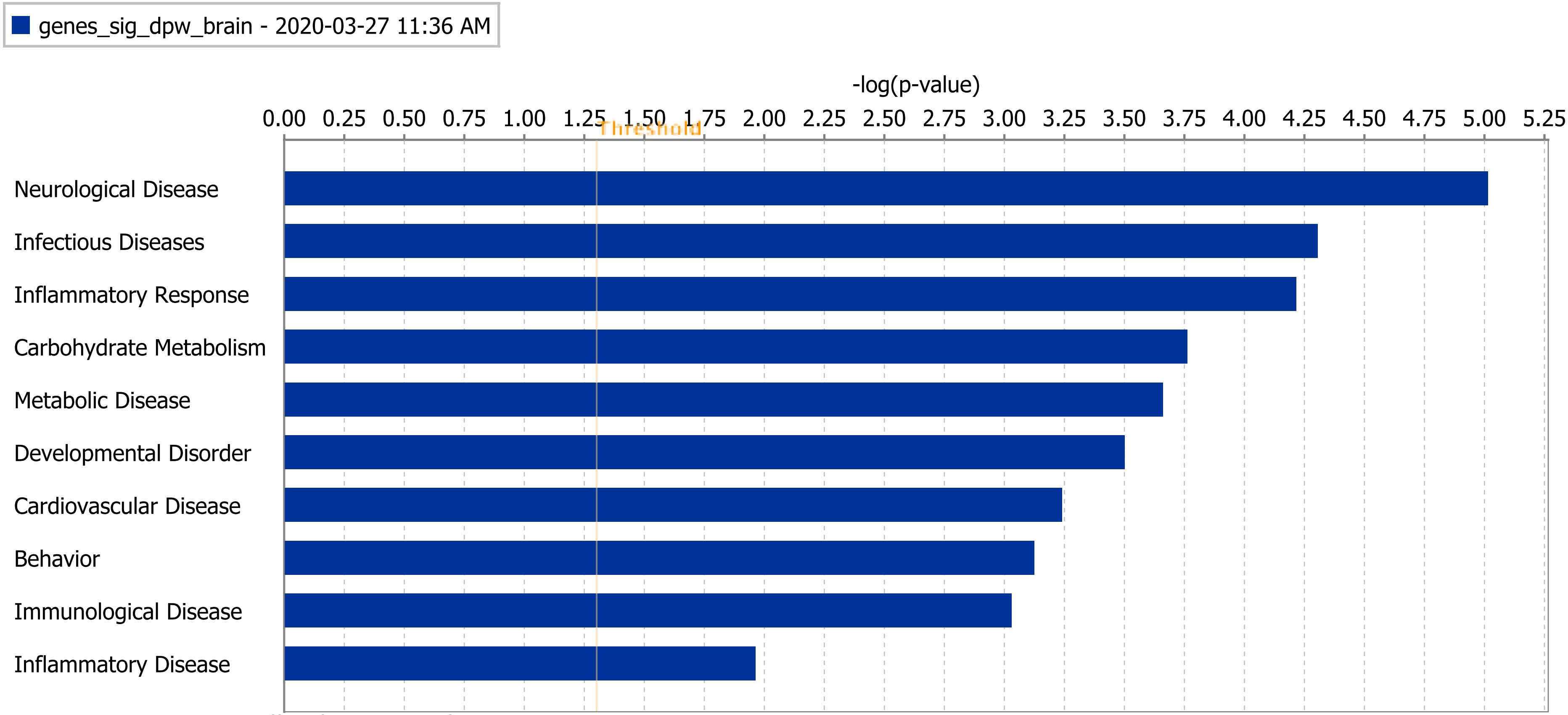


© 2000-2020 QIAGEN. All rights reserved.
