## Additional Supplementary data for "Multi-omics integration analysis identifies novel genes for alcoholism with potential link to neurodegenerative diseases"

**Additional Supplementary Figure 1: Manhattan plot for AUD GWAS meta-analysis**
**(FIgure generated through Fuma)**

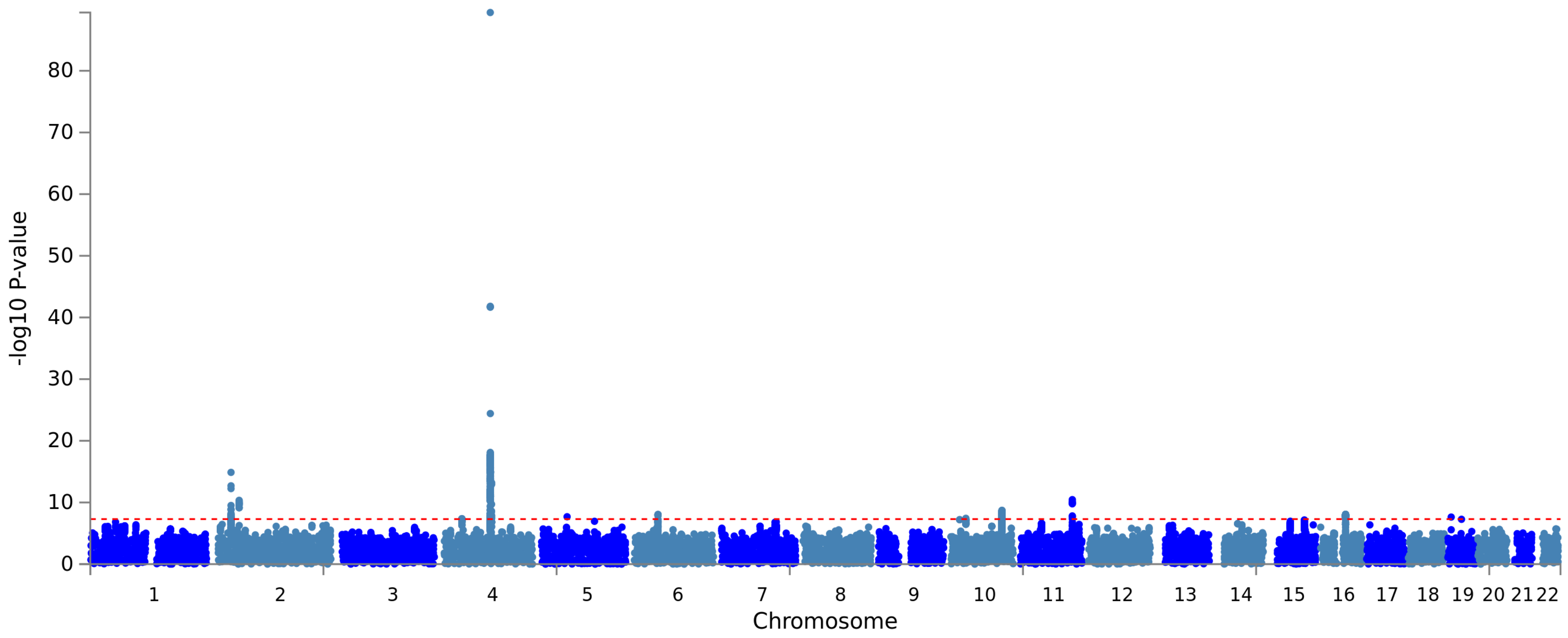

**Additional Supplementary Figure 2: Manhattan plot for gene-based association analysis results from MAGMA for AUD GWAS meta-analysis**
**(Figure generated through Fuma)**

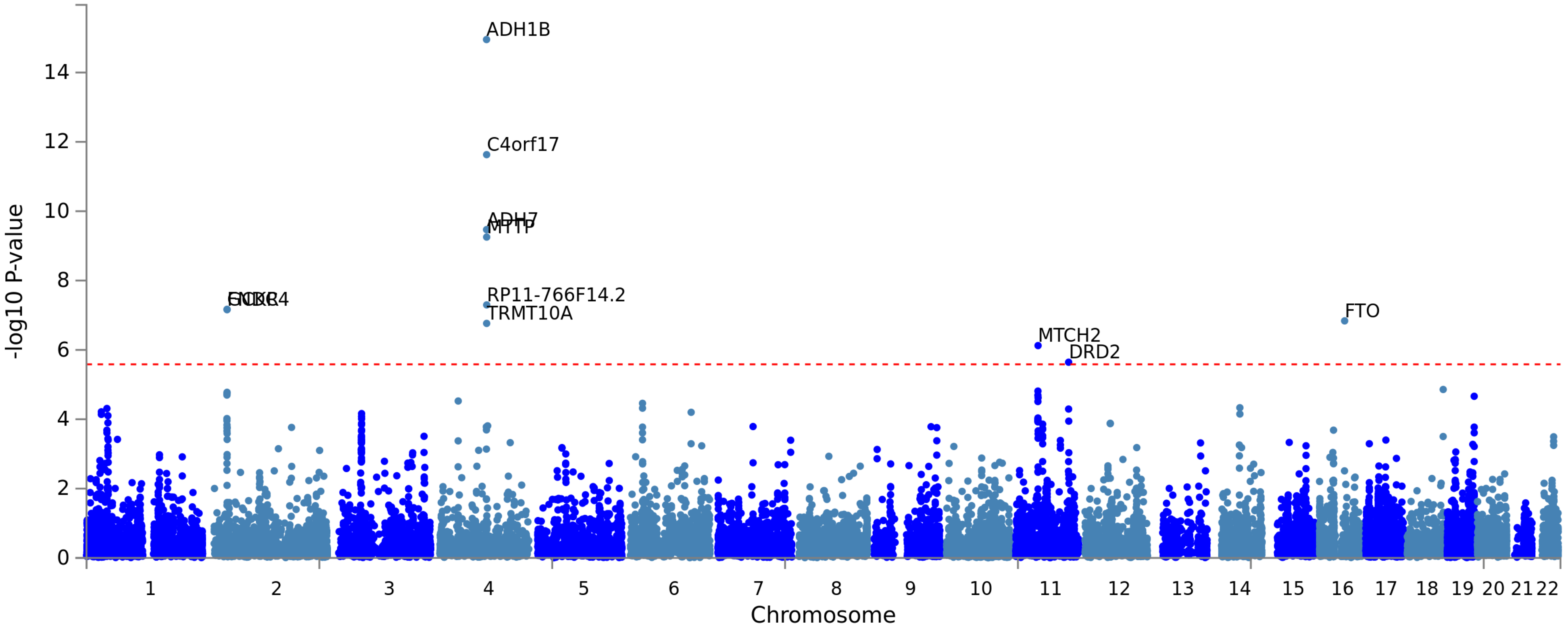

**Additional Supplementary Figure 3: Distribution of AUD GWAS SNPs across the genome (Annotations and information from FUMA)**

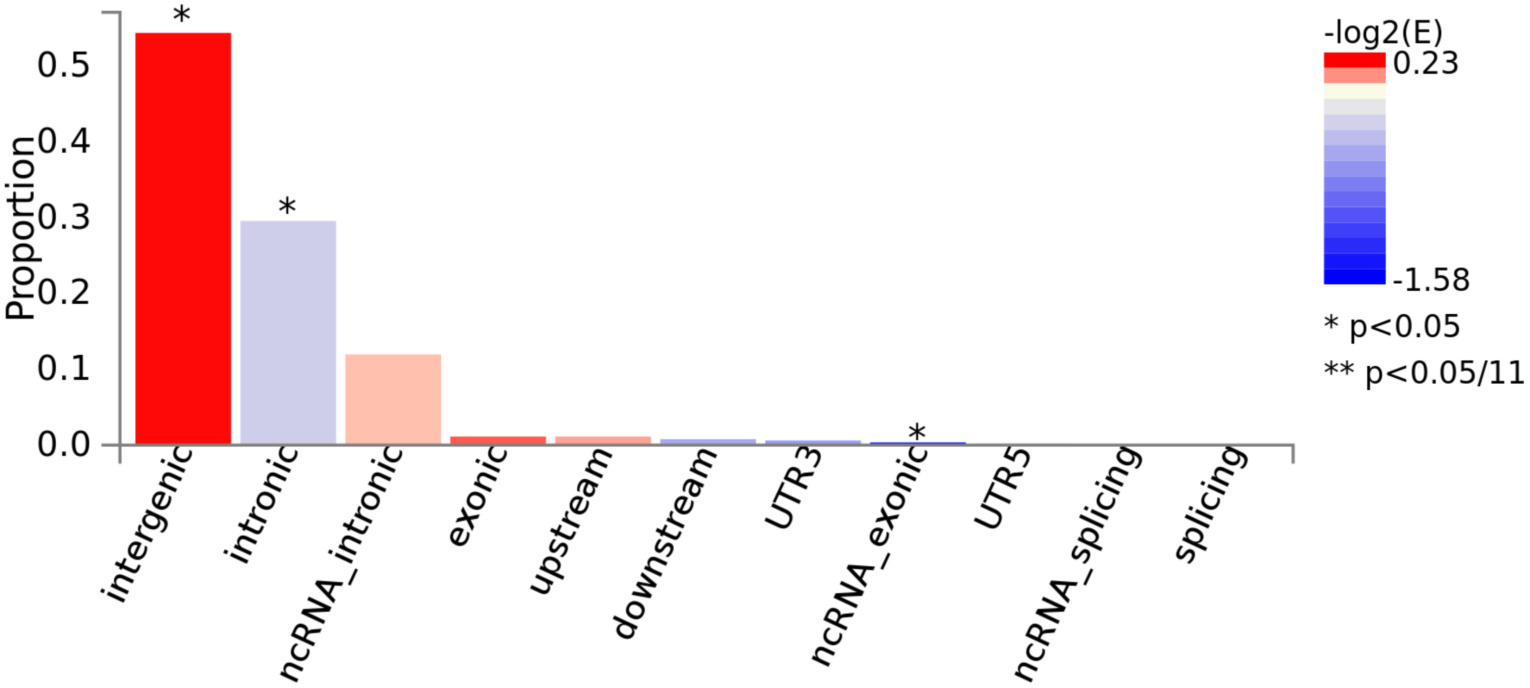

**Additional Supplementary Figure 4: Number of SNPs and Genes at each locus associated with AUD**

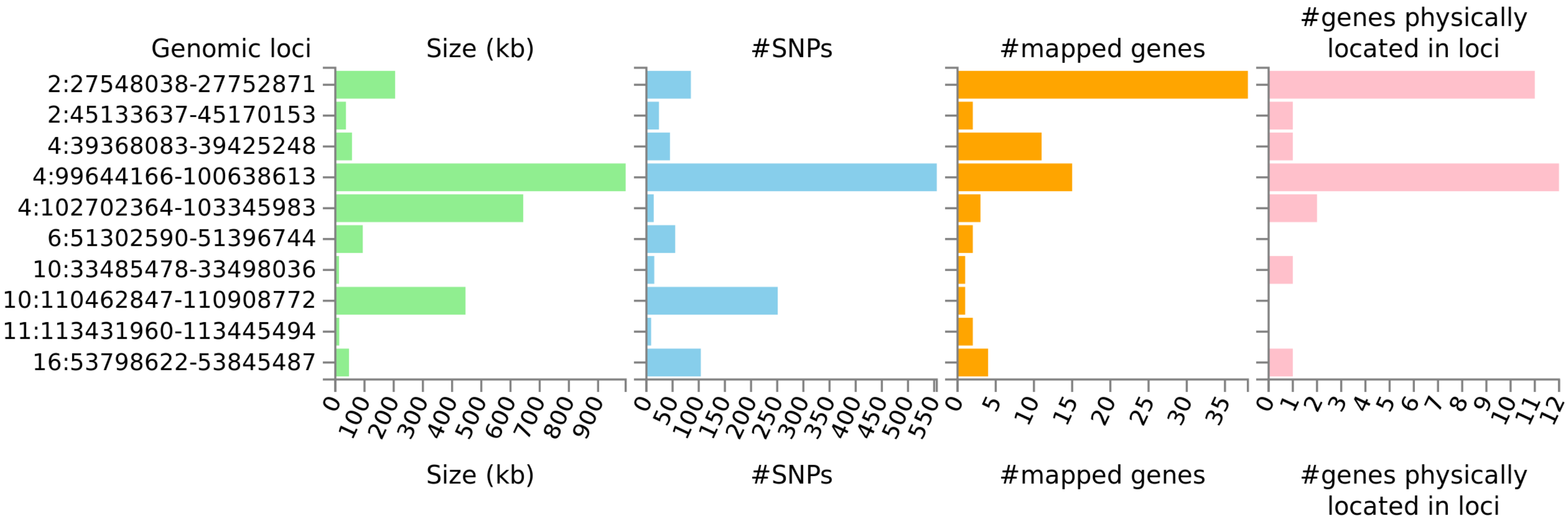

**Additional Supplementary Figure 5: Number of overlapping SNPs and genes shared between AUD and DPW GWAS analysis**

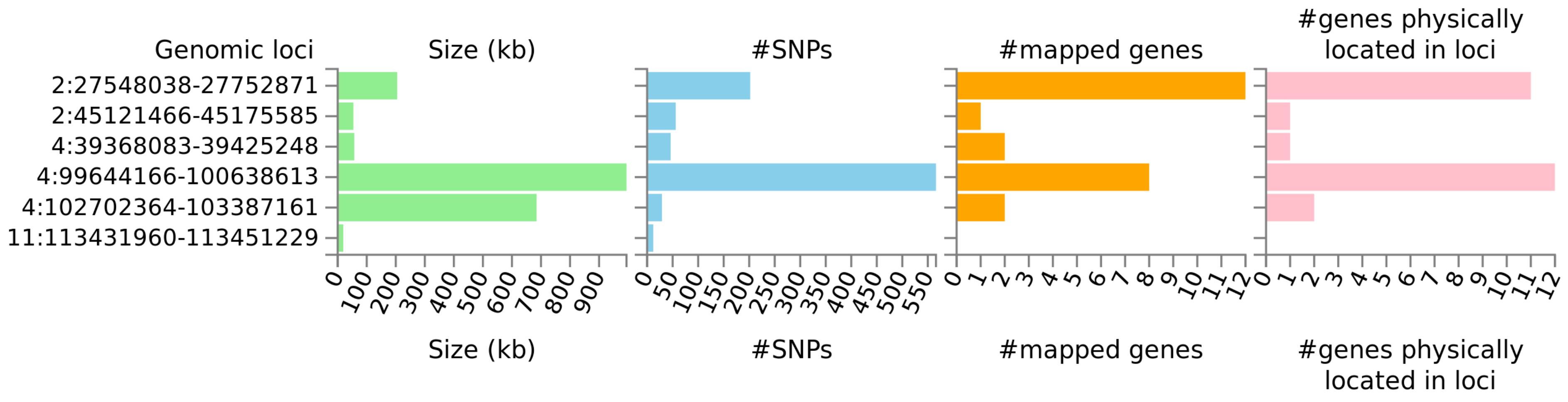

**Additional Supplementary Figure 6: Distribution of SNPs shared between AUD GWAS and DPW GWAS across the genome (Annotations and information from FUMA)**

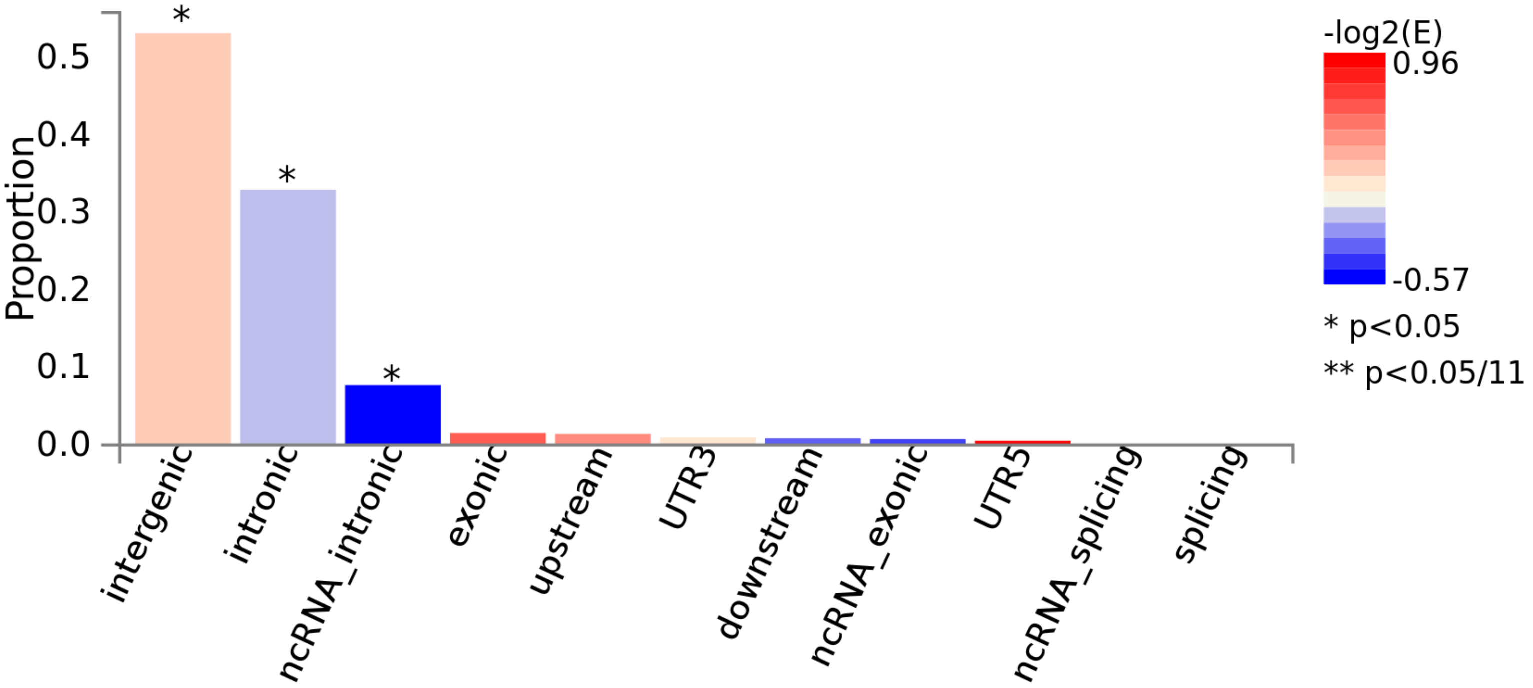

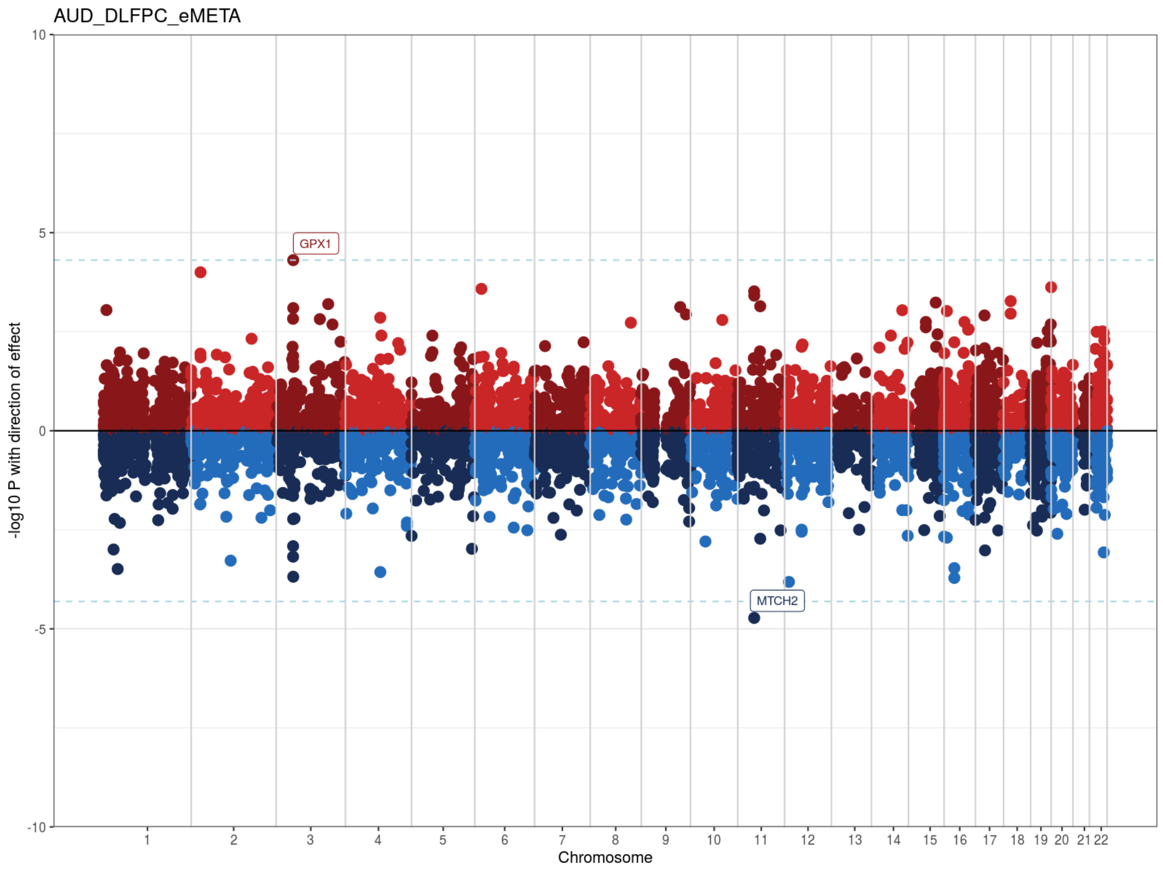


**Additional Supplementary Figure 7: Results of SMR based integration analysis of AUD GWAS with eQTL from adult brain.** X-axis represents the chromosomes and Y axis shows the direction of effect (Z scores) on gene expression/ methylation. Genes marked on the plots represent the genes nominated through threshold of co-localization (FDR < 20%) and/ or multiple levels of transcriptomic and epigenetic evidence.

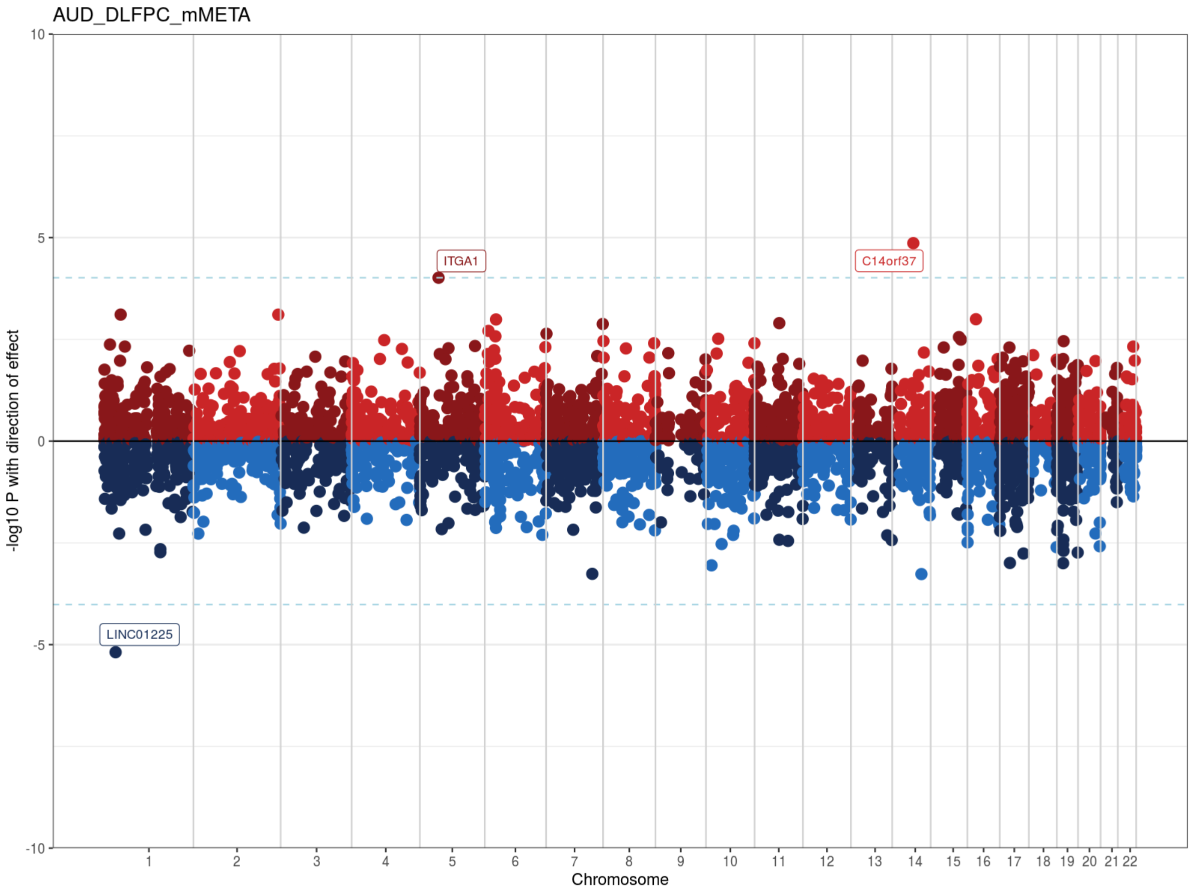


**Additional Supplementary Figure 8: Results of SMR based integration analysis of AUD GWAS with mQTL from adult brain.** X-axis represents the chromosomes and Y axis shows the direction of effect (Z scores) on gene expression/ methylation. Genes marked on the plots represent the genes nominated through threshold of co-localization (FDR < 20%) and/ or multiple levels of transcriptomic and epigenetic evidence.

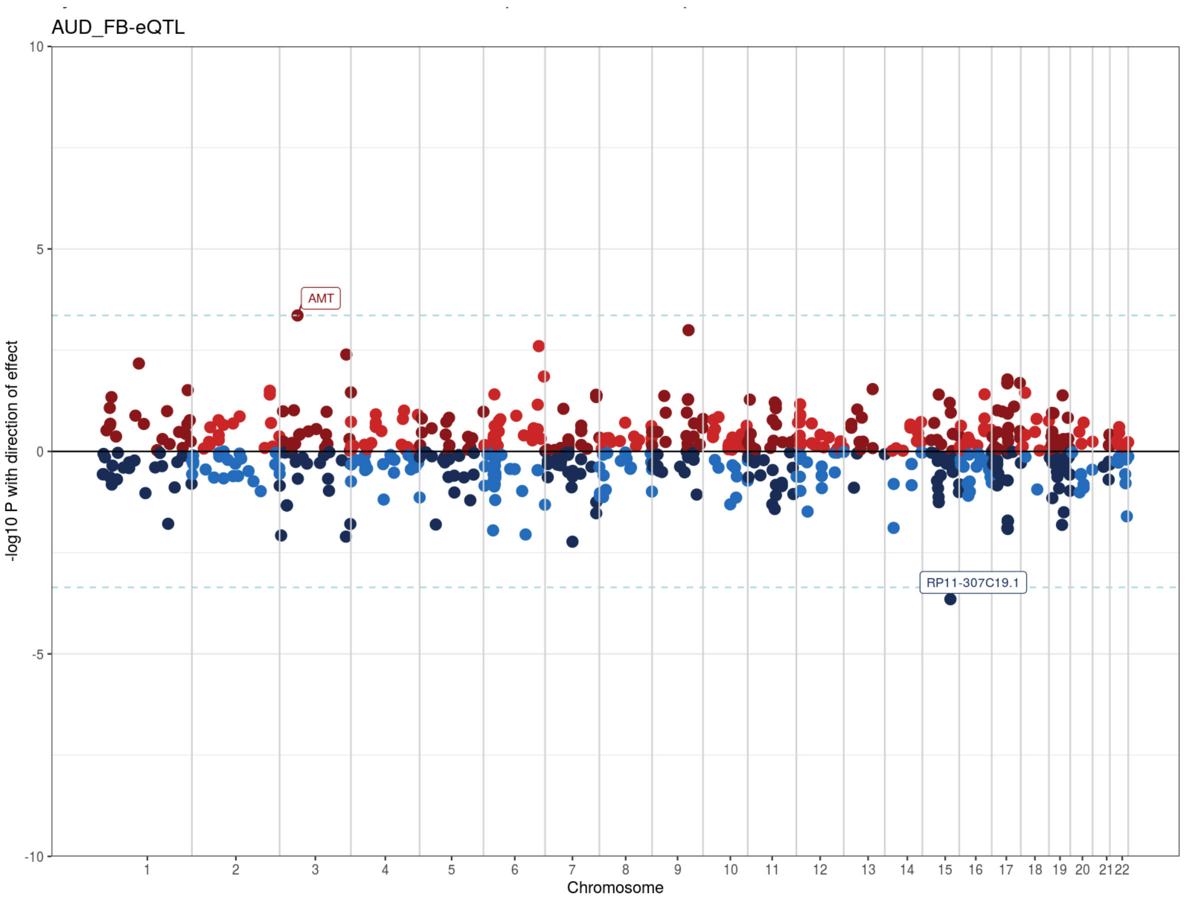


**Additional Supplementary Figure 9: Results of SMR based integration analysis of AUD GWAS with eQTL from fetal brain.** X-axis represents the chromosomes and Y axis shows the direction of effect (Z scores) on gene expression/ methylation. Genes marked on the plots represent the genes nominated through threshold of co-localization (FDR < 20%) and/ or multiple levels of transcriptomic and epigenetic evidence.

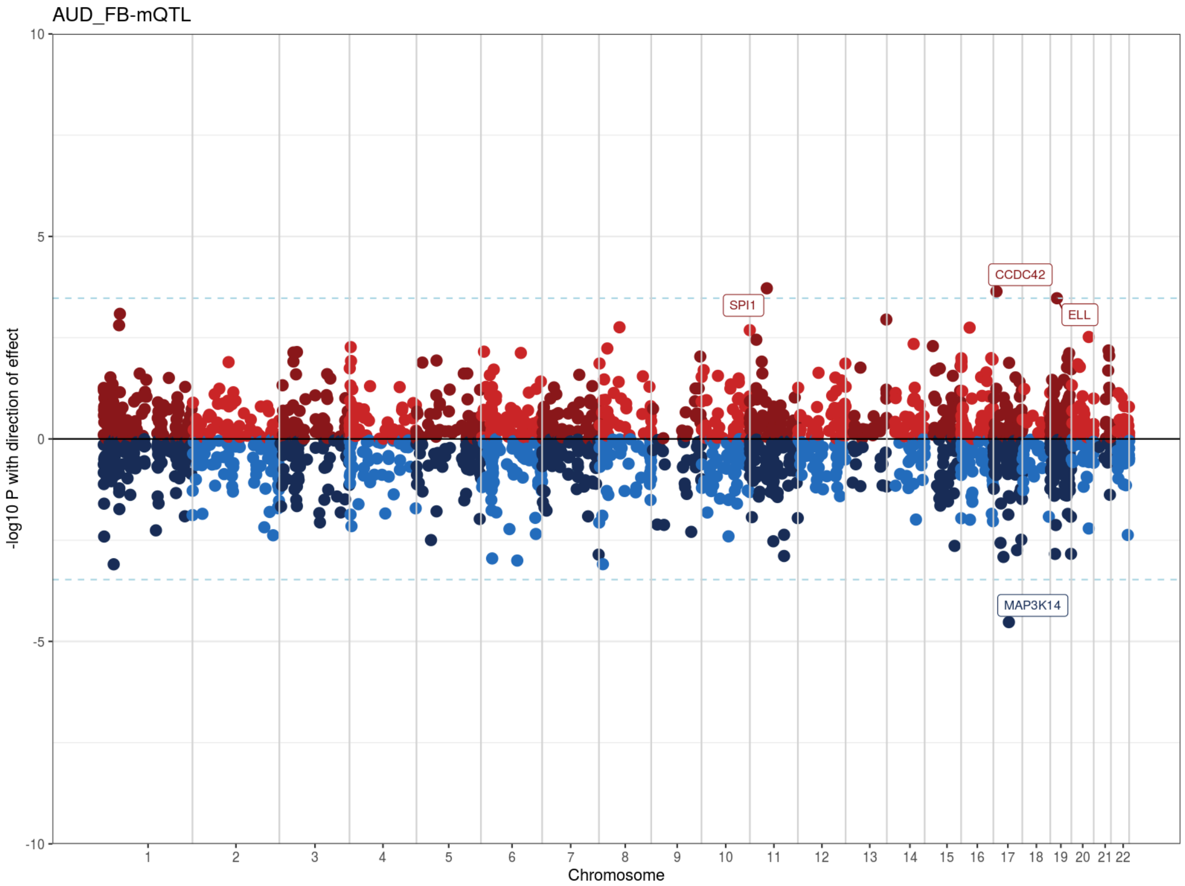


**Additional Supplementary Figure 10: Results of SMR based integration analysis of AUD GWAS with mQTL from fetal brain.** X-axis represents the chromosomes and Y axis shows the direction of effect (Z scores) on gene expression/ methylation. Genes marked on the plots represent the genes nominated through threshold of co-localization (FDR < 20%) and/ or multiple levels of transcriptomic and epigenetic evidence.

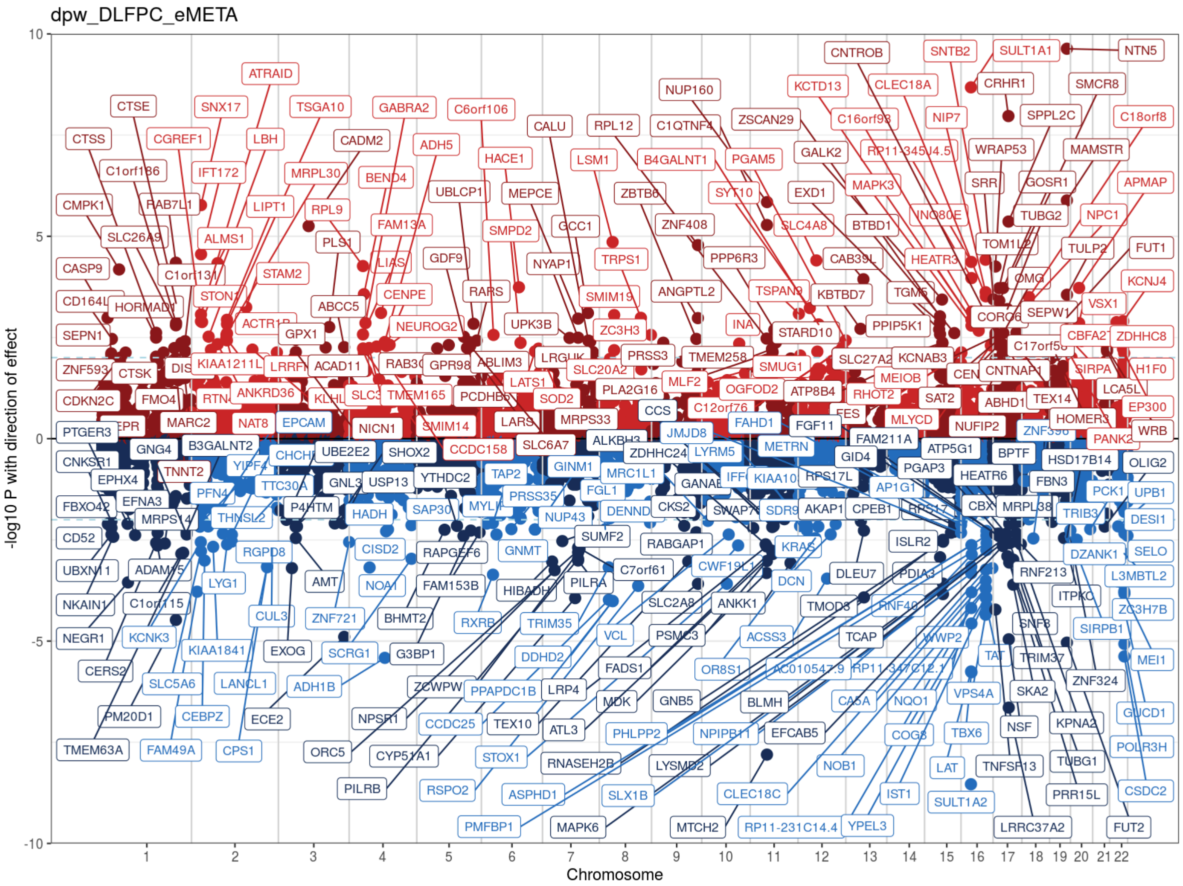


**Additional Supplementary Figure 11: Results of SMR based integration analysis of DPW GWAS with eQTL from adult brain.** X-axis represents the chromosomes and Y axis shows the direction of effect (Z scores) on gene expression/ methylation. Genes marked on the plots represent the genes nominated through threshold of co-localization (FDR < 20%) and/ or multiple levels of transcriptomic and epigenetic evidence.

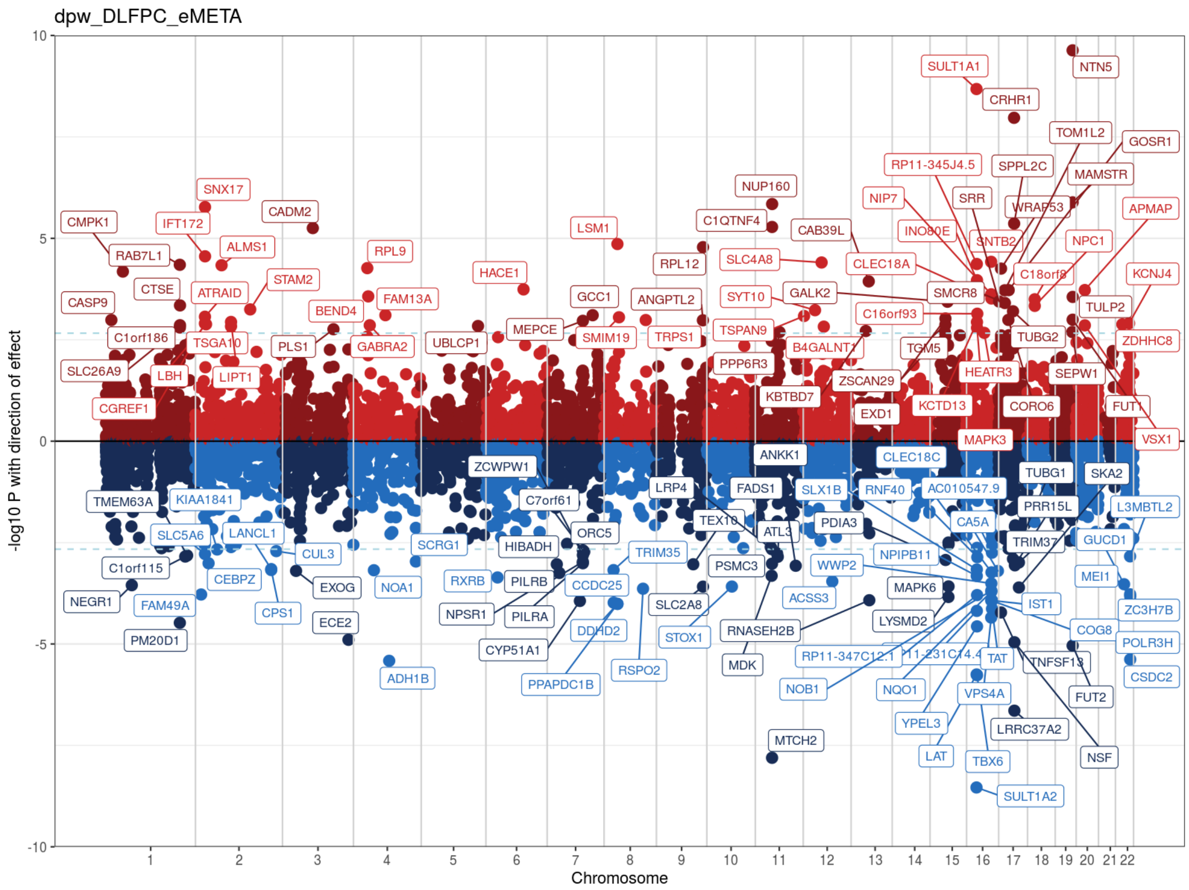


**Additional Supplementary Figure 12: Results of SMR based integration analysis of DPW GWAS with eQTL from adult brain.** X-axis represents the chromosomes and Y axis shows the direction of effect (Z scores) on gene expression/ methylation. Genes marked on the plots represent the genes nominated through threshold of co-localization (FDR < 10%) and/ or multiple levels of transcriptomic and epigenetic evidence.

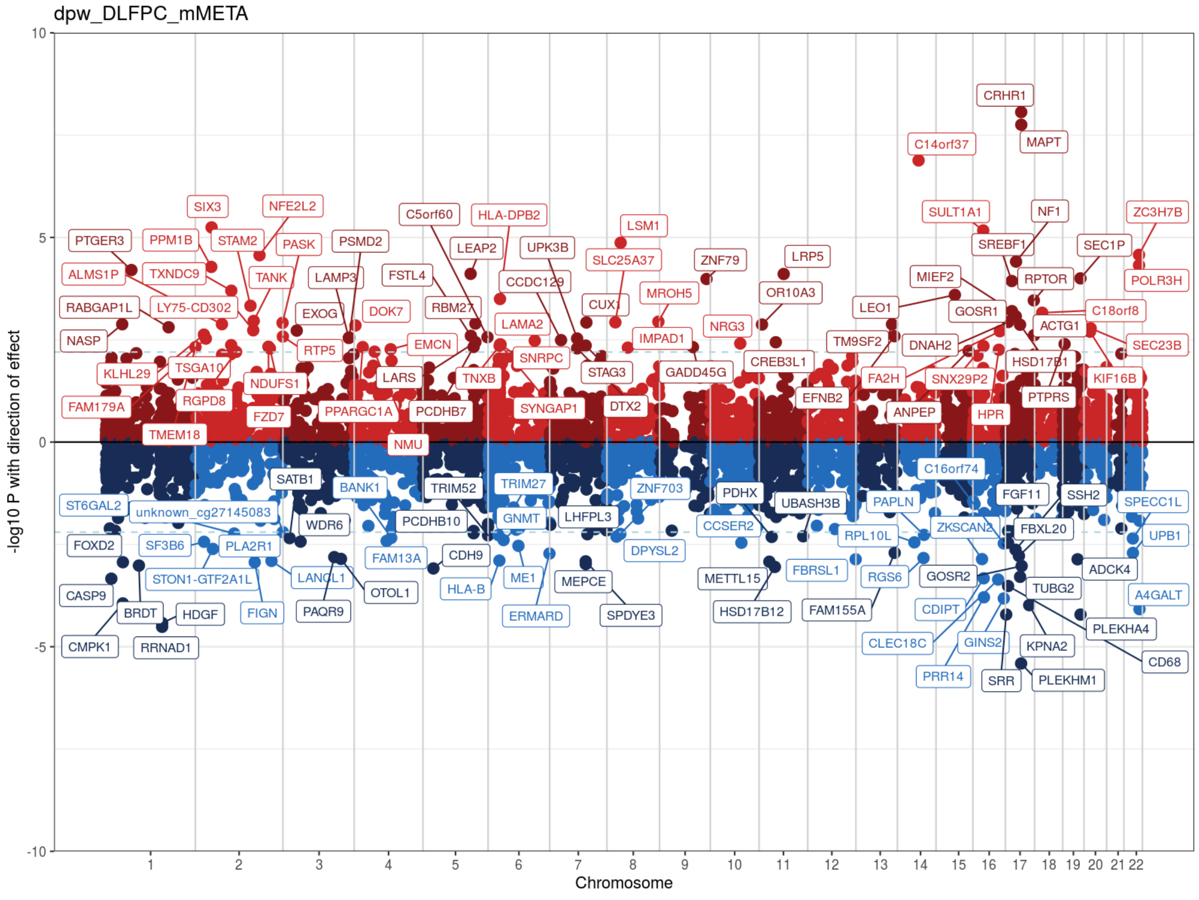


**Additional Supplementary Figure 13: Results of SMR based integration analysis of DPW GWAS with mQTL from adult brain.** X-axis represents the chromosomes and Y axis shows the direction of effect (Z scores) on gene expression/ methylation. Genes marked on the plots represent the genes nominated through threshold of co-localization (FDR < 20%) and/ or multiple levels of transcriptomic and epigenetic evidence.

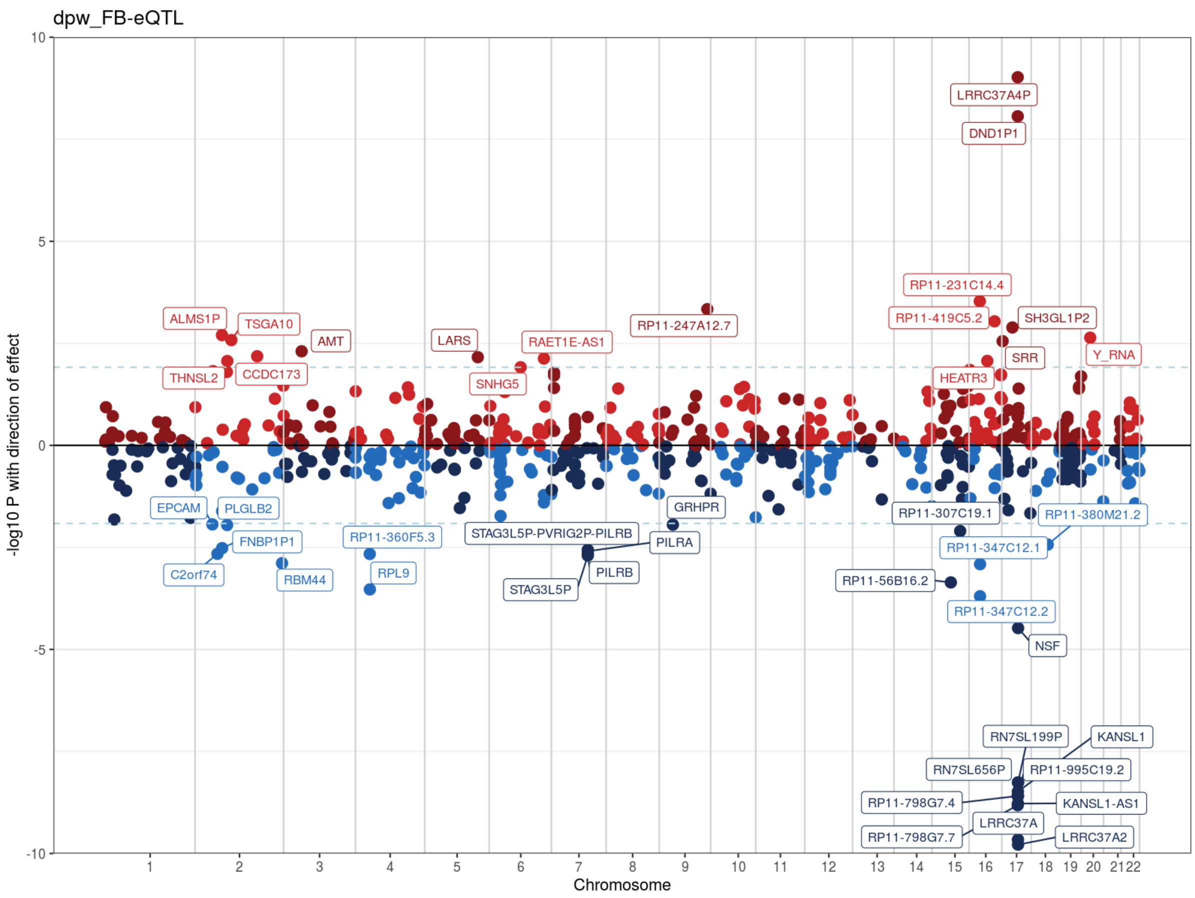


**Additional Supplementary Figure 14: Results of SMR based integration analysis of DPW GWAS with eQTL from fetal brain.** X-axis represents the chromosomes and Y axis shows the direction of effect (Z scores) on gene expression/ methylation. Genes marked on the plots represent the genes nominated through threshold of co-localization (FDR < 20%) and/ or multiple levels of transcriptomic and epigenetic evidence.

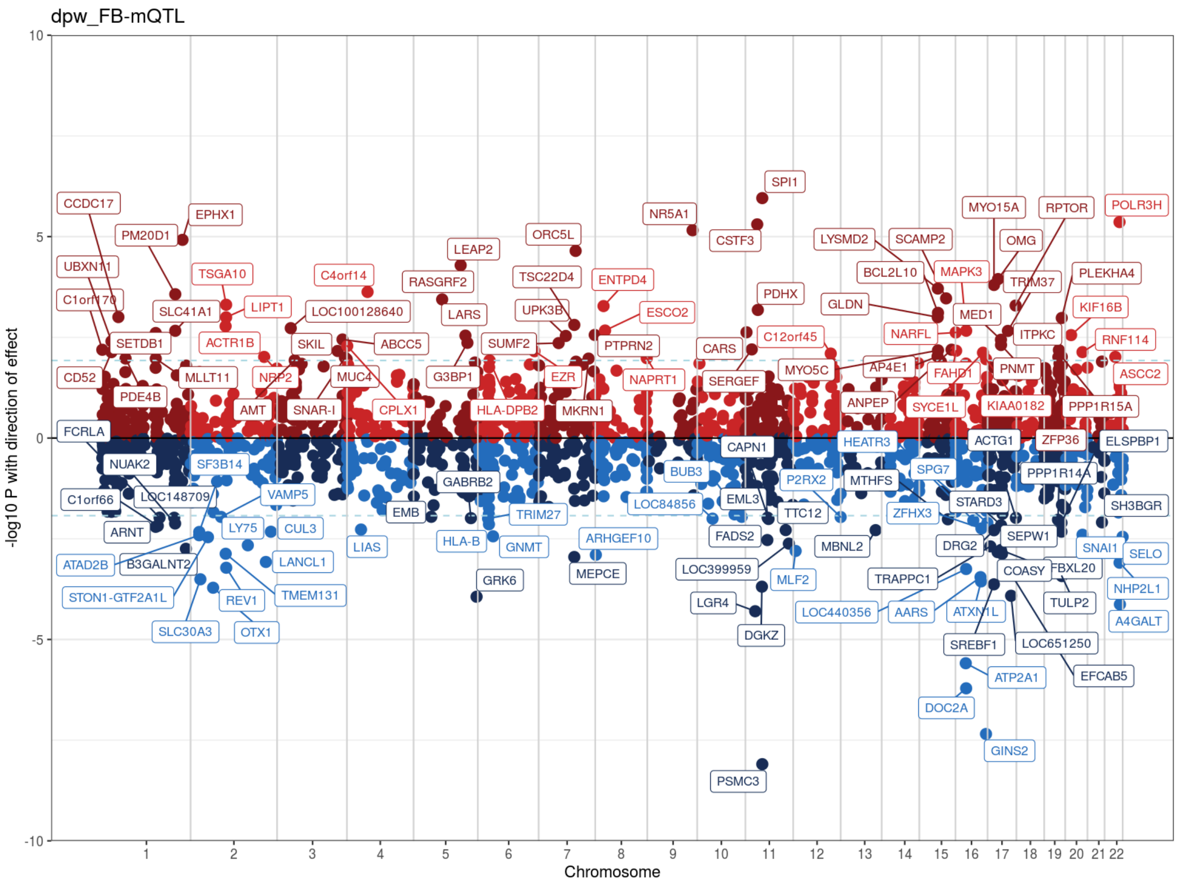


**Additional Supplementary Figure 15: Results of SMR based integration analysis of DPW GWAS with mQTL from fetal brain.** X-axis represents the chromosomes and Y axis shows the direction of effect (Z scores) on gene expression/ methylation. Genes marked on the plots represent the genes nominated through threshold of co-localization (FDR < 20%) and/ or multiple levels of transcriptomic and epigenetic evidence.
