## Supplementary table 4 for "Multi-omics integration analysis identifies novel genes for alcoholism with potential link to neurodegenerative diseases"

**Supp table 4: Results of pathway analysis with genes priortized in DPW integration analysis**

© 2000-2020 QIAGEN. All rights reserved.

| Ingenuity Canonical Pathways | -log(p-value) | Ratio | Molecules |
|---|---|---|---|
| TR/RXR Activation | 3.84 | 0.0952 | ATP2A1,EP300,ME1,MED1,PCK1,PPARGC1A,RXRB,SREBF1 |
| Lipoate Biosynthesis and Incorporation II | 3.48 | 1 | LIAS,LIPT1 |
| γ-linolenate Biosynthesis II (Animals) | 2.47 | 0.176 | FADS1,FADS2,SLC27A2 |
| Estrogen Biosynthesis | 2.19 | 0.0976 | CYP51A1,HSD17B1,HSD17B12,HSD17B14 |
| Sirtuin Signaling Pathway | 2.14 | 0.0412 | CPS1,GADD45G,MAPK6,MLYCD,NDUFS1,NFE2L2,NQO1,PCK1,PPARGC1A,RPTOR,SOD2,SREBF1 |
| Superoxide Radicals Degradation | 2.07 | 0.25 | NQO1,SOD2 |
| Glycine Betaine Degradation | 1.87 | 0.2 | BHMT2,SRR |
| Cardiac β-adrenergic Signaling | 1.83 | 0.0496 | AKAP11,ATP2A1,GNB5,GNG4,PDE4B,PPP1R14A,TULP2 |
| RhoGDI Signaling | 1.76 | 0.0444 | ACTG1,ARHGEF10,CDH9,EP300,EZR,GNB5,GNG4,RHOT2 |
| Lipoate Salvage and Modification | 1.74 | 1 | LIPT1 |
| 4-hydroxybenzoate Biosynthesis | 1.74 | 1 | TAT |
| 4-hydroxyphenylpyruvate Biosynthesis | 1.74 | 1 | TAT |
| Unfolded protein response | 1.73 | 0.0714 | CEBPZ,NFE2L2,PPP1R15A,SREBF1 |
| Phagosome Maturation | 1.69 | 0.0464 | CTSK,CTSS,GOSR1,GOSR2,HLA-B,TUBG1,TUBG2 |
| Oleate Biosynthesis II (Animals) | 1.65 | 0.154 | FADS1,FADS2 |
| NRF2-mediated Oxidative Stress Response | 1.65 | 0.0423 | ACTG1,CUL3,EP300,EPHX1,KRAS,NFE2L2,NQO1,SOD2 |
| Synaptogenesis Signaling Pathway | 1.55 | 0.0353 | CDH9,CNTNAP1,CPLX1,EFNA3,EFNB2,GOSR1,GOSR2,KRAS,MAPT,SYNGAP1,SYT10 |
| Regulation of Cellular Mechanics by Calpain | 1.52 | 0.0615 | CAPN1,EZR,KRAS,VCL |
| Antigen Presentation Pathway | 1.47 | 0.0769 | HLA-B,PDIA3,TAP2 |
| Formaldehyde Oxidation II (Glutathione-dep | 1.44 | 0.5 | ADH5 |
| Xenobiotic Metabolism Signaling | 1.42 | 0.0348 | ARNT,CUL3,EP300,KRAS,MED1,NFE2L2,NQO1,PPARGC1A,SULT1A1,SULT1A2 |
| Mechanisms of Viral Exit from Host Cells | 1.42 | 0.0732 | ACTG1,SNF8,VPS4A |
| Estrogen-Dependent Breast Cancer Signalir | 1.34 | 0.0541 | HSD17B1,HSD17B12,HSD17B14,KRAS |
| Aryl Hydrocarbon Receptor Signaling | 1.33 | 0.042 | ARNT,EP300,MED1,NFE2L2,NQO1,RXRB |
| CDK5 Signaling | 1.33 | 0.0463 | CAPN1,KRAS,MAPK6,MAPT,PPP1R14A |
| Role of Oct4 in Mammalian Embryonic Stem | 1.29 | 0.0652 | NR5A1,RXRB,WWP2 |
| Uracil Degradation II (Reductive) | 1.27 | 0.333 | UPB1 |
| Coenzyme A Biosynthesis | 1.27 | 0.333 | COASY |
| Methionine Salvage II (Mammalian) | 1.27 | 0.333 | BHMT2 |
| Thymine Degradation | 1.27 | 0.333 | UPB1 |
| L-serine Degradation | 1.27 | 0.333 | SRR |
| HIF1α Signaling | 1.26 | 0.0442 | ARNT,EP300,KRAS,MAPK6,SLC2A8 |
| Agrin Interactions at Neuromuscular Junctio | 1.25 | 0.0506 | ACTG1,KRAS,LAMA2,NRG3 |
| Polyamine Regulation in Colon Cancer | 1.22 | 0.0909 | KRAS,SAT2 |
| Pyrimidine Deoxyribonucleotides De Novo B | 1.22 | 0.0909 | CMPK1,PCK1 |
| Axonal Guidance Signaling | 1.22 | 0.0289 | ABLIM3,ADAM15,ECE2,EFNA3,EFNB2,FES,FZD7,GNB5,GNG4,KRAS,NRP2,PDIA3,RTN4,TUBG1 |
| Superpathway of D-myo-inositol (1,4,5)-trispl | 1.19 | 0.087 | BPNT2,ITPKC |
| LXR/RXR Activation | 1.15 | 0.0413 | CYP51A1,HADH,MYLIP,RXRB,SREBF1 |
| Spermine and Spermidine Degradation I | 1.15 | 0.25 | SAT2 |
| Catecholamine Biosynthesis | 1.15 | 0.25 | PNMT |
| Glutathione Redox Reactions II | 1.15 | 0.25 | PDIA3 |
| Acetate Conversion to Acetyl-CoA | 1.15 | 0.25 | ACSS3 |
| RhoA Signaling | 1.13 | 0.0407 | ACTG1,EZR,NRP2,PFN4,RAPGEF6 |
| Transcriptional Regulatory Network in Embry | 1.13 | 0.0556 | OTX1,SKIL,ZFHX3 |
| Myo-inositol Biosynthesis | 1.06 | 0.2 | BPNT2 |
| Galactose Degradation I (Leloir Pathway) | 1.06 | 0.2 | GALK2 |
| Tyrosine Degradation I | 1.06 | 0.2 | TAT |
| Nur77 Signaling in T Lymphocytes | 1.04 | 0.0508 | CASP9,EP300,HLA-B |
| FAK Signaling | 1.02 | 0.0421 | ACTG1,CAPN1,KRAS,VCL |
| Melatonin Degradation I | 1.02 | 0.05 | CYP51A1,SULT1A1,SULT1A2 |
| PCP pathway | 1.02 | 0.05 | FZD7,LGR4,PFN4 |
| Adipogenesis pathway | 1.01 | 0.0373 | FZD7,PPIP5K1,SAP30,SETDB1,SREBF1 |
| Phospholipase C Signaling | 1.01 | 0.0311 | ARHGEF10,EP300,GNB5,GNG4,KRAS,LAT,PPP1R14A,RHOT2 |
| Amyotrophic Lateral Sclerosis Signaling | 1 | 0.0412 | CAPN1,CASP9,CCS,GPX1 |
| Salvage Pathways of Pyrimidine Ribonucleo | 1 | 0.0412 | CMPK1,GRK6,MAPK6,PCK1 |
| p53 Signaling | 0.987 | 0.0408 | EP300,GADD45G,GNL3,MED1 |
| Dopamine Degradation | 0.987 | 0.0667 | SULT1A1,SULT1A2 |
| UVA-Induced MAPK Signaling | 0.987 | 0.0408 | CASP9,KRAS,PDIA3,SMPD2 |
| Urea Cycle | 0.983 | 0.167 | CPS1 |
| Glycine Cleavage Complex | 0.983 | 0.167 | AMT |
| Zymosterol Biosynthesis | 0.983 | 0.167 | CYP51A1 |
| Estrogen Receptor Signaling | 0.979 | 0.0365 | EP300,KRAS,MED1,PCK1,PPARGC1A |
| Apoptosis Signaling | 0.975 | 0.0404 | BCL2L10,CAPN1,CASP9,KRAS |
| VEGF Signaling | 0.975 | 0.0404 | ACTG1,ARNT,KRAS,VCL |
| Apelin Cardiomyocyte Signaling Pathway | 0.975 | 0.0404 | ARNT,ATP2A1,MAPK6,PDIA3 |
| G Protein Signaling Mediated by Tubby | 0.967 | 0.0645 | GNB5,GNG4 |
| D-myo-inositol (1,4,5,6)-Tetrakisphosphate E | 0.951 | 0.0357 | PGAM5,PPIP5K1,PPP1R14A,SIRPA,UBLCP1 |
| D-myo-inositol (3,4,5,6)-tetrakisphosphate B | 0.951 | 0.0357 | PGAM5,PPIP5K1,PPP1R14A,SIRPA,UBLCP1 |
| Glutathione-mediated Detoxification | 0.943 | 0.0625 | ANPEP,LANCL1 |
| Superpathway of Melatonin Degradation | 0.943 | 0.0462 | CYP51A1,SULT1A1,SULT1A2 |
| Ethanol Degradation II | 0.943 | 0.0625 | ACSS3,ADH5 |
| Fatty Acid β-oxidation I | 0.943 | 0.0625 | HADH,SLC27A2 |
| EIF2 Signaling | 0.939 | 0.0312 | KRAS,PPP1R15A,RPL12,RPL9,RPS17,SREBF1,TRIB3 |
| Calcium-induced T Lymphocyte Apoptosis | 0.928 | 0.0455 | ATP2A1,EP300,HLA-B |
| PPAR Signaling | 0.917 | 0.0385 | EP300,KRAS,MED1,PPARGC1A |
| Serotonin Degradation | 0.914 | 0.0448 | ADH5,SULT1A1,SULT1A2 |
| Cytotoxic T Lymphocyte-mediated Apoptosis | 0.9 | 0.0588 | CASP9,HLA-B |
| Remodeling of Epithelial Adherens Junction | 0.9 | 0.0441 | ACTG1,TUBG1,VCL |

| | | | |
|---|---|---|---|
| Endothelin-1 Signaling | 0.889 | 0.0319 | CASP9,ECE2,KRAS,MAPK6,PDIA3,PLA2R1 |
| Virus Entry via Endocytic Pathways | 0.883 | 0.0374 | ACTG1,AP1G1,HLA-B,KRAS |
| Noradrenaline and Adrenaline Degradation | 0.876 | 0.0571 | ADH5,PNMT |
| Inositol Pyrophosphates Biosynthesis | 0.866 | 0.125 | PPIP5K1 |
| Sphingomyelin Metabolism | 0.866 | 0.125 | SMPD2 |
| Huntington's Disease Signaling | 0.848 | 0.0295 | CAPN1,CASP9,EP300,GNB5,GNG4,GOSR1,GOSR2 |
| Ephrin B Signaling | 0.845 | 0.0417 | EFNB2,GNB5,GNG4 |
| Epithelial Adherens Junction Signaling | 0.842 | 0.0329 | ACTG1,KRAS,SNAI1,TUBG1,VCL |
| Non-Small Cell Lung Cancer Signaling | 0.833 | 0.0411 | CASP9,KRAS,RXRB |
| Cardiac Hypertrophy Signaling | 0.83 | 0.0292 | ELSPBP1,EP300,GNB5,GNG4,KRAS,PDIA3,RHOT2 |
| Superpathway of Inositol Phosphate Compo | 0.824 | 0.0305 | ITPKC,PGAM5,PPIP5K1,PPP1R14A,SIRPA,UBLCP1 |
| 3-phosphoinositide Degradation | 0.824 | 0.0325 | PGAM5,PPIP5K1,PPP1R14A,SIRPA,UBLCP1 |
| Thyroid Hormone Metabolism II (via Conjuga | 0.821 | 0.0526 | SULT1A1,SULT1A2 |
| Hypoxia Signaling in the Cardiovascular Sys | 0.821 | 0.0405 | ARNT,EP300,NQO1 |
| D-myo-inositol-5-phosphate Metabolism | 0.818 | 0.0323 | PGAM5,PPIP5K1,PPP1R14A,SIRPA,UBLCP1 |
| Folate Transformations I | 0.818 | 0.111 | MTHFS |
| Signaling by Rho Family GTPases | 0.804 | 0.0287 | ACTG1,ARHGEF10,CDH9,EZR,GNB5,GNG4,RHOT2 |
| Breast Cancer Regulation by Stathmin1 | 0.801 | 0.03 | ARHGEF10,GNB5,GNG4,KRAS,PPP1R14A,TUBG1 |
| Neuroprotective Role of THOP1 in Alzheime | 0.793 | 0.0345 | ECE2,HLA-B,MAPT,PRSS3 |
| Sphingosine-1-phosphate Signaling | 0.785 | 0.0342 | CASP9,PDIA3,RHOT2,SMPD2 |
| Antiproliferative Role of Somatostatin Recep | 0.783 | 0.039 | GNB5,GNG4,KRAS |
| Calcium Transport I | 0.775 | 0.1 | ATP2A1 |
| VDR/RXR Activation | 0.772 | 0.0385 | EP300,MED1,RXRB |
| Pyrimidine Ribonucleotides Interconversion | 0.767 | 0.0488 | CMPK1,PCK1 |
| Renal Cell Carcinoma Signaling | 0.75 | 0.0375 | ARNT,EP300,KRAS |
| Thrombin Signaling | 0.747 | 0.0288 | ARHGEF10,GNB5,GNG4,KRAS,PDIA3,RHOT2 |
| Colorectal Cancer Metastasis Signaling | 0.747 | 0.0277 | CASP9,FZD7,GNB5,GNG4,KRAS,PTGER3,RHOT2 |
| 3-phosphoinositide Biosynthesis | 0.747 | 0.0305 | PGAM5,PPIP5K1,PPP1R14A,SIRPA,UBLCP1 |
| γ-glutamyl Cycle | 0.738 | 0.0909 | ANPEP |
| Pyrimidine Ribonucleotides De Novo Biosynt | 0.735 | 0.0465 | CMPK1,PCK1 |
| tRNA Splicing | 0.735 | 0.0465 | PDE4B,TULP2 |
| Apelin Adipocyte Signaling Pathway | 0.728 | 0.0366 | GPX1,MAPK6,PPARGC1A |
| Protein Kinase A Signaling | 0.721 | 0.0251 | AKAP11,GNB5,GNG4,PDE4B,PDIA3,PPP1R14A,PTPRS,SIRPA,TULP2,UBASH3B |
| Tight Junction Signaling | 0.719 | 0.0298 | ACTG1,CSTF3,GOSR1,GOSR2,VCL |
| Integrin Signaling | 0.717 | 0.0282 | ACTG1,CAPN1,KRAS,PFN4,RHOT2,VCL |
| Gαi Signaling | 0.714 | 0.032 | GNB5,GNG4,KRAS,PTGER3 |
| Cleavage and Polyadenylation of Pre-mRNA | 0.703 | 0.0833 | CSTF3 |
| 14-3-3-mediated Signaling | 0.699 | 0.0315 | KRAS,MAPT,PDIA3,TUBG1 |
| P2Y Purigenic Receptor Signaling Pathway | 0.699 | 0.0315 | GNB5,GNG4,KRAS,PDIA3 |
| Germ Cell-Sertoli Cell Junction Signaling | 0.697 | 0.0292 | ACTG1,KRAS,RHOT2,TUBG1,VCL |
| Dermatan Sulfate Biosynthesis (Late Stages | 0.693 | 0.0435 | SULT1A1,SULT1A2 |
| Synaptic Long Term Potentiation | 0.682 | 0.031 | EP300,KRAS,PDIA3,PPP1R14A |
| Fatty Acid Activation | 0.674 | 0.0769 | SLC27A2 |
| Cholesterol Biosynthesis I | 0.674 | 0.0769 | CYP51A1 |
| Cholesterol Biosynthesis II (via 24,25-dihydr | 0.674 | 0.0769 | CYP51A1 |
| Cholesterol Biosynthesis III (via Desmostero | 0.674 | 0.0769 | CYP51A1 |
| Ceramide Signaling | 0.668 | 0.0341 | CNKSR1,KRAS,SMPD2 |
| Chondroitin Sulfate Biosynthesis (Late Stag | 0.664 | 0.0417 | SULT1A1,SULT1A2 |
| CTLA4 Signaling in Cytotoxic T Lymphocyte | 0.658 | 0.0337 | AP1G1,HLA-B,LAT |
| Cell Cycle: G2/M DNA Damage Checkpoint I | 0.652 | 0.0408 | CKS2,EP300 |
| Phenylalanine Degradation IV (Mammalian, | 0.646 | 0.0714 | SLC27A2 |
| Colanic Acid Building Blocks Biosynthesis | 0.646 | 0.0714 | GALK2 |
| Amyloid Processing | 0.638 | 0.04 | CAPN1,MAPT |
| Ephrin Receptor Signaling | 0.638 | 0.0278 | EFNA3,EFNB2,GNB5,GNG4,KRAS |
| Role of NFAT in Regulation of the Immune I | 0.633 | 0.0276 | GNB5,GNG4,HLA-B,KRAS,LAT |
| Senescence Pathway | 0.629 | 0.0255 | CAPN1,EP300,GADD45G,KRAS,MAPK6,PDHX,SOD2 |
| Thyroid Cancer Signaling | 0.625 | 0.0392 | KRAS,RXRB |
| UVC-Induced MAPK Signaling | 0.625 | 0.0392 | KRAS,SMPD2 |
| Androgen Biosynthesis | 0.62 | 0.0667 | HSD17B14 |
| Superpathway of Citrulline Metabolism | 0.62 | 0.0667 | CPS1 |
| α-Adrenergic Signaling | 0.604 | 0.0316 | GNB5,GNG4,KRAS |
| Lymphotoxin β Receptor Signaling | 0.602 | 0.0377 | CASP9,EP300 |
| Hereditary Breast Cancer Signaling | 0.602 | 0.0286 | EP300,GADD45G,KRAS,TUBG1 |
| Neuregulin Signaling | 0.595 | 0.0312 | DCN,KRAS,NRG3 |
| Granzyme B Signaling | 0.595 | 0.0625 | CASP9 |
| Glutaryl-CoA Degradation | 0.595 | 0.0625 | HADH |
| Parkinson's Signaling | 0.595 | 0.0625 | CASP9 |
| Systemic Lupus Erythematosus In T Cell Sig | 0.583 | 0.024 | CASP9,EZR,HLA-B,KRAS,LAT,LEPR,RHOT2,RPTOR |
| Endocannabinoid Cancer Inhibition Pathway | 0.582 | 0.028 | CASP9,RPTOR,SMPD2,TRIB3 |
| PPARα/RXRα Activation | 0.578 | 0.0263 | EP300,KRAS,MED1,PDIA3,PPARGC1A |
| RAN Signaling | 0.572 | 0.0588 | KPNA2 |
| Mitochondrial L-carnitine Shuttle Pathway | 0.572 | 0.0588 | SLC27A2 |
| D-myo-inositol (1,4,5)-trisphosphate Degrada | 0.572 | 0.0588 | BPNT2 |
| Chondroitin Sulfate Biosynthesis | 0.567 | 0.0357 | SULT1A1,SULT1A2 |
| RAR Activation | 0.562 | 0.0259 | EP300,MED1,PPARGC1A,RXRB,SDR9C7 |
| 1D-myo-inositol Hexakisphosphate Biosynth | 0.551 | 0.0556 | ITPKC |
| Valine Degradation I | 0.551 | 0.0556 | HIBADH |
| D-myo-inositol (1,3,4)-trisphosphate Biosynth | 0.551 | 0.0556 | ITPKC |
| Adrenomedullin signaling pathway | 0.541 | 0.0254 | ARNT,KRAS,MAPK6,PDIA3,RXRB |
| Relaxin Signaling | 0.536 | 0.0267 | GNB5,GNG4,PDE4B,TULP2 |
| Dermatan Sulfate Biosynthesis | 0.536 | 0.0339 | SULT1A1,SULT1A2 |

| | | | |
|---|---|---|---|
| Granzyme A Signaling | 0.532 | 0.0526 | EP300 |
| GADD45 Signaling | 0.532 | 0.0526 | GADD45G |
| Oxidative Ethanol Degradation III | 0.532 | 0.0526 | ACSS3 |
| Apelin Muscle Signaling Pathway | 0.532 | 0.0526 | PPARGC1A |
| Semaphorin Signaling in Neurons | 0.526 | 0.0333 | FES,RHOT2 |
| Endometrial Cancer Signaling | 0.526 | 0.0333 | CASP9,KRAS |
| Retinoic acid Mediated Apoptosis Signaling | 0.526 | 0.0333 | CASP9,RXRB |
| PD-1, PD-L1 cancer immunotherapy pathwa | 0.519 | 0.0283 | HLA-B,LAT,LATS1 |
| Autophagy | 0.517 | 0.0328 | CTSK,CTSS |
| Gαs Signaling | 0.511 | 0.028 | CRHR1,GNB5,GNG4 |
| Wnt/Ca+ pathway | 0.507 | 0.0323 | FZD7,PDIA3 |
| Paxillin Signaling | 0.504 | 0.0278 | ACTG1,KRAS,VCL |
| Phospholipases | 0.498 | 0.0317 | PDIA3,PLA2R1 |
| Antioxidant Action of Vitamin C | 0.498 | 0.0275 | PDIA3,PLA2R1,SLC2A8 |
| Putrescine Degradation III | 0.495 | 0.0476 | SAT2 |
| Endoplasmic Reticulum Stress Pathway | 0.495 | 0.0476 | CASP9 |
| CREB Signaling in Neurons | 0.489 | 0.0242 | EP300,GNB5,GNG4,KRAS,PDIA3 |
| iCOS-iCOSL Signaling in T Helper Cells | 0.484 | 0.027 | HLA-B,LAT,PLEKHA4 |
| Myc Mediated Apoptosis Signaling | 0.48 | 0.0308 | CASP9,KRAS |
| ErbB2-ErbB3 Signaling | 0.48 | 0.0308 | KRAS,NRG3 |
| Pyridoxal 5'-phosphate Salvage Pathway | 0.48 | 0.0308 | GRK6,MAPK6 |
| Osteoarthritis Pathway | 0.471 | 0.0237 | C1QTNF4,CASP9,DCN,FZD7,PPARGC1A |
| Eicosanoid Signaling | 0.471 | 0.0303 | PLA2R1,PTGER3 |
| AMPK Signaling | 0.466 | 0.0236 | EP300,MLYCD,PPARGC1A,PPM1B,RPTOR |
| ErbB4 Signaling | 0.463 | 0.0299 | KRAS,NRG3 |
| Tryptophan Degradation III (Eukaryotic) | 0.463 | 0.0435 | HADH |
| Ethanol Degradation IV | 0.463 | 0.0435 | ACSS3 |
| Tec Kinase Signaling | 0.458 | 0.0244 | ACTG1,GNB5,GNG4,RHOT2 |
| Endocannabinoid Developing Neuron Pathv | 0.458 | 0.0261 | KRAS,MAPK6,RPTOR |
| Role of NFAT in Cardiac Hypertrophy | 0.457 | 0.0234 | EP300,GNB5,GNG4,KRAS,PDIA3 |
| Glioblastoma Multiforme Signaling | 0.453 | 0.0242 | FZD7,KRAS,PDIA3,RHOT2 |
| fMLP Signaling in Neutrophils | 0.452 | 0.0259 | GNB5,GNG4,KRAS |
| Tumoricidal Function of Hepatic Natural Kille | 0.449 | 0.0417 | CASP9 |
| Glutathione Redox Reactions I | 0.449 | 0.0417 | GPX1 |
| Role of JAK1 and JAK3 in γc Cytokine Signa | 0.446 | 0.029 | FES,KRAS |
| SPINK1 General Cancer Pathway | 0.446 | 0.029 | KRAS,PRSS3 |
| CXCR4 Signaling | 0.444 | 0.024 | GNB5,GNG4,KRAS,RHOT2 |
| Actin Cytoskeleton Signaling | 0.44 | 0.0229 | ACTG1,EZR,KRAS,SSH2,VCL |
| Bupropion Degradation | 0.434 | 0.04 | CYP51A1 |
| Small Cell Lung Cancer Signaling | 0.431 | 0.0282 | CASP9,RXRB |
| Heparan Sulfate Biosynthesis (Late Stages) | 0.431 | 0.0282 | SULT1A1,SULT1A2 |
| G-Protein Coupled Receptor Signaling | 0.428 | 0.0221 | CRHR1,KRAS,PDE4B,PTGER3,SYNGAP1,TULP2 |
| Mitochondrial Dysfunction | 0.424 | 0.0234 | CASP9,NDUFS1,RHOT2,SOD2 |
| Actin Nucleation by ARP-WASP Complex | 0.424 | 0.0278 | KRAS,RHOT2 |
| Lipid Antigen Presentation by CD1 | 0.421 | 0.0385 | PDIA3 |
| Gluconeogenesis I | 0.421 | 0.0385 | ME1 |
| G Beta Gamma Signaling | 0.418 | 0.0246 | GNB5,GNG4,KRAS |
| Inhibition of ARE-Mediated mRNA Degradat | 0.418 | 0.0246 | MAPK6,TNFSF13,ZFP36 |
| Caveolar-mediated Endocytosis Signaling | 0.416 | 0.0274 | ACTG1,HLA-B |
| LPS/IL-1 Mediated Inhibition of RXR Functic | 0.415 | 0.0223 | PPARGC1A,SLC27A2,SREBF1,SULT1A1,SULT1A2 |
| Leptin Signaling in Obesity | 0.409 | 0.027 | LEPR,PDIA3 |
| Glioma Invasiveness Signaling | 0.409 | 0.027 | KRAS,RHOT2 |
| CCR3 Signaling in Eosinophils | 0.407 | 0.0242 | GNB5,GNG4,KRAS |
| Angiopoietin Signaling | 0.402 | 0.0267 | CASP9,KRAS |
| cAMP-mediated signaling | 0.399 | 0.0219 | AKAP11,CRHR1,PDE4B,PTGER3,TULP2 |
| Systemic Lupus Erythematosus Signaling | 0.395 | 0.0218 | HLA-B,KRAS,LAT,LSM1,SNRPC |
| GDNF Family Ligand-Receptor Interactions | 0.395 | 0.0263 | DOK7,KRAS |
| Superpathway of Cholesterol Biosynthesis | 0.384 | 0.0345 | CYP51A1 |
| p70S6K Signaling | 0.381 | 0.0233 | KRAS,MAPT,PDIA3 |
| Heparan Sulfate Biosynthesis | 0.381 | 0.0256 | SULT1A1,SULT1A2 |
| Molecular Mechanisms of Cancer | 0.38 | 0.0205 | ARHGEF10,CASP9,EP300,FZD7,KRAS,RASGRF2,RHOT2,SYNGAP1 |
| Acetone Degradation I (to Methylglyoxal) | 0.374 | 0.0333 | CYP51A1 |
| Dendritic Cell Maturation | 0.371 | 0.0219 | HLA-B,LEPR,LY75,PDIA3 |
| Cellular Effects of Sildenafil (Viagra) | 0.371 | 0.0229 | ACTG1,PDE4B,PDIA3 |
| Role of MAPK Signaling in the Pathogenesi | 0.369 | 0.025 | KRAS,PLA2R1 |
| Prolactin Signaling | 0.363 | 0.0247 | EP300,KRAS |
| Sertoli Cell-Sertoli Cell Junction Signaling | 0.363 | 0.0216 | ACTG1,KRAS,TUBG1,VCL |
| PEDF Signaling | 0.357 | 0.0244 | KRAS,SOD2 |
| Production of Nitric Oxide and Reactive Oxy | 0.351 | 0.0213 | PPP1R14A,RHOT2,SIRPA,SPI1 |
| Synaptic Long Term Depression | 0.347 | 0.0212 | CRHR1,KRAS,PDIA3,PLA2R1 |
| Androgen Signaling | 0.347 | 0.0221 | EP300,GNB5,GNG4 |
| HER-2 Signaling in Breast Cancer | 0.344 | 0.0238 | CASP9,KRAS |
| VEGF Family Ligand-Receptor Interactions | 0.344 | 0.0238 | KRAS,NRP2 |
| ILK Signaling | 0.343 | 0.0211 | ACTG1,RHOT2,SNAI1,VCL |
| Retinoate Biosynthesis I | 0.343 | 0.0303 | SDR9C7 |
| PI3K Signaling in B Lymphocytes | 0.338 | 0.0217 | KRAS,PDIA3,PLEKHA4 |
| HIPPO signaling | 0.338 | 0.0235 | LATS1,PPP1R14A |
| IL-4 Signaling | 0.338 | 0.0235 | HLA-B,KRAS |
| DNA Methylation and Transcriptional Repres | 0.333 | 0.0294 | SAP30 |
| Role of JAK2 in Hormone-like Cytokine Sign | 0.333 | 0.0294 | SIRPA |
| Insulin Receptor Signaling | 0.333 | 0.0216 | KRAS,PPP1R14A,RPTOR |

| | | | |
|---|---|---|---|
| Neuroinflammation Signaling Pathway | 0.333 | 0.02 | GABRB2,HLA-B,MAPK6,MAPT,NFE2L2,SOD2 |
| TWEAK Signaling | 0.324 | 0.0286 | CASP9 |
| Acute Myeloid Leukemia Signaling | 0.316 | 0.0225 | KRAS,SPI1 |
| Crosstalk between Dendritic Cells and Natur | 0.316 | 0.0225 | ACTG1,HLA-B |
| B Cell Development | 0.316 | 0.0278 | HLA-B |
| Regulation of IL-2 Expression in Activated a | 0.316 | 0.0225 | KRAS,LAT |
| Gap Junction Signaling | 0.313 | 0.0202 | ACTG1,KRAS,PDIA3,TUBG1 |
| Altered T Cell and B Cell Signaling in Rheum | 0.312 | 0.0222 | HLA-B,TNFSF13 |
| Superpathway of Methionine Degradation | 0.307 | 0.027 | BHMT2 |
| Notch Signaling | 0.307 | 0.027 | DTX2 |
| IL-8 Signaling | 0.306 | 0.02 | GNB5,GNG4,KRAS,RHOT2 |
| Prostate Cancer Signaling | 0.306 | 0.022 | CASP9,KRAS |
| IL-1 Signaling | 0.306 | 0.022 | GNB5,GNG4 |
| Death Receptor Signaling | 0.306 | 0.022 | ACTG1,CASP9 |
| Docosahexaenoic Acid (DHA) Signaling | 0.299 | 0.0263 | CASP9 |
| April Mediated Signaling | 0.292 | 0.0256 | TNFSF13 |
| Fcγ Receptor-mediated Phagocytosis in Mac | 0.291 | 0.0213 | ACTG1,EZR |
| CCR5 Signaling in Macrophages | 0.291 | 0.0213 | GNB5,GNG4 |
| Melanocyte Development and Pigmentation | 0.291 | 0.0213 | EP300,KRAS |
| ErbB Signaling | 0.291 | 0.0213 | KRAS,NRG3 |
| Communication between Innate and Adaptiv | 0.281 | 0.0208 | HLA-B,TNFSF13 |
| TGF-β Signaling | 0.281 | 0.0208 | EP300,KRAS |
| ATM Signaling | 0.277 | 0.0206 | CBX1,GADD45G |
| Role of PKR in Interferon Induction and Ant | 0.277 | 0.0244 | CASP9 |
| nNOS Signaling in Skeletal Muscle Cells | 0.277 | 0.0244 | SNTB2 |
| mTOR Signaling | 0.274 | 0.019 | KRAS,RHOT2,RPS17,RPTOR |
| Gustation Pathway | 0.274 | 0.0195 | P2RX2,PDE4B,TULP2 |
| PKCθ Signaling in T Lymphocytes | 0.27 | 0.0194 | HLA-B,KRAS,LAT |
| Retinol Biosynthesis | 0.27 | 0.0238 | DDHD2 |
| Gαq Signaling | 0.264 | 0.0191 | GNB5,GNG4,RHOT2 |
| Oncostatin M Signaling | 0.263 | 0.0233 | KRAS |
| BAG2 Signaling Pathway | 0.263 | 0.0233 | MAPT |
| Role of RIG1-like Receptors in Antiviral Inna | 0.256 | 0.0227 | EP300 |
| HOTAIR Regulatory Pathway | 0.256 | 0.0189 | EP300,PCDHB5,SETDB1 |
| Mouse Embryonic Stem Cell Pluripotency | 0.251 | 0.0194 | FZD7,KRAS |
| Sumoylation Pathway | 0.251 | 0.0194 | EP300,RHOT2 |
| IGF-1 Signaling | 0.247 | 0.0192 | CASP9,KRAS |
| Stearate Biosynthesis I (Animals) | 0.244 | 0.0217 | SLC27A2 |
| Dopamine-DARPP32 Feedback in cAMP Sig | 0.243 | 0.0184 | ATP2A1,PDIA3,PPP1R14A |
| T Cell Receptor Signaling | 0.243 | 0.019 | KRAS,LAT |
| nNOS Signaling in Neurons | 0.238 | 0.0213 | CAPN1 |
| Ephrin A Signaling | 0.238 | 0.0213 | EFNA3 |
| Triacylglycerol Degradation | 0.238 | 0.0213 | DDHD2 |
| HMGB1 Signaling | 0.237 | 0.0182 | KRAS,RHOT2,TNFSF13 |
| Graft-versus-Host Disease Signaling | 0.232 | 0.0208 | HLA-B |
| Pancreatic Adenocarcinoma Signaling | 0.228 | 0.0183 | CASP9,KRAS |
| Autoimmune Thyroid Disease Signaling | 0.227 | 0.0204 | HLA-B |
| Melanoma Signaling | 0.221 | 0.02 | KRAS |
| Primary Immunodeficiency Signaling | 0.221 | 0.02 | TAP2 |
| TNFR1 Signaling | 0.221 | 0.02 | CASP9 |
| CD27 Signaling in Lymphocytes | 0.206 | 0.0189 | CASP9 |
| Phototransduction Pathway | 0.206 | 0.0189 | GNB5 |
| Glucocorticoid Receptor Signaling | 0 | 0.0149 | EP300,KRAS,MED1,PCK1,TAT |
| Natural Killer Cell Signaling | 0 | 0.0168 | KRAS,LAT |
| Leukocyte Extravasation Signaling | 0 | 0.0152 | ACTG1,EZR,VCL |
| Fc Epsilon RI Signaling | 0 | 0.0171 | KRAS,LAT |
| Acute Phase Response Signaling | 0 | 0.0112 | KRAS,SOD2 |
| Hepatic Cholestasis | 0 | 0.0109 | SREBF1,TNFSF13 |
| Hepatic Fibrosis / Hepatic Stellate Cell Activa | 0 | 0.00538 | LEPR |
| FXR/RXR Activation | 0 | 0.0159 | PPARGC1A,SREBF1 |
| PXR/RXR Activation | 0 | 0.0154 | PPARGC1A |
| Regulation of Actin-based Motility by Rho | 0 | 0.0106 | RHOT2 |
| Erythropoietin Signaling | 0 | 0.0132 | KRAS |
| Clathrin-mediated Endocytosis Signaling | 0 | 0.0104 | ACTG1,AP1G1 |
| IL-12 Signaling and Production in Macropha | 0 | 0.0152 | EP300,SPI1 |
| Role of Pattern Recognition Receptors in Re | 0 | 0.00649 | TNFSF13 |
| FcγRIIB Signaling in B Lymphocytes | 0 | 0.0133 | KRAS |
| LPS-stimulated MAPK Signaling | 0 | 0.0122 | KRAS |
| NF-κB Activation by Viruses | 0 | 0.0122 | KRAS |
| IL-3 Signaling | 0 | 0.0127 | KRAS |
| IL-17 Signaling | 0 | 0.0125 | KRAS |
| Thrombopoietin Signaling | 0 | 0.0159 | KRAS |
| Induction of Apoptosis by HIV1 | 0 | 0.0164 | CASP9 |
| IL-15 Production | 0 | 0.00826 | FES |
| T Helper Cell Differentiation | 0 | 0.0137 | HLA-B |
| CD28 Signaling in T Helper Cells | 0 | 0.0167 | HLA-B,LAT |
| IL-15 Signaling | 0 | 0.0141 | KRAS |
| Reelin Signaling in Neurons | 0 | 0.0155 | ARHGEF10,MAPT |
| Melatonin Signaling | 0 | 0.0139 | PDIA3 |
| Neuropathic Pain Signaling In Dorsal Horn N | 0 | 0.0099 | PDIA3 |
| Factors Promoting Cardiogenesis in Vertebra | 0 | 0.0108 | FZD7 |

| | | | |
|---|---|---|---|
| CNTF Signaling | 0 | 0.0175 | KRAS |
| Renin-Angiotensin Signaling | 0 | 0.00847 | KRAS |
| Corticotropin Releasing Hormone Signaling | 0 | 0.0069 | CRHR1 |
| Mitotic Roles of Polo-Like Kinase | 0 | 0.0152 | CAPN1 |
| HGF Signaling | 0 | 0.00901 | KRAS |
| FLT3 Signaling in Hematopoietic Progenitor | 0 | 0.0125 | KRAS |
| GNRH Signaling | 0 | 0.00578 | KRAS |
| Cholecystokinin/Gastrin-mediated Signaling | 0 | 0.0168 | KRAS,RHOT2 |
| Human Embryonic Stem Cell Pluripotency | 0 | 0.00741 | FZD7 |
| Aldosterone Signaling in Epithelial Cells | 0 | 0.0127 | KRAS,PDIA3 |
| Role of NANOG in Mammalian Embryonic St | 0 | 0.0168 | FZD7,KRAS |
| Type I Diabetes Mellitus Signaling | 0 | 0.018 | CASP9,HLA-B |
| Basal Cell Carcinoma Signaling | 0 | 0.0139 | FZD7 |
| Allograft Rejection Signaling | 0 | 0.0116 | HLA-B |
| Glioma Signaling | 0 | 0.00909 | KRAS |
| Bladder Cancer Signaling | 0 | 0.0103 | KRAS |
| Type II Diabetes Mellitus Signaling | 0 | 0.0141 | SLC27A2,SMPD2 |
| Chronic Myeloid Leukemia Signaling | 0 | 0.00971 | KRAS |
| Gα12/13 Signaling | 0 | 0.0154 | CDH9,KRAS |
| ERK5 Signaling | 0 | 0.0139 | KRAS |
| Cdc42 Signaling | 0 | 0.00599 | HLA-B |
| PAK Signaling | 0 | 0.0103 | KRAS |
| Rac Signaling | 0 | 0.00893 | KRAS |
| Role of Osteoblasts, Osteoclasts and Chonc | 0 | 0.0136 | CASP9,CTSK,FZD7 |
| Ovarian Cancer Signaling | 0 | 0.0144 | FZD7,KRAS |
| Atherosclerosis Signaling | 0 | 0.00794 | PLA2R1 |
| Role of Macrophages, Fibroblasts and Endo | 0 | 0.016 | CEBPZ,FZD7,KRAS,PDIA3,PRSS3 |
| Regulation of eIF4 and p70S6K Signaling | 0 | 0.0127 | KRAS,RPS17 |
| Role of Wnt/GSK-3β Signaling in the Pathog | 0 | 0.0128 | FZD7 |
| Role of PI3K/AKT Signaling in the Pathoger | 0 | 0.0156 | CASP9 |
| MSP-RON Signaling Pathway | 0 | 0.0172 | ACTG1 |
| OX40 Signaling Pathway | 0 | 0.0111 | HLA-B |
| Cell Cycle Control of Chromosomal Replicati | 0 | 0.0179 | ORC5 |
| Role of Tissue Factor in Cancer | 0 | 0.00855 | KRAS |
| NGF Signaling | 0 | 0.0175 | KRAS,SMPD2 |
| Telomerase Signaling | 0 | 0.00935 | KRAS |
| eNOS Signaling | 0 | 0.00629 | CASP9 |
| Netrin Signaling | 0 | 0.0154 | ABLIM3 |
| Nicotine Degradation III | 0 | 0.0179 | CYP51A1 |
| Nicotine Degradation II | 0 | 0.0154 | CYP51A1 |
| Agranulocyte Adhesion and Diapedesis | 0 | 0.0104 | ACTG1,EZR |
| Granulocyte Adhesion and Diapedesis | 0 | 0.00559 | EZR |
| Regulation of the Epithelial-Mesenchymal Tr | 0 | 0.0156 | FZD7,KRAS,SNAI1 |
| Sperm Motility | 0 | 0.0179 | FES,PDE4B,PDIA3,PLA2R1 |
| STAT3 Pathway | 0 | 0.00741 | KRAS |
| Oxidative Phosphorylation | 0 | 0.00917 | NDUFS1 |
| Cell Cycle: G1/S Checkpoint Regulation | 0 | 0.0149 | GNL3 |
| ERK/MAPK Signaling | 0 | 0.0104 | KRAS,PPP1R14A |
| SAPK/JNK Signaling | 0 | 0.0098 | KRAS |
| PI3K/AKT Signaling | 0 | 0.00575 | KRAS |
| PTEN Signaling | 0 | 0.0159 | CASP9,KRAS |
| Nitric Oxide Signaling in the Cardiovascular S | 0 | 0.0101 | ATP2A1 |
| Protein Ubiquitination Pathway | 0 | 0.011 | HLA-B,TAP2,USP13 |
| IL-2 Signaling | 0 | 0.0164 | KRAS |
| JAK/Stat Signaling | 0 | 0.0125 | KRAS |
| GABA Receptor Signaling | 0 | 0.0105 | GABRB2 |
| B Cell Receptor Signaling | 0 | 0.00541 | KRAS |
| Wnt/β-catenin Signaling | 0 | 0.0116 | EP300,FZD7 |
| Chemokine Signaling | 0 | 0.0125 | KRAS |
| Neurotrophin/TRK Signaling | 0 | 0.0132 | KRAS |
| Dopamine Receptor Signaling | 0 | 0.013 | PPP1R14A |
| p38 MAPK Signaling | 0 | 0.00847 | MAPT |
| Glutamate Receptor Signaling | 0 | 0.0175 | HOMER3 |
| NF-κB Signaling | 0 | 0.0112 | EP300,KRAS |
| IL-6 Signaling | 0 | 0.008 | KRAS |
| PDGF Signaling | 0 | 0.0116 | KRAS |
| BMP signaling pathway | 0 | 0.0118 | KRAS |
| GPCR-Mediated Integration of Enteroendoc | 0 | 0.0137 | PDIA3 |
| GPCR-Mediated Nutrient Sensing in Enteroe | 0 | 0.0179 | GNG4,PDIA3 |
| Phagosome Formation | 0 | 0.016 | PDIA3,RHOT2 |
| Macropinocytosis Signaling | 0 | 0.0132 | KRAS |
| Cancer Drug Resistance By Drug Efflux | 0 | 0.0172 | KRAS |
| Th1 and Th2 Activation Pathway | 0 | 0.0117 | HLA-B,SPI1 |
| Th1 Pathway | 0 | 0.00826 | HLA-B |
| Th2 Pathway | 0 | 0.0147 | HLA-B,SPI1 |
| GP6 Signaling Pathway | 0 | 0.0168 | LAMA2,LAT |
| Opioid Signaling Pathway | 0 | 0.0162 | EP300,GRK6,KRAS,MAPK6 |
| Iron homeostasis signaling pathway | 0 | 0.0146 | ARNT,SLC25A37 |
| SPINK1 Pancreatic Cancer Pathway | 0 | 0.0167 | PRSS3 |
| NER Pathway | 0 | 0.00971 | EP300 |
| Endocannabinoid Neuronal Synapse Pathw | 0 | 0.0156 | MAPK6,PDIA3 |

| | | | |
|---|---|---|---|
| Apelin Endothelial Signaling Pathway | 0 | 0.0174 | ARNT,KRAS |
| Cardiac Hypertrophy Signaling (Enhanced) | 0 | 0.0164 | ATP2A1,EP300,FZD7,KRAS,PDE4B,PDIA3,TNFSF13,TULP2 |
| T Cell Exhaustion Signaling Pathway | 0 | 0.0114 | HLA-B,KRAS |
| Systemic Lupus Erythematosus In B Cell Sig | 0 | 0.00727 | KRAS,TNFSF13 |
| White Adipose Tissue Browning Pathway | 0 | 0.0155 | PPARGC1A,RXRB |
| Hepatic Fibrosis Signaling Pathway | 0 | 0.0163 | FZD7,KRAS,LEPR,RHOT2,SNAI1,SOD2 |
| Calcium Signaling | 0 | 0.0146 | ATP2A1,EP300,TNNT2 |
| GM-CSF Signaling | 0 | 0.0143 | KRAS |
