## Supplementary table 3 for "Multi-omics integration analysis identifies novel genes for alcoholism with potential link to neurodegenerative diseases"

**Table 3 a: SMR analysis results with summary statistics of AUD GWAS**

| ProbeChr | Probe_bp | Gene | Adult brain (SMR P value) | | Fetal brain (SMR P value) | |
|---|---|---|---|---|---|---|
| | | | eQTL | mQTL | eQTL | mQTL |
| 1 | 27696114 | *PHACTR4* | 1.01E-03 | x | x | 8.00E-04 |
| 1 | 42824626 | *CDC20* | 1.70E-02 | x | x | 1.56E-03 |
| 1 | 46090169 | *CCDC17* | x | x | x | 8.16E-04 |
| 3 | 48395716 | *GPX1* | 4.93E-05 | x | x | x |
| | 48459884 | *AMT* | 2.07E-04 | x | 4.39E-04 | 4.47E-01 |
| 6 | 31324491 | *HLA-B* | x | 4.67E-02 | x | 1.12E-03 |
| 6 | 100329187 | *ASCC3* | 5.55E-01 | x | x | 9.92E-04 |
| 7 | 157334298 | *PTPRN2* | 5.00E-02 | 1.33E-03 | x | 1.39E-03 |
| 8 | 9859991 | *XKR6* | 8.62E-01 | 6.81E-01 | x | 8.06E-04 |
| 8 | 56358589 | *PENK* | 4.00E-01 | 4.16E-01 | x | 1.75E-03 |
| 9 | 99975832 | *TBC1D2* | 2.22E-01 | x | 1.02E-03 | x |
| 11 | 46399942 | *SPI1* | x | x | x | 1.91E-04 |
| | 46664086 | *MTCH2* | 1.89E-05 | x | x | x |
| | 46843734 | *NUP160* | 3.88E-04 | x | x | x |
| 13 | 112548692 | *MCF2L* | 6.22E-02 | 3.71E-03 | x | 1.13E-03 |
| 16 | 22765948 | *CHP2* | 1.10E-01 | 1.01E-03 | x | 1.79E-03 |
| 17 | 27256874 | *EFCAB5* | 9.57E-04 | x | x | 1.22E-03 |
| 17 | 43361331 | *MAP3K14* | x | x | x | 2.99E-05 |
| 19 | 12617038 | *CACNA1A* | x | x | x | 1.45E-03 |
| 19 | 17586743 | *ELL* | 3.03E-03 | x | x | 3.36E-04 |
| 19 | 56988872 | *ZNF772* | 8.63E-01 | x | x | 1.45E-03 |

**Table 3 b: SMR analysis results with summary statistics of DPW GWAS**

| ProbeChr | Probe_bp | Gene | Adult brain (SMR P values) | | Fetal brain (SMR P values) | |
|---|---|---|---|---|---|---|
| | | | eQTL | mQTL | eQTL | mQTL |
| 1 | 14850790 | *CASP9* | 1.03E-03 | 4.60E-04 | NA | NA |
| | 25644448 | *CD52* | 6.28E-03 | NA | NA | 7.37E-03 |
| | 15678949 | *FBXO42* | 7.78E-03 | NA | NA | NA |
| | 25496362 | *ZNF593* | 8.88E-03 | 2.32E-02 | NA | NA |
| | 25503894 | *CNKSR1* | 9.05E-03 | NA | 1.53E-02 | 3.48E-02 |
| | 25633111 | *UBXN11* | 4.30E-03 | 1.55E-01 | NA | 3.96E-03 |
| 1 | 30661000 | *NKAIN1* | 3.76E-03 | NA | NA | NA |
| 1 | 46799469 | *CMPK1* | 6.60E-05 | 1.14E-04 | NA | NA |
| | 46090169 | *CCDC17* | NA | NA | NA | 9.93E-04 |
| | 46045924 | *NASP* | NA | 1.32E-03 | NA | NA |
| 1 | 64886248 | *LEPR* | 7.46E-03 | 5.57E-01 | NA | NA |
| 1 | 66257822 | *PDE4B* | NA | NA | NA | 1.06E-02 |
| 1 | 70513281 | *PTGER3* | 7.88E-03 | 6.26E-05 | NA | NA |
| | 71566613 | *NEGR1* | 2.81E-04 | NA | NA | NA |
| 1 | 92415048 | *BRDT* | NA | 9.63E-04 | NA | NA |
| | 91495539 | *EPHX4* | 4.15E-03 | NA | NA | NA |
| 1 | 155698234 | *RRNAD1* | NA | 3.07E-05 | NA | NA |
| | 156722410 | *HDGF* | NA | 3.78E-05 | NA | NA |
| | 154023762 | *ADAM15* | 2.29E-03 | NA | NA | NA |
| | 149738268 | *CTSS* | 2.42E-03 | NA | NA | NA |
| | 150897856 | *SETDB1* | NA | NA | NA | 2.47E-03 |
| | 149941588 | *CERS2* | 2.67E-03 | NA | NA | NA |
| | 149693352 | *HORMAD1* | 3.19E-03 | NA | NA | NA |
| | 154036224 | *EFNA3* | 4.72E-03 | NA | NA | NA |
| | 149849055 | *ARNT* | 1.04E-02 | NA | NA | 6.25E-03 |
| | 149780799 | *CTSK* | 9.41E-03 | NA | NA | NA |
| | 151040405 | *MLLT11* | NA | 1.09E-02 | NA | 1.07E-02 |
| 1 | 204819245 | *PM20D1* | 3.32E-05 | NA | NA | 2.69E-04 |
| | 200342382 | *TNNT2* | 5.79E-03 | NA | NA | NA |
| | 204290883 | *NUAK2* | 1.04E-02 | NA | NA | 7.56E-03 |
| | 204904950 | *SLC26A9* | 1.58E-03 | 9.76E-01 | NA | NA |
| 1 | 219863187 | *C1orf115* | 1.43E-03 | NA | NA | NA |
| | 225054448 | *TMEM63A* | 1.52E-03 | NA | NA | NA |
| | 234667722 | *B3GALNT2* | 4.98E-03 | NA | NA | 1.78E-03 |
| | 230376913 | *C1orf131* | 3.91E-03 | NA | NA | NA |
| | 234758779 | *GNG4* | 4.30E-03 | NA | NA | NA |
| | 219921611 | *2-Mar* | 4.34E-03 | NA | NA | NA |
| | 230762561 | *DISC1* | 6.94E-03 | NA | NA | NA |
| | 224997797 | *EPHX1* | 3.34E-01 | NA | NA | 1.20E-05 |
| 2 | 1 | *TMEM18* | 2.23E-01 | 4.79E-03 | NA | 2.30E-01 |
| 2 | 63277830 | *OTX1* | NA | NA | NA | 1.92E-04 |
| | 36458856 | *CEBPZ* | 9.77E-04 | NA | NA | NA |
| | 60293068 | *KIAA1841* | 2.13E-03 | NA | NA | NA |
| | 25915619 | *KCNK3* | 2.78E-03 | NA | NA | NA |
| | 24295212 | *SF3B6* | NA | 3.66E-03 | NA | NA |
| | 24149416 | *ATAD2B* | NA | NA | NA | 3.86E-03 |
| | 54237585 | *RTN4* | 6.37E-03 | NA | NA | NA |
| | 23346347 | *PFN4* | 6.69E-03 | NA | NA | NA |
| | 46572297 | *EPCAM* | 6.79E-03 | NA | 1.16E-02 | NA |

| | | | | | | |
|---|---|---|---|---|---|---|
| | 47757064 | *TON1-GTF2A1* | 1.99E-02 | 2.45E-03 | 1.52E-02 | 3.46E-03 |
| | 61370764 | *C2orf74* | NA | 1.02E-01 | 2.20E-03 | NA |
| | 29454397 | *LBH* | 1.31E-03 | 1.65E-01 | NA | NA |
| | 44312696 | *PPM1B* | NA | 5.25E-05 | NA | 2.04E-01 |
| | 31502979 | *YIPF4* | 9.42E-03 | NA | NA | 3.53E-01 |
| | 15804344 | *FAM49A* | 1.66E-04 | 3.85E-01 | NA | NA |
| | 23921236 | *KLHL29* | NA | 2.36E-03 | NA | 4.29E-01 |
| 2 | 72612886 | *ALMS1* | 4.61E-05 | NA | NA | NA |
| | 99952887 | *TXNDC9* | NA | 2.00E-04 | NA | NA |
| | 99106435 | *REV1* | NA | NA | NA | 6.03E-04 |
| | 98771418 | *LIPT1* | 1.15E-03 | NA | NA | 1.01E-03 |
| | 98561179 | *TMEM131* | NA | NA | NA | 1.34E-03 |
| | 97280570 | *ACTR1B* | 2.71E-03 | NA | NA | 1.67E-03 |
| | 98735242 | *TSGA10* | 1.47E-03 | 4.28E-03 | 2.63E-03 | 4.91E-04 |
| | 98771461 | *MRPL30* | 2.51E-03 | NA | NA | NA |
| | 98552722 | *KIAA1211L* | 3.72E-03 | NA | NA | NA |
| | 112342018 | *CHCHD5* | 4.25E-03 | NA | NA | NA |
| | 87469835 | *THNSL2* | 4.54E-03 | NA | 8.59E-03 | NA |
| | 106503102 | *ST6GAL2* | 1.07E-02 | 5.83E-03 | NA | NA |
| | 84811531 | *VAMP5* | NA | NA | NA | 1.10E-02 |
| | 112191107 | *RGPD8* | 2.88E-03 | 6.25E-03 | NA | 9.86E-01 |
| 2 | 152032506 | *STAM2* | 5.66E-04 | 4.67E-04 | NA | NA |
| | 160758480 | *LY75-CD302* | NA | 1.86E-03 | NA | NA |
| | 160760732 | *LY75* | NA | NA | NA | 2.18E-03 |
| | 160917029 | *PLA2R1* | NA | 3.69E-03 | NA | NA |
| | 169551001 | *KLHL23* | 6.30E-03 | NA | NA | NA |
| | 163592517 | *FIGN* | 4.48E-01 | 1.16E-03 | NA | NA |
| 2 | 178160898 | *NFE2L2* | NA | 2.72E-05 | NA | 1.73E-01 |
| 2 | 210341429 | *LANCL1* | 7.02E-04 | 1.25E-03 | NA | 8.40E-04 |
| | 202900408 | *FZD7* | NA | 4.68E-03 | NA | NA |
| | 206023918 | *NDUFS1* | 9.52E-02 | 5.17E-03 | NA | NA |
| | 205547224 | *NRP2* | NA | 6.17E-01 | NA | 9.67E-03 |
| | 210342433 | *CPS1* | 6.67E-04 | 9.02E-01 | NA | NA |
| 2 | 224428016 | *CUL3* | 1.95E-03 | 1.02E-01 | NA | 4.73E-03 |
| 2 | 237536219 | *LRRFIP1* | 7.42E-03 | 4.50E-01 | NA | NA |
| 2 | 238707032 | *RBM44* | NA | NA | 1.30E-03 | NA |
| | 242930205 | *RTP5* | NA | 2.64E-03 | NA | NA |
| | 241082488 | *PASK* | 7.39E-01 | 1.22E-03 | NA | 6.38E-01 |
| 3 | 18735633 | *SATB1* | NA | 4.50E-03 | NA | NA |
| 3 | 37537763 | *EXOG* | 6.34E-04 | 1.89E-03 | NA | NA |
| 3 | 48395716 | *GPX1* | 3.61E-03 | NA | NA | NA |
| | 48459884 | *AMT* | 3.51E-03 | NA | 4.96E-03 | 1.18E-02 |
| 3 | 51719936 | *GNL3* | 6.79E-03 | NA | NA | 5.19E-02 |
| 3 | 131378688 | *ACAD11* | 5.59E-03 | 2.27E-01 | NA | NA |
| 3 | 142666759 | *PAQR9* | NA | 1.55E-03 | NA | NA |
| 3 | 161249750 | *OTOL1* | NA | 1.40E-03 | NA | NA |
| | 182883005 | *LAMP3* | NA | 2.85E-03 | NA | NA |
| | 182732192 | *ABCC5* | 5.47E-03 | 8.91E-03 | NA | 3.57E-03 |
| | 170076028 | *SKIL* | NA | NA | NA | 6.89E-03 |
| | 178370543 | *USP13* | 8.15E-03 | NA | NA | NA |
| 3 | 182967438 | *ECE2* | 1.27E-05 | NA | NA | NA |
| 3 | 194538728 | *MUC4* | 9.34E-01 | 5.62E-02 | NA | 1.11E-02 |

| | | | | | | |
|---|---|---|---|---|---|---|
| 4 | 1 | *ZNF721* | 2.79E-03 | NA | NA | NA |
| 4 | 817492 | *CPLX1* | NA | 9.73E-01 | NA | 5.17E-03 |
| 4 | 2465033 | *DOK7* | 2.38E-01 | 1.41E-03 | NA | 2.27E-01 |
| 4 | 22891658 | *PPARGC1A* | NA | 4.78E-03 | NA | NA |
| 4 | 38460508 | *RPL9* | 5.43E-05 | NA | 2.95E-04 | NA |
| | 38460620 | *LIAS* | 2.66E-03 | 8.95E-03 | NA | 5.36E-03 |
| | 40992489 | *SLC30A9* | 7.62E-03 | NA | NA | NA |
| 4 | 56500362 | *NMU* | NA | 6.19E-03 | NA | NA |
| 4 | 56844989 | *NOA1* | 6.57E-04 | NA | NA | NA |
| 4 | 76328458 | *CCDC158* | 5.29E-03 | NA | NA | NA |
| 4 | 88744352 | *FAM13A* | 7.85E-04 | 3.84E-03 | NA | NA |
| 4 | 101341118 | *BANK1* | NA | 4.57E-03 | NA | NA |
| | 103119565 | *CENPE* | 5.10E-03 | NA | NA | NA |
| | 102790135 | *CISD2* | 5.15E-03 | NA | NA | NA |
| | 107910870 | *HADH* | 5.72E-03 | NA | NA | NA |
| | 99009902 | *ADH5* | 4.10E-03 | NA | NA | 1.62E-02 |
| | 100439158 | *EMCN* | 1.66E-01 | 5.39E-03 | NA | NA |
| 4 | 112437328 | *NEUROG2* | 4.81E-03 | NA | NA | NA |
| 4 | 173292094 | *SAP30* | 7.28E-03 | NA | NA | NA |
| | 173320687 | *SCRG1* | 1.08E-03 | NA | NA | NA |
| 5 | 26038693 | *CDH9* | 1.82E-01 | 8.17E-04 | NA | NA |
| 5 | 49708521 | *EMB* | NA | 1.05E-02 | NA | 1.11E-02 |
| 5 | 56878867 | *RAB3C* | 5.89E-03 | 8.59E-01 | NA | NA |
| 5 | 77365540 | *BHMT2* | 5.62E-03 | NA | NA | NA |
| 5 | 79256491 | *RASGRF2* | 2.82E-02 | 2.97E-02 | NA | 3.62E-04 |
| 5 | 111849380 | *YTHDC2* | 7.01E-03 | NA | NA | NA |
| 5 | 132209139 | *LEAP2* | NA | 7.75E-05 | NA | 5.15E-05 |
| | 145583025 | *RBM27* | NA | 1.27E-03 | NA | NA |
| | 157690089 | *UBLCP1* | 1.45E-03 | NA | NA | NA |
| | 129843850 | *RAPGEF6* | 3.96E-03 | NA | NA | NA |
| | 150151471 | *G3BP1* | 5.37E-03 | NA | NA | 4.33E-03 |
| | 148569520 | *SLC6A7* | 5.07E-03 | NA | NA | NA |
| | 139514800 | *PCDHB5* | 5.71E-03 | NA | NA | NA |
| | 139552243 | *PCDHB7* | 8.77E-02 | 4.74E-03 | NA | NA |
| | 139571942 | *PCDHB10* | 1.21E-01 | 5.92E-03 | NA | NA |
| | 131948255 | *FSTL4* | 4.18E-01 | 2.53E-03 | NA | NA |
| | 147521046 | *ABLIM3* | 4.16E-03 | NA | NA | 8.06E-01 |
| 5 | 160975347 | *GABRB2* | NA | NA | NA | 1.03E-02 |
| 5 | 175853679 | *GRK6* | NA | NA | NA | 1.15E-04 |
| | 174487692 | *FAM153B* | 4.84E-03 | NA | NA | NA |
| | 179688119 | *TRIM52* | 2.95E-02 | 5.28E-03 | NA | NA |
| 6 | 15129356 | *MYLIP* | 8.00E-03 | NA | NA | NA |
| 6 | 32168391 | *RXRB* | 4.37E-04 | NA | NA | NA |
| | 33396215 | *SYNGAP1* | NA | 4.09E-03 | NA | NA |
| | 31324491 | *HLA-B* | NA | 1.27E-03 | NA | 7.23E-03 |
| | 32030123 | *TNXB* | NA | 4.44E-03 | NA | NA |
| | 27891765 | *TRIM27* | NA | 5.34E-03 | NA | 1.00E-02 |
| | 31806504 | *TAP2* | 8.27E-03 | 2.01E-02 | NA | 4.59E-02 |
| | 33725183 | *SNRPC* | 8.62E-02 | 4.15E-03 | NA | NA |
| 6 | 41928496 | *GNMT* | 4.28E-03 | 4.04E-03 | NA | 3.64E-03 |
| 6 | 83140764 | *ME1* | 4.23E-02 | 2.92E-03 | NA | NA |
| | 83222194 | *PRSS35* | 5.83E-03 | NA | NA | NA |

| | | | | | | |
|---|---|---|---|---|---|---|
| 6 | 104307794 | *HACE1* | 1.82E-04 | NA | NA | NA |
| | 108761966 | *SMPD2* | 4.29E-03 | NA | NA | NA |
| 6 | 128204342 | *LAMA2* | 8.62E-01 | 3.33E-03 | NA | NA |
| 6 | 149067658 | *NUP43* | 5.69E-03 | NA | NA | NA |
| | 148887430 | *GINM1* | 5.72E-03 | NA | NA | NA |
| | 149023490 | *LATS1* | 6.98E-03 | NA | NA | NA |
| | 159114260 | *SOD2* | 5.95E-03 | 1.18E-01 | NA | NA |
| | 158239259 | *EZR* | 1.75E-01 | NA | NA | 6.98E-03 |
| 6 | 169151718 | *ERMARD* | 3.22E-01 | 1.89E-03 | NA | NA |
| 7 | 26702614 | *HIBADH* | 9.23E-04 | 2.21E-01 | NA | 2.11E-01 |
| | 33697851 | *NPSR1* | 5.64E-04 | NA | NA | NA |
| 7 | 55131695 | *SUMF2* | 2.95E-03 | NA | NA | 4.39E-03 |
| 7 | 75091018 | *DTX2* | 1.70E-01 | 5.14E-03 | NA | NA |
| | 75139745 | *UPK3B* | 3.24E-03 | 2.95E-03 | NA | 2.92E-03 |
| 7 | 99026440 | *MEPCE* | 1.08E-03 | 1.05E-03 | NA | 1.11E-03 |
| | 99934006 | *SPDYE3* | NA | 1.22E-03 | NA | NA |
| | 99076855 | *TSC22D4* | NA | NA | NA | 1.55E-03 |
| | 99017500 | *ZCWPW1* | 1.56E-03 | NA | NA | NA |
| | 98933737 | *PILRB* | 1.07E-03 | NA | 2.18E-03 | NA |
| | 98971068 | *PILRA* | 9.78E-04 | NA | 2.56E-03 | NA |
| | 102848405 | *ORC5* | 1.87E-03 | NA | NA | NA |
| | 99061894 | *C7orf61* | 2.04E-03 | NA | NA | NA |
| | 99081550 | *NYAP1* | 3.20E-03 | NA | NA | NA |
| | 104580787 | *LHFPL3* | NA | 5.60E-03 | NA | NA |
| | 98775186 | *STAG3* | 1.04E-01 | 4.39E-03 | NA | 3.73E-02 |
| | 90763844 | *CYP51A1* | 1.15E-04 | 7.56E-01 | NA | NA |
| 7 | 126225661 | *GCC1* | 7.86E-04 | NA | NA | NA |
| | 127379346 | *CALU* | 3.78E-03 | 9.31E-03 | NA | NA |
| | 140178322 | *MKRN1* | NA | 1.08E-02 | NA | 1.07E-02 |
| | 139714229 | *MRPS33* | 7.23E-03 | 5.79E-01 | NA | NA |
| 7 | 157334298 | *PTPRN2* | 6.42E-01 | 6.97E-03 | NA | 2.76E-03 |
| 8 | 772142 | *ARHGEF10* | 7.22E-01 | 1.30E-02 | NA | 1.26E-03 |
| 8 | 37034192 | *LSM1* | 1.38E-05 | 1.35E-05 | NA | NA |
| | 37088861 | *DDHD2* | 9.76E-05 | NA | NA | NA |
| | 26630151 | *CCDC25* | 1.05E-04 | NA | NA | NA |
| | 41396298 | *SMIM19* | 8.84E-04 | NA | NA | NA |
| | 27630920 | *ESCO2* | NA | NA | NA | 2.14E-03 |
| | 37218068 | *ZNF703* | NA | 5.69E-03 | NA | NA |
| | 16743288 | *FGL1* | 5.77E-03 | NA | NA | NA |
| | 41358978 | *SLC20A2* | 6.51E-03 | NA | NA | NA |
| | 22386318 | *SLC25A37* | 1.48E-02 | 1.19E-03 | NA | NA |
| 8 | 56906403 | *IMPAD1* | NA | 4.93E-03 | NA | NA |
| 8 | 108094184 | *RSPO2* | 2.30E-04 | 8.89E-01 | NA | NA |
| | 115632297 | *TRPS1* | 1.03E-03 | 4.95E-01 | NA | NA |
| 8 | 141138711 | *DENND3* | 8.49E-03 | 7.63E-02 | NA | NA |
| | 143623623 | *ZC3H3* | 2.73E-03 | 1.75E-01 | NA | NA |
| 9 | 32750515 | *PRSS3* | 4.26E-03 | 6.84E-03 | NA | 9.27E-01 |
| 9 | 90926113 | *CKS2* | 7.04E-03 | NA | NA | NA |
| | 92506456 | ***GADD45G*** | NA | 4.79E-03 | NA | NA |
| | 102114853 | *TEX10* | 9.21E-04 | NA | NA | NA |
| | 127244764 | *NR5A1* | NA | NA | NA | 6.94E-06 |
| | 129213652 | *RPL12* | 1.65E-05 | NA | NA | NA |

| | | | | | | |
|---|---|---|---|---|---|---|
| 9 | 129186661 | *ZNF79* | NA | 1.04E-04 | NA | NA |
| | 128884913 | *ANGPTL2* | 1.05E-03 | NA | NA | NA |
| | 124675609 | *ZBTB6* | 3.43E-03 | NA | NA | NA |
| | 83069149 | *NRG3* | NA | 3.90E-03 | NA | NA |
| | 74757872 | *VCL* | 4.29E-03 | NA | NA | NA |
| | 124703288 | *RABGAP1* | 4.86E-03 | NA | NA | NA |
| | 129159421 | *SLC2A8* | 2.59E-04 | 5.79E-02 | NA | NA |
| | 85088342 | *CCSER2* | 2.54E-01 | 3.46E-03 | NA | NA |
| | 69587298 | *STOX1* | 2.60E-04 | 6.56E-01 | 1.58E-01 | NA |
| 10 | 104036920 | *INA* | 4.49E-03 | NA | NA | NA |
| 10 | 123913793 | *BUB3* | NA | NA | NA | 1.11E-02 |
| 11 | 7960899 | *OR10A3* | NA | 1.34E-03 | NA | NA |
| | 8685624 | *SWAP70* | 6.19E-03 | NA | NA | 6.78E-01 |
| | 17034500 | *SERGEF* | 9.44E-01 | NA | NA | 6.31E-03 |
| 11 | 46399942 | *SPI1* | NA | NA | NA | 1.11E-06 |
| | 46843734 | *NUP160* | 1.43E-06 | NA | NA | NA |
| | 46616211 | *C1QTNF4* | 5.26E-06 | NA | NA | NA |
| | 45722368 | *ZNF408* | 3.37E-03 | NA | NA | NA |
| | 46260618 | *CREB3L1* | NA | 3.64E-03 | NA | NA |
| | 33185376 | *CSTF3* | NA | 7.94E-02 | NA | 4.96E-06 |
| | 42702259 | *HSD17B12* | 5.55E-01 | 8.90E-04 | NA | NA |
| | 33937677 | *PDHX* | 9.77E-01 | 4.83E-03 | NA | 6.61E-04 |
| | 42902389 | *ALKBH3* | 2.46E-03 | 6.91E-01 | NA | NA |
| | 27494710 | *LGR4* | NA | 9.25E-01 | NA | 5.02E-05 |
| 11 | 62439084 | *ATL3* | 1.41E-03 | NA | NA | NA |
| | 60572152 | *FADS1* | 1.71E-03 | NA | NA | NA |
| | 67228186 | *PPP6R3* | 2.05E-03 | NA | NA | NA |
| | 60560452 | *FADS2* | NA | NA | NA | 2.94E-03 |
| | 60560082 | *TMEM258* | 6.10E-03 | NA | NA | NA |
| | 65313706 | *ZDHHC24* | 6.94E-03 | NA | NA | NA |
| | 65360292 | *CCS* | 8.56E-03 | NA | NA | NA |
| | 63948665 | *CAPN1* | NA | NA | NA | 9.79E-03 |
| | 61378479 | *EML3* | NA | NA | NA | 1.06E-02 |
| | 71492887 | *STARD10* | 4.20E-03 | NA | NA | 3.51E-02 |
| 11 | 121526383 | *UBASH3B* | NA | 4.99E-03 | NA | NA |
| 12 | 2186521 | *TSPAN9* | 8.22E-04 | 7.38E-01 | NA | 1.41E-01 |
| | 5661066 | *IFFO1* | 6.96E-03 | NA | NA | NA |
| | 5862082 | *MLF2* | 8.81E-03 | NA | NA | 1.57E-03 |
| 12 | 24403737 | *KRAS* | 5.48E-03 | 7.27E-01 | NA | NA |
| | 32592581 | *SYT10* | 5.92E-04 | NA | NA | NA |
| 12 | 50785101 | *SLC4A8* | 3.95E-05 | NA | NA | NA |
| | 57026430 | *B4GALNT1* | 1.50E-03 | NA | NA | NA |
| | 47919415 | *OR8S1* | 3.49E-03 | NA | NA | NA |
| | 56328189 | *SDR9C7* | 6.73E-03 | NA | NA | NA |
| | 53582391 | *SMUG1* | 7.05E-03 | NA | NA | NA |
| 12 | 80470990 | *ACSS3* | 3.49E-04 | NA | NA | NA |
| | 90572329 | *DCN* | 4.31E-03 | NA | NA | NA |
| | 109486327 | *C12orf76* | 7.36E-03 | 1.35E-02 | NA | NA |
| | 104380088 | *C12orf45* | 2.73E-01 | NA | NA | 7.99E-03 |
| 12 | 122459127 | *OGFOD2* | 5.81E-03 | NA | NA | NA |
| | 132195366 | *P2RX2* | 3.03E-02 | NA | NA | 1.09E-02 |
| | 132287405 | *PGAM5* | 3.73E-03 | 3.39E-02 | NA | NA |

| | | | | | | |
|---|---|---|---|---|---|---|
| | 133049993 | *FBRSL1* | NA | 1.37E-03 | NA | 4.12E-02 |
| 13 | 50483814 | *RNASEH2B* | 1.20E-04 | NA | NA | NA |
| | 40768702 | *KBTBD7* | 1.93E-03 | NA | NA | NA |
| | 50417832 | *DLEU7* | 5.36E-03 | NA | NA | NA |
| | 41846289 | *AKAP11* | 9.17E-03 | NA | NA | NA |
| | 48925263 | *CAB39L* | 1.15E-04 | 6.27E-01 | NA | NA |
| 13 | 100150906 | *TM9SF2* | NA | 1.32E-03 | NA | NA |
| | 106866120 | *EFNB2* | NA | 2.59E-03 | NA | NA |
| | 96873688 | *MBNL2* | NA | NA | NA | 5.17E-03 |
| | 107519083 | *FAM155A* | 3.10E-01 | 1.97E-03 | NA | NA |
| 14 | 47121033 | *RPL10L* | NA | 3.55E-03 | NA | NA |
| 14 | 72704205 | *PAPLN* | NA | 5.40E-03 | NA | NA |
| 15 | 51030318 | *LYSMD2* | 1.44E-04 | NA | NA | 1.95E-04 |
| | 52264396 | *LEO1* | NA | 2.54E-04 | NA | NA |
| | 51311398 | *MAPK6* | 2.63E-04 | NA | NA | NA |
| | 48447962 | *GALK2* | 3.66E-04 | NA | NA | NA |
| | 52405000 | *BCL2L10* | NA | NA | NA | 7.81E-04 |
| | 42662108 | *ZSCAN29* | 9.49E-04 | NA | NA | NA |
| | 43038590 | *PDIA3* | 1.18E-03 | NA | NA | NA |
| | 42559055 | *TGM5* | 1.35E-03 | NA | NA | NA |
| | 40522889 | *EXD1* | 2.00E-03 | NA | NA | NA |
| | 49474393 | *SLC27A2* | 4.22E-03 | NA | NA | NA |
| | 52526871 | *MYO5C* | NA | 7.86E-02 | NA | 6.66E-03 |
| | 42876702 | *PPIP5K1* | 2.63E-03 | 2.34E-01 | NA | NA |
| | 51472084 | *GNB5* | 2.85E-03 | 3.32E-01 | NA | NA |
| | 49406701 | *ATP8B4* | 4.41E-03 | 3.60E-01 | NA | NA |
| | 50633826 | *GLDN* | 8.00E-01 | NA | NA | 1.00E-03 |
| | 50200870 | *AP4E1* | 9.90E-01 | 2.58E-01 | NA | 9.40E-03 |
| 15 | 1 | *JMJD8* | 4.77E-03 | NA | NA | NA |
| | 82240509 | *CPEB1* | 5.02E-03 | NA | NA | NA |
| | 89358094 | *ANPEP* | NA | 5.91E-03 | NA | 6.41E-03 |
| | 1 | *RHOT2* | 7.88E-03 | NA | NA | NA |
| | 81824867 | *RPS17* | 9.09E-03 | NA | NA | NA |
| | 876968 | *FAHD1* | 9.19E-03 | NA | 1.42E-02 | 6.80E-03 |
| | 74165706 | *SCAMP2* | 2.34E-02 | NA | NA | 3.41E-04 |
| | 79189391 | *MTHFS* | 2.82E-02 | NA | NA | 9.61E-03 |
| | 90427642 | *FES* | 9.66E-03 | NA | NA | 8.25E-02 |
| | 82735903 | *BTBD1* | 2.46E-03 | NA | NA | 1.42E-01 |
| | 918186 | *MEIOB* | 7.39E-03 | 3.71E-01 | NA | NA |
| 16 | 30022548 | *DOC2A* | NA | NA | NA | 6.13E-07 |
| | 29102570 | *TBX6* | 1.70E-06 | NA | NA | NA |
| | 28915196 | *ATP2A1* | NA | NA | NA | 2.57E-06 |
| | 27620241 | *SULT1A1* | 2.09E-09 | 6.75E-06 | NA | NA |
| | 29662188 | *PRR14* | NA | 1.62E-04 | NA | NA |
| | 28937557 | *KCTD13* | 1.16E-03 | NA | NA | NA |
| | 28911700 | *ASPHD1* | 2.34E-03 | NA | NA | NA |
| | 27607801 | *SULT1A2* | 2.90E-09 | 1.56E-02 | NA | NA |
| | 27996147 | *LAT* | 1.77E-06 | 9.09E-02 | NA | NA |
| | 24269252 | *ZKSCAN2* | 2.68E-01 | 1.38E-03 | NA | NA |
| | 68221032 | *SNTB2* | 3.79E-05 | NA | NA | NA |
| | 68345259 | *VPS4A* | 4.49E-05 | NA | NA | NA |
| | 70611033 | *TAT* | 7.38E-05 | NA | NA | NA |

| | | | | | | |
|---|---|---|---|---|---|---|
| | 85731409 | *GINS2* | NA | 1.51E-04 | NA | 4.50E-08 |
| | 68760409 | *NQO1* | 1.95E-04 | NA | NA | NA |
| | 68373333 | *NIP7* | 2.40E-04 | NA | NA | NA |
| | 71879545 | *ATXN1L* | NA | NA | NA | 2.77E-04 |
| | 68796209 | *WWP2* | 3.11E-04 | NA | NA | NA |
| | 69207722 | *CLEC18C* | 5.84E-04 | 4.37E-04 | NA | NA |
| | 86970122 | *CA5A* | 6.32E-04 | NA | NA | NA |
| 16 | 84744456 | *C16orf74* | NA | 3.34E-03 | NA | NA |
| | 82932731 | *MLYCD* | 4.97E-03 | NA | NA | NA |
| | 70781299 | *AP1G1* | 5.25E-03 | NA | NA | NA |
| | 49099852 | *HEATR3* | 2.14E-03 | NA | 8.53E-03 | 9.01E-03 |
| | 77246968 | *SYCE1L* | NA | NA | NA | 7.61E-03 |
| | 72929007 | *ZFHX3* | NA | 2.99E-01 | NA | 5.69E-03 |
| | 88574811 | *SPG7* | 1.13E-01 | 4.48E-01 | NA | 7.84E-03 |
| | 74808750 | *FA2H* | NA | 1.96E-03 | NA | 3.80E-01 |
| | 68984810 | *CLEC18A* | 3.04E-04 | NA | 8.31E-01 | NA |
| | 71158752 | *PMFBP1* | 2.63E-03 | 9.05E-01 | NA | NA |
| | 6589389 | *WRAP53* | 5.58E-05 | NA | NA | NA |
| | 6461609 | *TNFSF13* | 5.99E-05 | NA | NA | NA |
| | 16723265 | *SREBF1* | NA | 1.16E-04 | NA | 2.33E-04 |
| | 6482785 | *CD68* | NA | 3.04E-04 | NA | NA |
| | 17218624 | *SMCR8* | 3.94E-04 | NA | NA | NA |
| | 1206998 | *SRR* | 3.61E-04 | 6.13E-05 | 2.80E-03 | NA |
| | 7835329 | *TRAPPC1* | NA | NA | NA | 2.04E-03 |
| 17 | 6832753 | *KCNAB3* | 2.96E-03 | NA | NA | NA |
| | 16942606 | *GID4* | 3.91E-03 | NA | NA | NA |
| | 6531123 | *SAT2* | 4.15E-03 | NA | NA | NA |
| | 15256770 | *CENPV* | 4.83E-03 | NA | NA | NA |
| | 16775378 | *TOM1L2* | 1.91E-04 | NA | NA | 1.75E-02 |
| | 17012020 | *MYO15A* | 1.92E-02 | 4.51E-02 | 2.58E-02 | 1.58E-04 |
| | 6620672 | *DNAH2* | 4.74E-02 | 1.17E-03 | NA | NA |
| | 16991200 | *DRG2* | 5.28E-02 | NA | NA | 5.14E-03 |
| | 17163848 | *MIEF2* | 9.84E-02 | 6.31E-04 | NA | NA |
| | 43370099 | *LRRC37A* | 8.35E-16 | 3.89E-13 | 2.18E-10 | NA |
| | 42971748 | *MAPT* | 4.84E-16 | 1.77E-08 | NA | 1.80E-12 |
| | 42699267 | *CRHR1* | 1.07E-08 | 8.67E-09 | NA | NA |
| | 42568094 | *PLEKHM1* | NA | 3.89E-06 | NA | NA |
| | 56232535 | *SKA2* | 2.47E-04 | NA | NA | NA |
| | 27804380 | *GOSR1* | 1.89E-04 | 8.45E-04 | NA | NA |
| | 39811323 | *TUBG2* | 6.29E-04 | 5.10E-04 | NA | NA |
| | 26945135 | *CORO6* | 1.03E-03 | NA | NA | NA |
| | 39761694 | *TUBG1* | 1.25E-03 | NA | NA | NA |
| | 28624429 | *OMG* | 2.45E-03 | NA | NA | 1.13E-04 |
| | 40713999 | *COASY* | NA | NA | NA | 1.34E-03 |
| | 65031635 | *KPNA2* | 2.90E-03 | 1.04E-04 | NA | NA |
| | 56183829 | *TRIM37* | 1.02E-03 | NA | NA | 2.19E-03 |
| | 27256874 | *EFCAB5* | 2.24E-03 | NA | NA | 1.47E-03 |
| | 27257080 | *SSH2* | NA | 2.34E-03 | NA | NA |
| | 64989640 | *C17orf58* | 2.40E-03 | NA | NA | NA |
| | 46022172 | *SNF8* | 2.62E-03 | NA | NA | NA |
| 17 | 36820440 | *TCAP* | 3.14E-03 | NA | NA | NA |
| | 37608096 | *MED1* | NA | NA | NA | 3.62E-03 |

| | | | | | | |
|---|---|---|---|---|---|---|
| | 55769384 | *TEX14* | 3.79E-03 | NA | NA | NA |
| | 36844310 | *PGAP3* | 4.48E-03 | NA | NA | NA |
| | 64821780 | *BPTF* | 4.59E-03 | NA | NA | NA |
| | 37824526 | *PNMT* | NA | NA | NA | 4.90E-03 |
| | 39834631 | *CNTNAP1* | 5.20E-03 | NA | NA | NA |
| | 36793318 | *STARD3* | NA | NA | NA | 6.83E-03 |
| | 57156282 | *HEATR6* | 7.71E-03 | NA | NA | NA |
| | 26621133 | *NUFIP2* | 7.82E-03 | NA | NA | NA |
| | 45178560 | *CBX1* | 7.82E-03 | NA | NA | NA |
| | 72901494 | *MRPL38* | 8.99E-03 | NA | NA | NA |
| | 39701232 | *HSD17B1* | 1.76E-02 | 1.36E-03 | NA | NA |
| | 77234665 | *RNF213* | 2.46E-03 | 1.72E-01 | 2.17E-02 | NA |
| | 78479806 | *ACTG1* | 4.89E-01 | 2.45E-03 | NA | 1.04E-02 |
| | 36557876 | *FBXL20* | 7.81E-01 | 1.65E-03 | NA | 1.66E-03 |
| | 77518619 | *RPTOR* | 9.21E-01 | 3.46E-04 | NA | 5.14E-04 |
| | 44000483 | *GOSR2* | 9.83E-01 | 9.38E-04 | NA | NA |
| 18 | 20140306 | *NPC1* | 4.62E-04 | NA | NA | NA |
| 19 | 4286174 | *PTPRS* | 6.68E-01 | 4.03E-03 | NA | NA |
| 19 | 7212650 | *FBN3* | 8.78E-03 | 4.21E-02 | NA | NA |
| 19 | 18049791 | *HOMER3* | 8.55E-03 | NA | NA | NA |
| 19 | 48199232 | *FUT2* | 9.13E-06 | NA | NA | NA |
| | 48401990 | *TULP2* | 2.78E-04 | NA | NA | 3.75E-04 |
| | 49342101 | *PLEKHA4* | NA | 6.10E-05 | NA | 1.05E-03 |
| | 57978423 | *ZNF324* | 3.52E-03 | NA | NA | NA |
| | 48339767 | *HSD17B14* | 3.68E-03 | NA | NA | NA |
| | 40223008 | *ITPKC* | 3.51E-03 | NA | NA | 6.55E-03 |
| | 48497828 | *ELSPBP1* | NA | NA | NA | 7.88E-03 |
| | 38747292 | *PPP1R14A* | NA | NA | NA | 9.36E-03 |
| | 48375649 | *PPP1R15A* | NA | NA | NA | 9.64E-03 |
| | 39897275 | *ZFP36* | NA | NA | NA | 9.76E-03 |
| 20 | 1 | *TRIB3* | 5.43E-03 | NA | NA | NA |
| 20 | 875154 | *SIRPA* | 3.68E-03 | NA | NA | NA |
| | 569278 | *SIRPB1* | 3.70E-03 | 2.58E-01 | NA | NA |
| 20 | 17446025 | *DZANK1* | 3.20E-03 | NA | NA | NA |
| | 31077893 | *CBFA2T2* | 3.85E-03 | NA | NA | NA |
| | 15554078 | *KIF16B* | 5.96E-02 | 2.01E-03 | NA | 2.78E-03 |
| | 23973360 | *APMAP* | 1.88E-04 | 5.50E-02 | NA | NA |
| | 17488199 | *SEC23B* | 2.15E-01 | 1.66E-03 | NA | 2.86E-01 |
| 20 | 47599536 | *SNAI1* | 3.69E-02 | NA | NA | 3.98E-03 |
| | 47552948 | *RNF114* | 1.20E-01 | 2.24E-02 | NA | 7.36E-03 |
| | 55136136 | *PCK1* | 5.82E-03 | 2.01E-01 | NA | NA |
| 21 | 33398153 | *OLIG2* | 6.73E-03 | NA | NA | NA |
| | 39816128 | *LCA5L* | 6.86E-03 | 6.94E-03 | NA | NA |
| | 39817781 | *SH3BGR* | 1.05E-01 | 7.71E-03 | NA | 8.08E-03 |
| 22 | 23951057 | *GUCD1* | 2.97E-04 | NA | NA | NA |
| | 19119330 | *ZDHHC8* | 1.32E-03 | NA | NA | NA |
| | 23863206 | *UPB1* | 6.83E-03 | 1.96E-03 | NA | NA |
| | 23666786 | *SPECC1L* | 4.95E-01 | 4.42E-03 | NA | NA |
| | 29234263 | *ASCC2* | 4.91E-01 | NA | NA | 9.68E-03 |
| | 40956767 | *CSDC2* | 4.17E-06 | NA | NA | NA |
| | 40940449 | *POLR3H* | 8.53E-06 | 4.77E-05 | NA | 4.35E-06 |
| | 40697526 | *ZC3H7B* | 1.67E-04 | 2.66E-05 | NA | NA |

| | | | | | | |
|---|---|---|---|---|---|---|
| 22 | 41095503 | *MEI1* | 1.49E-04 | NA | NA | NA |
| | 41017100 | *DESI1* | 4.62E-03 | NA | NA | NA |
| | 40487790 | *EP300* | 7.72E-03 | NA | NA | NA |
| | 40601209 | *L3MBTL2* | 1.45E-03 | 3.46E-02 | NA | NA |
| | 42090992 | *A4GALT* | 3.14E-01 | 8.26E-05 | NA | 7.43E-05 |
